# Barking up the microbiome tree: a multi-body-site characterization of maternal postpartum and pup early colonization dynamics

**DOI:** 10.64898/2026.08.04.742690

**Authors:** Tali Magory Cohen, Alisa Cohen, Sondra Turjeman, Sharon Kuzi, Smadar Tal, Omry Koren

**Affiliations:** Azrieli Faculty of Medicine, Bar-Ilan University, Safed, Israel; Hula Research Center, Tel-Hai, University of Kiryat Shmona in the Galilee, Israel; Koret School of Veterinary Medicine, The Hebrew University Veterinary Teaching Hospital, Hebrew University of Jerusalem, Rehovot, Israel

**Keywords:** Microbiome, Bacterial colonization, Maternal microbial transmission, Postpartum, Puppy microbiota, Canine model

## Abstract

Early-life microbial colonization is critical for shaping host development, yet how different maternal microbial reservoirs contribute to colonization of distinct offspring body sites remains poorly understood. Here, we explored microbial colonization and maturation of oral and rectal microbiota in puppies during the first postnatal week and assessed the contributions of maternal oral, vaginal, rectal, and milk microbiota to early colonization. We collected 505 samples from 33 dams and their litters at day 1 and day 8 postpartum and characterized microbial communities using 16S rRNA gene sequencing. We found that the pup oral microbiome on day 1 closely resembled maternal milk and vaginal microbiota. In contrast, the pup rectal microbiome was initially distinct from all maternal body sites, suggesting contributions from additional unmeasured sources. By day 8, both oral and rectal microbiomes exhibited signs of early maturation, with decreased relative abundance of opportunistic taxa and increased resemblance to maternal profiles. In dams, the vaginal microbiome showed the largest postpartum shift, while the rectal microbiome exhibited a smaller but significant change, and milk and oral microbiomes remained stable. Our results reveal that maternal contributions to early microbiome assembly are body-site specific, with oral and rectal microbiomes following distinct developmental trajectories during the first week of life, consistent with a dynamic colonization process shaped by both maternal and likely additional non-maternal sources. The observed patterns parallel those reported in humans, supporting the value of dogs as a comparative model for studying early-life microbiome colonization and maternal-offspring transmission in mammals.

## Introduction

Microbial colonization during early life plays a pivotal role in shaping host development and long-term health [1–4]. In particular, the establishment of the gut microbiome is critical for neonatal immune system maturation and metabolic function [5–7]. This early microbial community also provides a barrier against pathogens by competing for nutrients and modulating host responses [8, 9]. Disruptions to this natural colonization process (e.g., due to birth mode, antibiotic exposure, or environmental factors) can lead to adverse health outcomes, including increased susceptibility to infections, allergies, or metabolic disorders [10–12]. These effects have been well-documented in humans and are increasingly recognized in domestic species such as dogs [13].

Microbiome acquisition in early life occurs through multiple maternal and environmental pathways, and colonization patterns vary across host species and body sites. In humans, the process is well characterized: vaginal birth exposes neonates to maternal vaginal and gut microbiota [14, 15], while postnatal factors such as breastfeeding further shape the developing gut community [5, 16]. Breast milk, in particular, plays a dual role by mediating the transfer of nutrients and immunoglobulins and by supporting beneficial microbial colonization [5, 17]. Although less is known about these processes in dogs, evidence suggests that similar mechanisms are at play. Maternal transmission via the vaginal canal, fecal exposure, and milk or colostrum likely contribute to early microbial seeding [16, 18–21]. While the human gut microbiome reaches a stable, adult-like composition around three years of age, the fecal microbiota of dogs matures much earlier – typically by 7 to 9 weeks – making them a valuable model for studying early-life microbial development [22–24].

Although the importance of the microbiome in early development is well established, information on site-specific microbial dynamics in domestic dogs remains limited. Microbial niche specialization is well-documented in humans [25], but such distinctions may be less prominent in canines due to behaviors like self- and pup-grooming [13, 26]. Additionally, although maternal transmission is assumed to shape the early microbiome in puppies, the relative contributions of different microbial sources – such as the vaginal, rectal, oral, and milk microbiota – remain poorly understood. To address these gaps, we characterized the oral, vaginal, rectal, and milk microbiota of canine dams, compared these communities to the rectal and oral microbiota of their pups, and assessed patterns of microbial maturation in puppies during the early postnatal period. By mapping maternal-offspring microbial relationships across multiple body sites and timepoints (day 1 and day 8 postpartum), this study elucidates the timing, sources, and variability of early-life microbial colonization in dogs.

## Materials and methods

### Study design and sample collection

This study included purebred dogs from five kennels and two independent breeders located across northern and central Israel. We collected rectal and oral (buccal) swabs (Copan, Italy) from each dam and her pups on the day of birth and again one week later, timepoints chosen to capture the initial phase of neonatal microbiome development before stabilization and before weaning begins [24] (Table S1). We also collected dam vaginal swabs and milk samples at both timepoints. Vaginal swabs were collected from dams using sterile swabs (Puritan, USA). The swab was carefully inserted a few centimeters into the vaginal canal and rotated slightly upward to account for the anatomical angle of the tract. The procedure was brief and minimally invasive, consistent with the approved ethical sampling guidelines. Milk samples were collected directly from the dams’ teats into sterile 1.5 mL microcentrifuge tubes by a researcher wearing sterile gloves. No oxytocin or manual stimulation was applied to induce milk ejection; instead, milk was collected passively from available secretion at the time of sampling. Sample volumes varied among individuals (typically ≤0.7 mL), but equivalent aliquots were used downstream for DNA extraction. All samples were stored at −80 °C until processing. The collection protocol was approved by the Institutional Animal Care and Use Committee (IACUC) of the Hebrew University of Jerusalem, Ein Kerem campus (Approval # MD-21-16495-2). Birth mode was recorded by the attending veterinarian or breeder, and all included litters were confirmed to have been delivered vaginally.

### 16S rRNA gene sequencing and analysis

We extracted bacterial DNA from rectal, oral, and vaginal swabs using the PureLink Microbiome DNA Purification kit (Invitrogen, Thermo Fisher, Waltham, MA) according to the manufacturer’s instructions, following a 2-minute bead beating step. We subsequently amplified the V4 region of the 16S rRNA gene via PCR, using 515F-barcoded and 806R-non-barcoded primers [27]. Because bacterial load in milk samples is expected to be low [28], we extracted bacterial DNA from milk samples using the MasterPure Complete DNA & RNA Purification kit (Epicentre, Madison, WI) according to the manufacturer’s instructions, following centrifugation for the removal of the fat fraction, a subsequent two-minute bead beating step, and enzyme incubation with lysozyme for improved bacterial cell lysis. We amplified the extracted DNA using a two-step nested PCR approach. The first step employed 343F and 806R primers targeting the V3–V4 region of the 16S rRNA gene (5 cycles) [29, 30], followed by a second amplification using 515F and 806R primers, as described above.

We sequenced the samples on an Illumina MiSeq platform (Genomic Center, Azrieli Faculty of Medicine, Bar-Ilan University, Safed, Israel) using 2×250 bp paired-end chemistry for rectal, vaginal, and milk samples, and single-end 250 bp reads for oral samples. For the first sequencing batch (rectal, vaginal, and milk samples), the reverse reads showed low average quality scores and high truncation loss. Consequently, analyses were performed using only the forward reads. The subsequent batch (oral samples) was therefore sequenced in single-end mode to match analytical parameters. We included appropriate negative and positive controls.

We processed the 16S rRNA gene sequence data with QIIME2 version 2021.11 [31] using default parameters. We used DADA2 [32] to filter noisy sequences, correct sequencing errors, remove chimeras, and dereplicate reads into amplicon sequence variants (ASVs). We constructed a feature table from the denoised ASVs and assigned taxonomy using the *classify-sklearn* naïve Bayes classifier against the GreenGenes2 database [33].

After preprocessing with the QIIME2 pipeline, we performed downstream analysis using the R/bioconductor package ‘phyloseq’ (version 1.34.0), designed for analysis and visualization of high-throughput microbiome data [34]. Minimum read depth thresholds varied by sample group and comparison and are detailed in Table S2. Then, we normalized the samples either by rarefaction (using the ‘rarefy_even_depth’ function with a minimum sample size of 0.9 of the minimum read counts) or by scaling by relative abundance. Raw data and metadata are available online at https://www.ebi.ac.uk/ena/browser/view/PRJEB95990.

### Statistical analysis

We assessed alpha diversity with Faith’s Phylogenetic Diversity (PD) [35] using the twbattaglia/btools R package [36] and evaluated beta diversity using weighted UniFrac dissimilarity [37] in the R package ‘phyloseq’ [34], using the rarefied data set. We compared alpha diversity between dams and pups within the same timepoint and body site using matched dam-litter comparisons. For each litter, puppy alpha diversity was summarized as the median across pups and paired with the corresponding dam sample, and differences were tested using the Wilcoxon signed-rank test. Comparisons of paired samples across timepoints (day 1 vs. day 8) were likewise assessed using the Wilcoxon signed-rank test.

We visualized differences between groups via Principal Coordinates Analysis (PCoA) using the ‘plot_ordination’ function from the R package ‘phyloseq’. We tested group differences in community composition using permutational multivariate analysis of variance (PERMANOVA), implemented through the ‘adonis2’ function from the R package ‘vegan’ (v2.6-10) [38] with 9999 permutations. When only pups were analyzed, we included dam ID (mother) as a stratification variable to account for non-independence of samples from the same litter. When PERMANOVA revealed a significant effect for comparisons involving more than two groups, we tested pairwise differences in microbial community composition using the ‘pairwise.adonis2’ function from the R package ‘pairwiseAdonis’, applying weighted UniFrac distances, 9999 permutations, and false discovery rate (FDR, Benjamini-Hochberg) correction for multiple comparisons [39]. We assessed the homogeneity of group dispersions using the ‘betadisper’ function from the R package ‘vegan’ to ensure the validity of PERMANOVA assumptions.

To estimate maternal contributions to early puppy microbiome assembly, we applied FEAST (Fast Expectation-Maximization for microbial Source Tracking) [40]. FEAST estimates the proportional contribution of each source while accounting for an additional “unknown source” (i.e., taxa that cannot be attributed to the sampled maternal body sites) component representing unobserved environmental inputs. Because maternal oral samples were processed separately from the other maternal source types, source-tracking models that included oral samples were highly sensitive to batch structure and were not considered reliable for biological inference. We therefore restricted the primary FEAST analysis to a non-oral model in which maternal milk, vaginal, and rectal microbiota were treated as candidate sources and puppy rectal samples as sinks. FEAST was run separately for day 1 and day 8 postpartum, using only source samples from the corresponding dam for each puppy, and source proportions were normalized within each run to sum to 1. As a sensitivity analysis, we repeated the non-oral model in a leave-one-source-out framework to assess redistribution of attribution among the remaining measured sources and the unknown component.

To identify differentially abundant taxa across body sites, we applied MaAsLin2 (Multivariable Association Testing, version 1.18.0) [41] to the non-rarefied, filtered dataset. We specified a minimum prevalence threshold of 0.1 (i.e., present in ≥10% of samples) and a minimum relative abundance of 0.001 (i.e., ≥0.1%). Each model included a single fixed effect representing the grouping variable of interest (e.g., host type, sampling day, or body site) and random effects were incorporated where appropriate to account for repeated measures (e.g., maternal identity). To analyze taxa-specific differences in relative abundances across groups, we performed Kruskal-Wallis tests followed by Dunn’s post-hoc comparisons on the differentially abundant taxa and visualized the detected variation across taxa using the ‘ggplot2’ R package (version 3.5.1) [42].

To predict the functional potential of the bacterial communities, we applied PICRUSt2 (version 2) [43] with default parameters. The average Nearest Sequenced Taxon Index (NSTI) score was 0.159±0.327, suggesting moderate reference genome coverage. We then used the R package ‘ggpicrust2’ [44] to visualize functional predictions based on Enzyme Commission (EC) numbers. From these predictions, we generated error bar plots (‘pathway_errorbar’ function, R package ‘ggpicrust2’) and principal component analysis (PCA) plots (‘prcomp’ function, R package ‘stats’ [45]) using the unstratified output from PICRUSt2, with KEGG pathways as features. We tested group differences in community composition using PERMANOVA, implemented through the ‘adonis2’ function from the R package ‘vegan’ (v2.6-10) on Bray-Curtis dissimilarities with 9999 permutations. To assess the similarity between groups in functional profiles, we calculated Euclidean distances between group centroids in PCA space, derived from PICRUSt2-predicted functional profiles by averaging the coordinates of samples within each group.

## Results

### Sampling and sequencing success

We collected 505 samples from 33 litters comprising 33 dams and 210 puppies originating from 5 kennels and 2 breeders and representing 13 dog breeds (Table S1). The median litter size was 6 puppies (range 2-11), and the median dam age was 4 years (range 2-6). Seven puppies died before the second sampling on day 8 and were therefore omitted from the analysis. In addition, 75 samples were omitted due to low sequencing depth. A total of 195 dam samples and 235 pup samples, collected at two timepoints (day 1 and day 8 postpartum) from four body sites in dams and two in pups, were available for analysis (Table S1).

### Microbiome diversity of dams

To assess postpartum changes in the dam microbiome, we compared microbial communities between samples collected on day 1 and day 8. We found that alpha diversity increased significantly between collection timepoints in vaginal samples (*P* < 0.001, Figure 1A, Table S3). Additionally, microbial beta diversity was significantly different between samples collected at different timepoints in vaginal (F = 5.07, R2 = 0.11, P < 0.001) and rectal samples (F = 2.76, R2 = 0.04, P = 0.023), but not in milk or oral samples (Table S4, Figure 1B-E). Homogeneity of dispersion was similar across timepoints in all sampled body sites (Table S5).

**Figure 1.**
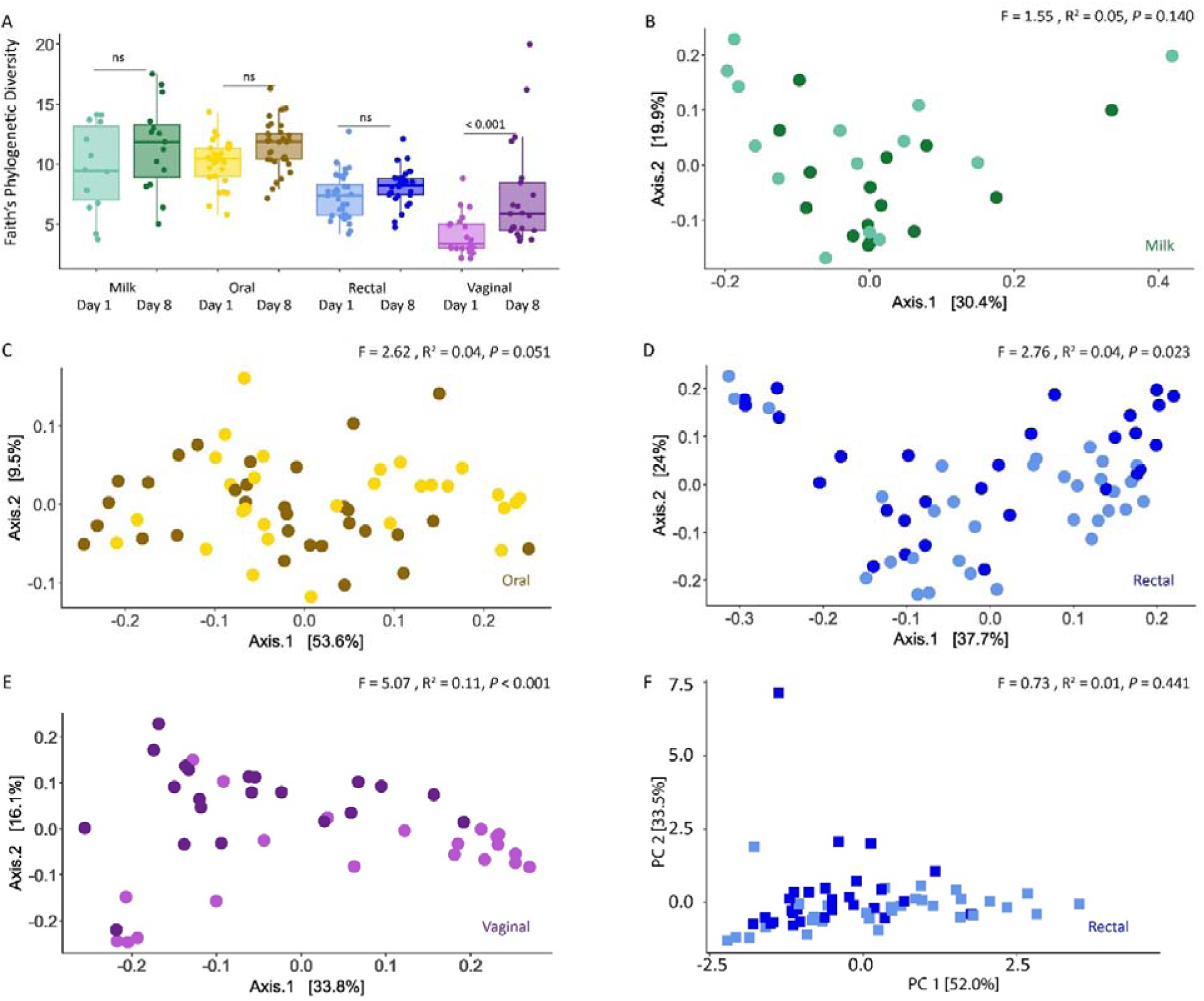
Microbiome diversity of dams in four body sites (milk, oral, rectal, and vaginal). (A) Faith’s phylogenetic diversity. Alpha diversity differed significantly between timepoints in vaginal samples (Wilcoxon signed-rank test; Table S3). (B-E) Principal Coordinate Analysis (PCoA) based on weighted UniFrac distances of (B) milk (green), (C) oral (yellow), (D) rectal (blue), and (E) vaginal (purple) samples. Day 1 are shown in l ighter shades and day 8 in darker shades throughout. Community composition differed significantly over time in rectal (P = 0.023) and vaginal (P < 0.001) samples, while oral (P = 0.051) and milk (P = 0.140) samples showed no significant change. (F) Principal Component Analysis (PCA) based on differentially abundant predicted functional pathways (adjusted P < 0.05) of rectal samples from day 1 (l ighter shades) and day 8 (darker shades).

To better understand how the microbial community of dams changes in response to parturition, we identified differentially abundant taxa as a function of time postpartum using MaAsLin2. Differentially abundant taxa were only identified in milk and rectal samples, but not in oral and vaginal samples (Table S6). Only one taxon (species *Ruminococcus B gnavus*) was found to be differentially abundant between day 1 and day 8 in milk samples, and two taxa (the species *Blautia A 141780* argi and the genus *Faecalibacterium*) in rectal samples (Adjusted P < 0.05, Table S6).

In addition, we examined predicted functional and pathway-level differences between day 1 and day 8 across dam body sites. No significant changes were observed in the milk, oral, or vaginal sites, whereas four features were differentially abundant in rectal samples (Table S7). Of these, three were more prevalent on day 1 (linked to the renin-angiotensin system and the RIG-I-like receptor signaling pathway), whereas a single feature associated with vasopressin-regulated water reabsorption was more abundant on day 8. These results suggest that some functional capabilities of the rectal microbiome shift during the early postpartum period. Functional differences were reflected in partial spatial separation of rectal samples from day 1 and day 8 (Figure 1F).

### Microbiome diversity ofpups

We compared pup oral and rectal microbial communities between samples collected on day 1 and day 8 to examine early-life microbial colonization and maturation. Alpha diversity increased significantly between collection timepoints in both oral and rectal samples (Figure 2A, Table S3). Similarly, microbial beta diversity was significantly different between samples collected at different timepoints in both oral (F = 21.71, R^2^ = 0.16, *P* < 0.001, Figure 2B) and rectal samples (F = 17.14, R^2^ = 0.14, *P* < 0.001, Table S4, Figure 2C). Homogeneity of dispersion differed between sampling timepoints in oral samples but not in rectal samples (Table S5), indicating that observed differences in oral beta diversity reflect both shifts in community composition and changes in within-group variability.

**Figure 2.**
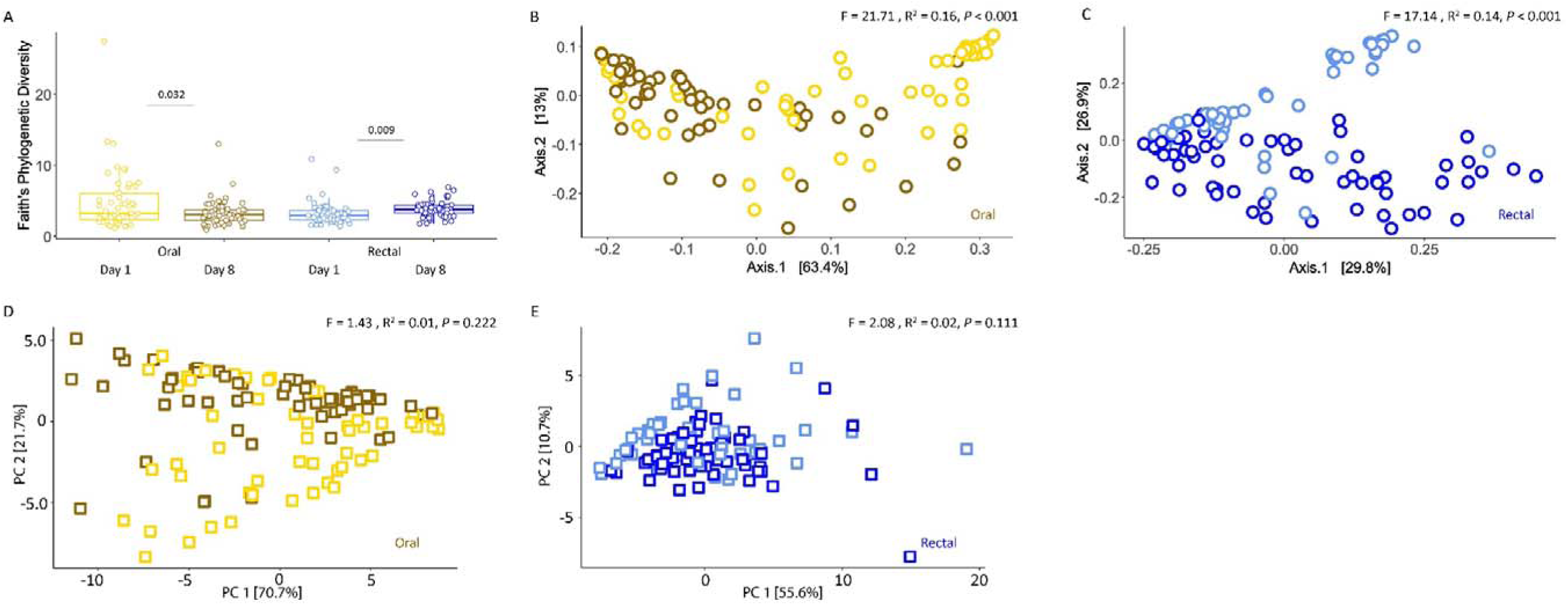
Microbiome diversity and functionality of pups in two body sites (oral and rectal). (A) Faith’s phylogenetic diversity. Alpha diversity differed significantly between timepoints in rectal samples (Wilcoxon signed-rank test, Table S3). (B, C) Principal Coordinate Analysis (PCoA) based on weighted Uni Frac distances of oral (yellow) and rectal (blue) samples, respectively, from day 1 (lighter shades) and day 8 (darker shades). Community composition differed significantly over time in both oral and rectal samples (P < 0.001). (D, E) Principal Component Analysis (PCA) based on differentially abundant predicted functional pathways (adjusted P < 0.05) of oral (yellow, panel D) and rectal (blue, panel E) samples, respectively, from day 1 (l ighter shades) and day 8 (darker shades).

Additionally, we identified differentially abundant taxa in pups over the first 8 days of life using MaAsLin2. A total of 34 taxa (1 order, 1 class, 4 families, 16 genera and 12 species) and 58 taxa (6 families, 28 genera and 24 species) were differentially abundant between day 1 and day 8 in oral and rectal samples (Adjusted P < 0.05, Table S6), respectively. In oral samples collected on day 1, 18 out of the 21 day 1-enriched taxa belonged to the phyla Pseudomonadota, Bacillota I, and Actinomycetota. In contrast, taxa enriched on day 8 were primarily from Bacillota I and Bacillota A 368345 (4 taxa from each, out of 13 total day 8-enriched taxa). In rectal samples, day 1-enriched taxa were mainly from Bacillota A 368345, Pseudomonadota, and Bacillota I (24 out of 27), while those more abundant on day 8 were mostly from Bacillota A 368345, Bacillota I, and Fusobacteriota (25 out of 31).

In addition, functional differences between day 1 and day 8 identified 192 differentially abundant features in oral samples and 136 in rectal samples (Table S7). Annotation of these features revealed distinct functional profiles between timepoints (Figure S1). In oral samples, functions enriched on day 1 were predominantly associated with metabolic pathways such as tyrosine metabolism, glyoxylate and dicarboxylate metabolism, carbon fixation, and toluene degradation. In contrast, functions enriched on day 8 included metabolic pathways (e.g., alanine, aspartate and glutamate metabolism; amino sugar and nucleotide sugar metabolism; sphingolipid metabolism; nitrogen metabolism), as well as pathways related to organismal systems (e.g., antigen processing and presentation, NOD-like receptor signaling) and cellular processes (e.g., lysosome) (Figure S1A). In rectal samples, day 1-enriched functions were primarily linked to carbohydrate metabolism (e.g., fructose and mannose metabolism, galactose metabolism, and pentose and glucuronate interconversions), glycan biosynthesis and metabolism (other glycan degradation, glycosphingolipid biosynthesis – globo and isoglobo series, glycosaminoglycan degradation, and glycosphingolipid biosynthesis – ganglio series), biosynthesis of other secondary metabolites (streptomycin biosynthesis) and cell growth and death (Figure S1B). By day 8, functions more abundant in rectal samples were related to amino acid metabolism (e.g., lysine degradation, tryptophan metabolism), lipid metabolism (e.g., glycerophospholipid metabolism, fatty acid degradation), xenobiotic biodegradation and metabolism (benzoate metabolism), and genetic information processing (replication and repair – base excision repair). Functional differences were sufficient to drive spatial separation between oral samples from day 1 and day 8 in PCA space, whereas no clear separation was detected for rectal samples despite their functional differences (Figure 2D, E).

### Comparison ofdams and pup microbiomes

To evaluate similarities between maternal and pup microbiomes, we compared microbial communities between dam and pup samples at each timepoint separately. Pairwise distances between group centroids indicated that pup oral microbiomes were most similar to dam vaginal and milk microbiomes at both timepoints (Table S8). PERMANOVA analyses confirmed significant differences among groups (Table S4, Figure 3A, B). Homogeneity of dispersion varied across body sites at both timepoints (Table S5), which should be considered when interpreting group-level comparisons.

**Figure 3.**
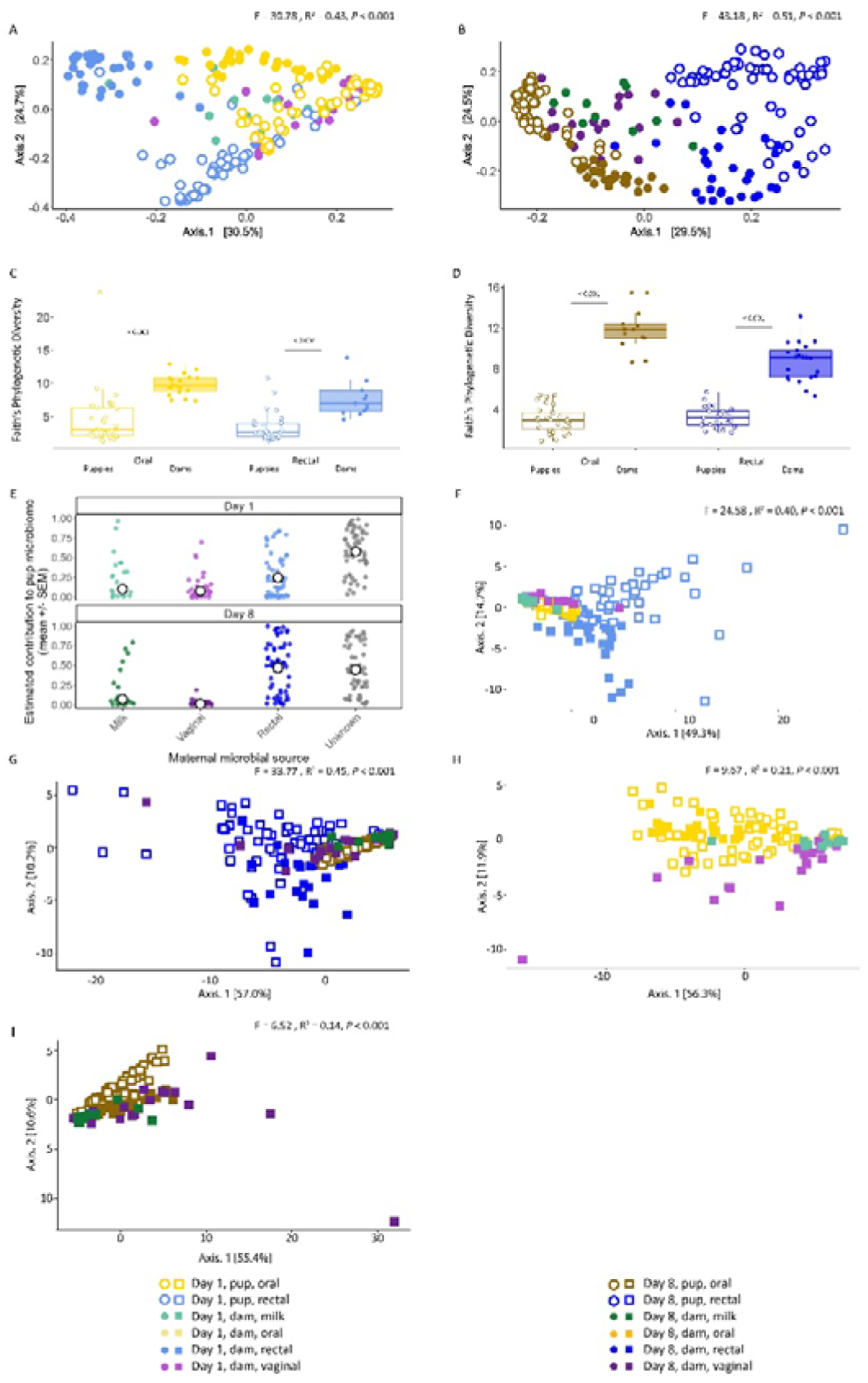
Comparison of microbiome diversity between dams and pups. (A, B) Principal Coordinate Analysis (PCoA) based on weighted UniFrac distances, showing microbial communities in milk (green), oral (yellow), rectal (blue), and vaginal (purple) samples collected on day 1 (A, lighter shades) and day 8 (B, darker shades). Samples from dams are represented by filled circles, and samples from pups by open circles. Pup oral samples clustered most closely with milk and vaginal samples from dams on day 1, whereas pup rectal samples remained distinct from maternal body sites in community composition (Table S8). (C, D) Faith’s phylogenetic diversity. Differences between oral (yellow) and rectal (blue) samples between dams (filled) and pups (open) were statistically significant in both body sites in both timepoints (Wilcoxon signed-rank test; Table S3). Alpha diversity was consistently higher in dams’ samples. (E) Source tracking analysis of maternal contributions to puppy rectal microbiomes using FEAST. Colored points represent individual puppy samples, and open circles with black error bars show mean estimated source contributions ± SEM for maternal milk, rectal, and vaginal microbiota, together with an unknown component, on day 1 (top) and day 8 (bottom). Contributions assigned to “unknown” represent taxa not attributable to the sampled maternal body sites. (F, G) Principal Component Analysis (PCA) based on differentially abundant predicted functional pathways (adjusted P < 0.05) of milk (green), oral (yellow), rectal (blue) and vaginal (purple) samples. Day 1 samples are shown in lighter shades (F) and day 8 in darker shades(G). At both timepoints, oral samples from pups clustered most closely with those from dams, and rectal samples from pups clustered most closely with dam rectal samples (Table S12). (H, I) PCA based on differentially abundant predicted functional pathways (adjusted P < 0.05) of milk (green), oral (yellow) and vaginal (purple) samples, excluding rectal samples to allow better resolution across these three body sites. Day 1 samples are shown in lighter shades (H) and day 8 in darker shades (I).

To better characterize potential maternal sources of pup microbiomes, we applied FEAST source tracking using maternal milk, rectal, and vaginal microbiota as candidate sources. A subset of pups was excluded from the FEAST analysis because either the puppy rectal sink sample or all corresponding maternal source samples did not pass the predefined read-depth filters. Using the non-oral FEAST model, we found that puppy rectal microbiomes could be traced to several maternal non-oral sources, although a substantial fraction remained unattributed. On day 1, maternal rectal microbiota made the strongest traced contribution (25%), followed by milk (10%) and then vaginal microbiota (8%), whereas by day 8 maternal rectal microbiota became even more prominent as the principal traced source (47%), with milk contributing modestly (7%) and vaginal contribution remaining low (1%). Across both timepoints, a substantial proportion of the pup rectal microbiome remained unattributed (57% on day 1, 45% on day 8), suggesting contributions from unmeasured sources.

In addition, we compared alpha diversity between oral and rectal samples of dams and pups at each timepoint. We observed significantly greater alpha diversity in dam oral and rectal microbiomes than in those of pups both at postpartum day 1 and day 8 (adjusted P < 0.001, Figure 3C, D, Table S3). The oral microbiome of pups was distinct from that of dams and was dominated by the genera Frederiksenia, Staphylococcus, and Canicola (Table 1). In contrast, the oral microbial communities of dams included the genera Porphyromonas A, Conchiformibius, and Moraxella C 651924 as well as Frederiksenia. Rectal microbial communities in pups showed greater variability between timepoints than oral communities, with the most abundant taxa including Escherichia, Clostridium T, Fusobacterium A, and Bacteroides H 857956. In dams, the rectal microbiome was characterized by Fusobacterium A and B, Phocaeicola A, Alloprevotella, Bacteroides H 857956, Peptacetobacter, Corynebacterium, and Escherichia.

**Table 1.**
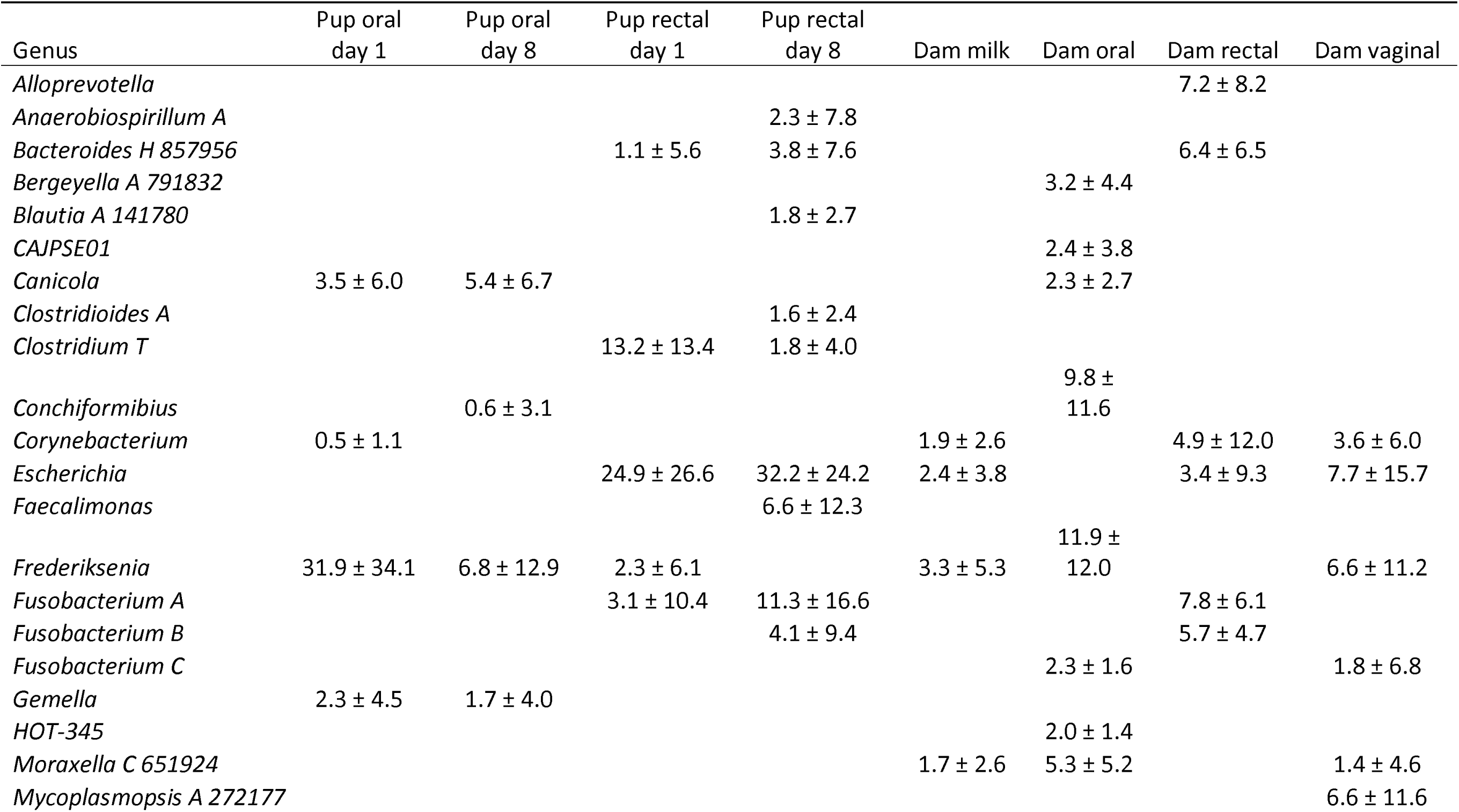
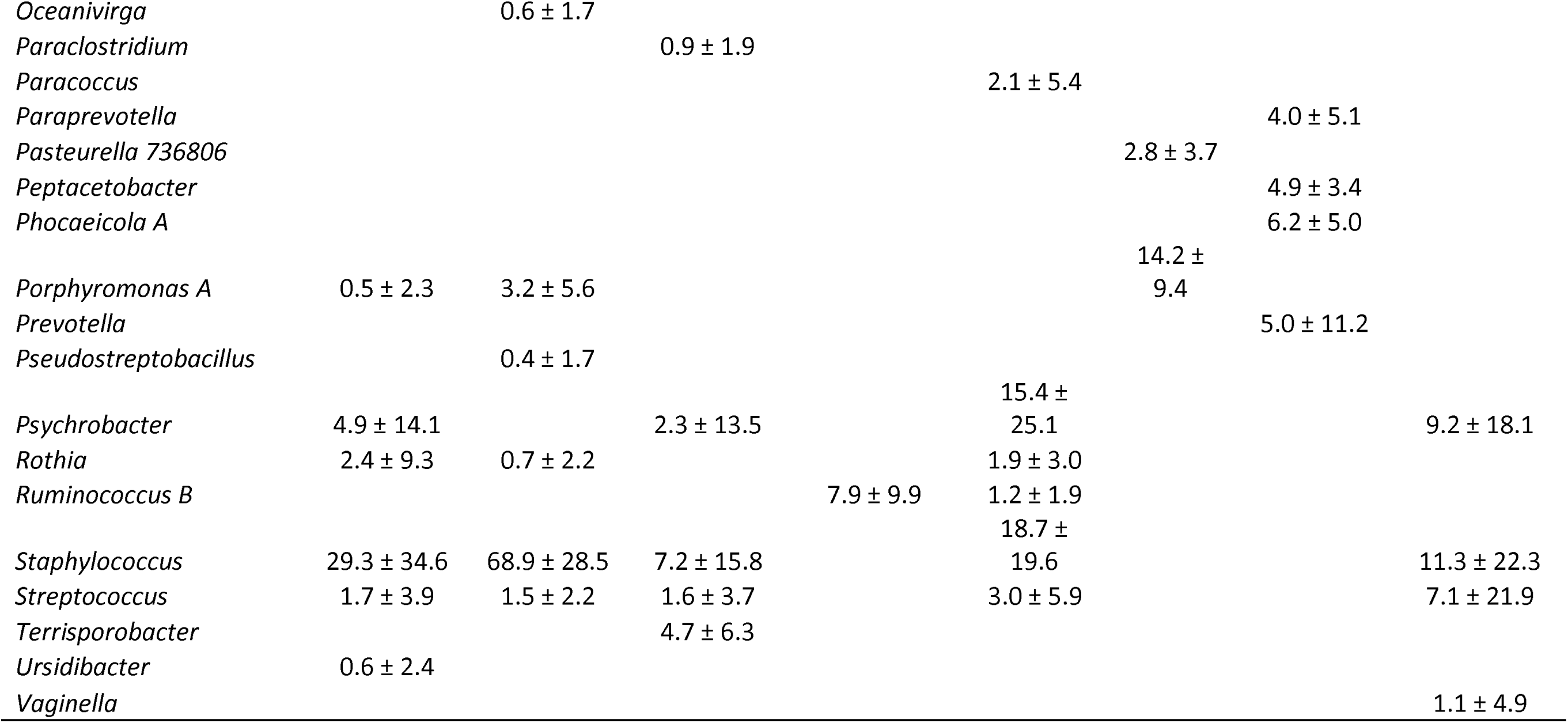
Top 10 most abundant genera in each microbial community of pups and dams, respectively. Values represent relative abundances (%) ± standard deviation. For pups, values are shown separately for day 1 and day 8 postpartum, while for dams, values are across both timepoints as previous analyses reveal uniformity in the rectal microbiome across postpartum timepoints. Taxonomic assignments follow the GreenGenes2 database and correspond to genus-level classifications derived from 16S rRNA V4 amplicon sequences.

Differential abundance analysis revealed many taxa that differed significantly between dams and pups in both oral and rectal samples at each timepoint. In oral samples, 108 taxa (2 orders, 1 class, 17 families, 39 genera, 48 species and one unclassified taxon assigned to the domain Bacteria) were differentially abundant on day 1, and 113 taxa (2 orders, 1 class, 18 families, 35 genera and 57 species) on day 8 (Adjusted P < 0.05, Table S9). Rectal samples showed 109 differentially abundant taxa on day 1 (15 families, 31 genera, and 63 species) and 105 on day 8 (12 families, 33 genera and 60 species). Across both body sites, the number of taxa more abundant in pups relative to dams decreased over time, reflecting a convergence in community composition consistent with ongoing microbial colonization in pups. Taxa from both body sites more abundant in dams belonged to the phyla Bacillota A 368345, Bacteroidota, and Pseudomonadota at both timepoints, with Bacillota I also enriched in dam rectal samples.

We further explored microbiome differences by comparing pup and dam oral and rectal samples at both postpartum timepoints (four groups per body site) to identify patterns of convergence over time. In total, 120 taxa in oral samples and 135 in rectal samples were differentially abundant across the four groups (Table S10). Notably, several taxa that were significantly more abundant in pups on day 1, including taxa often associated with neonatal opportunistic colonization, decreased in relative abundance by day 8 and were consistently low in dams at both timepoints (Figure 4).

**Figure 4.**
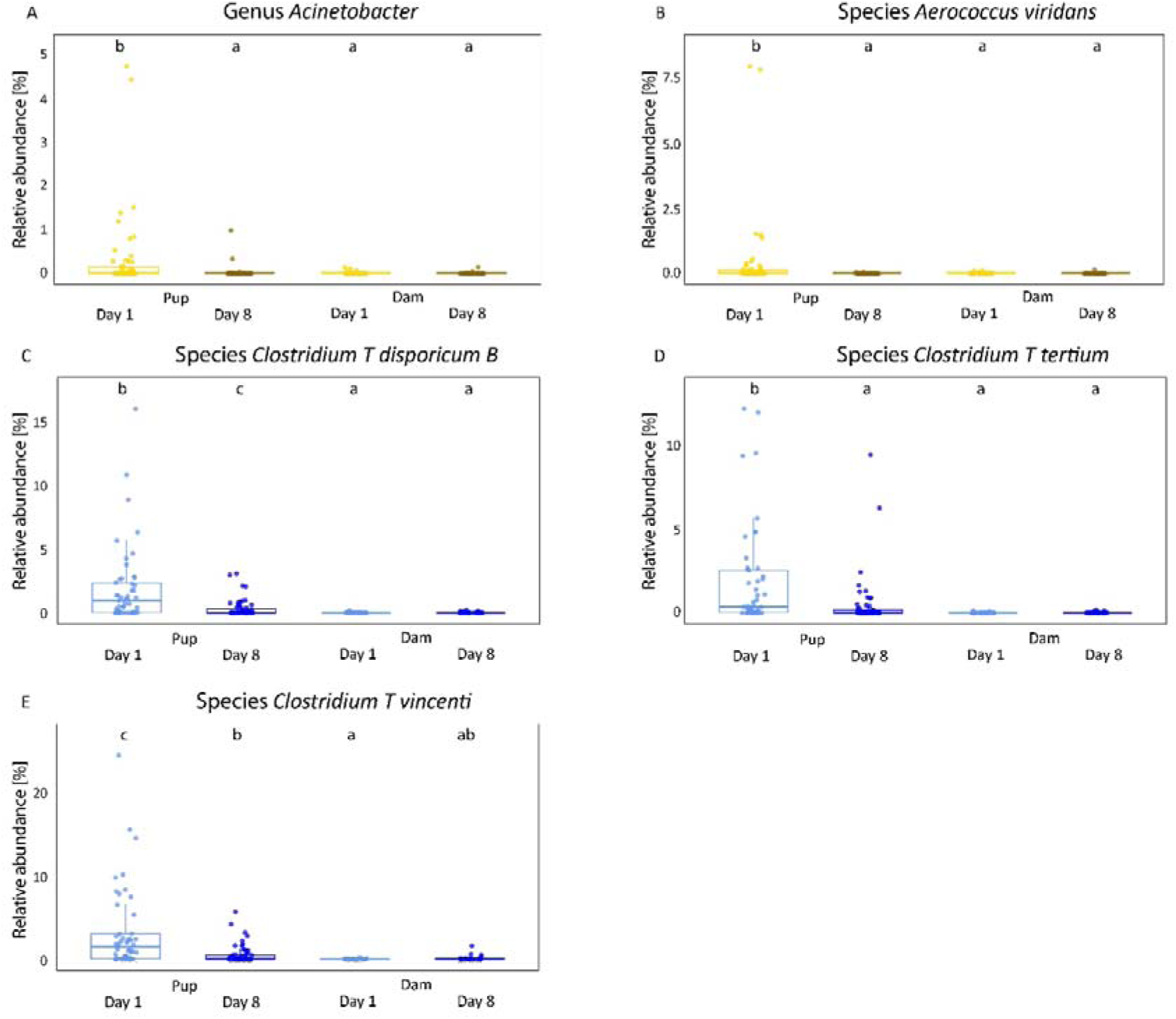
Bacterial taxa with significantly higher relative abundance in pups on day 1 postpartum compared to pups on day 8 and to dams at both timepoints (adjusted P < 0.05) in oral and rectal sites. Differentially abundant taxa were first identified using MaAsLin2. Differences between groups were further assessed with Kruskal–Wallis tests followed by Dunn’s post hoc comparisons. Different lowercase letters above the boxes indicate significant pairwise differences. Species shown here differed significantly between pups on day 1 and at least 2 other groups and are suspected or known pathogens. Additional significantly differentiated taxa and corresponding post hoc results are provided in Table S10.

Functional profiling revealed significant differences between dams and pups, with 156 differentially abundant features in oral samples and 179 in rectal samples on day 1, and 218 in oral and 214 in rectal samples on day 8 (Table S11). These functional distinctions reflected the spatial separation observed in microbial composition (Figure 3A, B). Notably, pup oral microbiomes clustered closely not only with those of dam milk and vaginal samples, but also with dam oral microbiome at both timepoints (Table S12, Figure 3F, G). In contrast, rectal microbiomes remained distinct between dams and pups and were separated from all other body sites. Therefore, we proceeded to visualize only milk, oral, and vaginal samples to better resolve spatial separation in predicted functional profiles across these microbiomes. When rectal samples were excluded, pup oral microbiome predicted functionality overlapped with that of all three microbiota communities (Figure 3H, I).

## Discussion

In this study, we explored microbial colonization and maturation in puppies during the early postnatal period, focusing on the role of the maternal microbiome in contributing to these processes. By comparing microbial profiles between dams and pups across different body sites, we found that the pup oral microbiome initially resembled the dam’s milk and vaginal microbiomes in overall community composition, and included several opportunistic pathogens, while the rectal microbiome showed a more distinct composition early after birth. Source-tracking in the non-oral model indicated that puppy rectal microbiomes received measurable contributions from multiple maternal sources, with both maternal rectal and milk microbiota contributing substantially on day 1, and maternal rectal microbiota emerging as the largest contributor by day 8. The leave-one-source-out sensitivity analysis further supported this pattern, indicating that rectal source attribution remained consistently important even when other measured maternal sources were excluded. Over time, both oral and rectal microbial communities in pups stabilized and became more similar to those of the dams, with the relative abundance of opportunistic taxa decreasing toward maternal profiles. Overall, maternal sites were relatively stable over days 1-8, with the largest change in the vaginal microbiome and a smaller but significant rectal shift. These dynamics are consistent with patterns observed in other mammals, including humans [46, 47], positioning this species as a promising model for studying microbiome colonization and maturation in mammals.

### Differential maternal contributions to oral and rectal microbiome development in pups

The development of the microbiome in puppies during early life is a dynamic process influenced by many factors, of which vertical transmission is thought to be particularly important [22]. Here, we show that from as early as one day postpartum, distinct microbial communities colonize the oral and rectal body sites in puppies. Although the pup oral microbiome has been linked to postnatal contact with the birthing dam and birth mode, it is distinct from that of adult dogs [48], and to our knowledge, has not been directly compared to microbial communities of dams (but see [20]). Our findings suggest that the pup oral microbiome shares substantial similarity with the dam’s milk and vaginal microbiota. However, because oral samples were processed separately and oral FEAST models were not considered reliable, these observations reflect community similarity rather than direct source attribution. Functional predictions suggested broadly similar metabolic potential across these sites, with the pup oral microbiome showing the closest similarity to the maternal oral functional profile. This pattern is consistent with evidence from humans indicating that colostrum and milk serve as major sources for early microbial colonization of the infant oral cavity [47, 49, 50], alongside contributions from the maternal vaginal microbiome [51, 52]. The high abundance of *Staphylococcus* and *Streptococcus* in pup oral, dam milk and vaginal samples is consistent with previous findings in pups [48] and in dam milk and vaginal microbiota [20, 28, 53]. While this pattern is consistent with maternal microbial inputs contributing to neonatal oral microbiome development, it does not directly demonstrate such transmission. Although the long-term consequences of oral microbiome dysbiosis on infant and adult health remain unclear, it is plausible that disruptions to early colonizers, that play a critical role in shaping the environment for beneficial anaerobic commensals, may impair nutrient absorption and trigger systemic inflammation, potentially impacting immunocompetence, development, and neurological health [54]. Because oral samples were sequenced in a separate run, a potential batch effect cannot be fully excluded. However, the clustering of pup oral samples with dam milk and vaginal communities, rather than forming a distinct run-specific group, suggests that any batch effect did not override the underlying biological signal.

Despite these apparent maternal associations in the oral microbiome, the rectal microbial communities of pups on the first day postpartum were notably distinct from all maternal body sites, including the dam’s own rectal microbiome, though some overlap with milk and vaginal profiles was observed. Source-tracking analyses nevertheless indicated measurable contributions from multiple maternal microbial sources, with the maternal rectal microbiome representing the largest identifiable contributor, particularly by day 8, although a substantial proportion remained unattributed. The presence of genera such as *Staphylococcus*, *Streptococcus*, *Escherichia*, and *Psychrobacter* in both pups and dams supports previous findings in pup meconium [20, 53] and is compatible with maternal contribution despite early compositional differences. As in other mammals, early neonatal gut communities may initially include opportunistic or environmentally derived bacteria before stabilizing over time [21, 55]. In dogs, the whelping process, during which pups are born within the amniotic sac and the dam may need to tear it open with her teeth, may further increase environmental microbial exposure immediately postpartum. Consistent with findings in pigs, where a large proportion of neonatal gut bacteria originate from maternal milk, skin, vagina, and feces [56], our results point to multisite maternal contributions to pup gut colonization. In humans, gut microbiota development is likewise shaped by maternal vaginal and fecal microbiota and heavily influenced by delivery mode [15], with cesarean-born infants showing increased similarity to maternal skin [51]. Consistent with these patterns, our findings underscore that maternal microbial contribution represents an important, though not exclusive, component of early gut microbiota colonization in pups. Together, these findings suggest that early gut colonization in pups reflects a combination of maternal microbial inputs and rapid postnatal microbial turnover.

### Early-life microbial maturation in pups

The neonatal microbiome undergoes rapid development, eventually converging toward an adult-like state. However, the pace and trajectory of this maturation are influenced by host species, diet, and life history. In mammals, this period is marked by rapid colonization and high microbial strain turnover compared to adults [57]. In canine pups, previous studies have documented significant shifts in microbial communities during pre-weaning development [21, 24], likely driven by the establishment of early colonizers. Consistent with these findings, our results support dynamic postpartum microbial maturation, with taxa often associated with early-life instability, particularly Clostridium species, present at higher abundance on day 1 but declining by day 8 toward maternal levels [21, 58]. The early presence of opportunistic taxa is a hallmark of microbiome instability immediately after birth and commonly occurs alongside early maternal and environmental microbial inputs [55, 59], and may reflect the oxygenated state of the neonatal gastrointestinal tract, which favors initial colonization by facultative anaerobes [60]. Depletion of oxygen and a reduction in redox potential promote taxa turnover in favor of obligate anaerobes, leading to a rapid change in microbiome composition together with milk ingestion [24, 61]. Notably, we show that this maturation trend extends beyond the gut to include the oral microbiome, where factors such as milk exposure, littermate interactions, and environmental contact likely play key roles. In parallel to these compositional changes, microbial diversity also increased over time. Consistent with this, microbial diversity remained significantly lower in pups across body sites at both postpartum timepoints, suggesting maturation toward adult-like communities was still ongoing at the time of sampling. Evidence indicates that gut microbial diversity in dogs stabilizes around three months of age and then remains relatively stable until 12 years of age [24]. Future longitudinal studies will be essential to fully resolve the timeline of this convergence and its underlying drivers. Although some studies have reported comparable microbial profiles between pups and dams within 24 hours of birth [58], our findings align with research showing that microbial diversity and community composition in pups remain distinct from adults until at least 60 days postpartum [20, 21]. Overall, these observations suggest that weaning is a critical milestone in microbial convergence in both canines [20, 22] and humans [62, 63] as well as in other mammals [64, 65], though exceptions have been noted [66, 67]. Understanding the timing and drivers of this microbial maturation may provide new insights into early-life health trajectories and interventions across species.

### Dynamics of postpartum microbial changes in dams

The periparturient period (immediately before, during, and after birth) is characterized by substantial shifts in the maternal microbiome, influenced by a complex interplay of physiological factors such as changes in hormone levels, immune responses, and metabolic adjustments [9, 68–70]. These changes, however, vary by body site. While the vaginal microbiome is known to shift rapidly postpartum, microbial alterations in milk, oral, and rectal communities are either observed later or remain relatively stable throughout lactation [46, 68, 71, 72]. Specifically, in humans, postpartum vaginal communities typically shift from a *Lactobacillus*-predominant state to a *Lactobacillus*-depleted state [73, 74]. Although *Lactobacillus* species were not specifically identified in antepartum and postpartum comparisons in canines using culture-based methods, differences in other taxa were detected [75], along with a reduction in microbial heterogeneity over time [20]. Direct, sequencing-based assessments of canine postpartum vaginal microbiota composition remain limited. Consistent with patterns observed in humans, our data show significant changes in the dam’s vaginal microbiome after birth, both in diversity and composition, despite no notable shifts in *Lactobacillus* levels. We also observed a small but significant difference in rectal microbiota between postpartum timepoints, accompanied by minor predicted functional differences. These subtle changes may reflect the short interval between sampling points, during which key hormonal effects might not yet be fully manifested. Maternal reproductive hormones have been implicated in postpartum microbial dynamics, including the role of declining estrogen levels in the postpartum loss of vaginal *Lactobacillus* predominance, and the role of progesterone in shaping rectal microbiome composition during lactation [76]. Future research would benefit from the simultaneous monitoring of reproductive hormone levels and microbial community composition across multiple body sites and timepoints throughout pregnancy and lactation.

This study characterizes early-life microbial colonization and maturation in canines, revealing distinct developmental trajectories between oral and rectal microbiomes and the importance of maternal contributions to early microbiome assembly. By capturing temporal and body site-specific dynamics in both pups and dams, we show that early colonization involves differential maternal inputs followed by rapid microbial turnover and convergence toward adult-like communities. These patterns closely parallel those observed in humans and other mammals, highlighting the value of dogs as a model for studying maternal microbial contribution and early-life microbiome maturation.

## Declarations

### Ethics approval and consent to participate

All animal procedures were conducted under approval from the Institutional Animal Care and Use Committee (IACUC) of the Hebrew University of Jerusalem, Ein Kerem campus (Approval #MD-21-16495-2). The study adhered to the Hebrew University’s institutional guidelines for animal care and use, the regulatory framework governing animal research at our institution.

### Consent for publication

Not applicable

### Availability of data and material

The dataset generated and analyzed during the current study is available in the European Bioinformatics Institute (EBI) – European Nucleotide Archive (ENA) data repository at https://www.ebi.ac.uk/ena/browser/view/PRJEB95990, and in the QIITA repository, https://qiita.ucsd.edu/public/?study_id=16099.

## Competing interests

The authors declare that they have no competing interests.

## Funding

TMC was supported by the Planning and Budgeting committee (PBC) of the Council for Higher Education (CHE) initiative for Non-Faculty Researchers.

## Authors’ contributions

AC, SmT and OK conceptualized the idea for the study; TMC, AC and SoT conceived and executed the experiments and subsequent analyses; OK supervised; TMC, AC, SoT and OK wrote the original draft; TMC, AC, SoT, SK, SmT, and OK edited the draft.

## Acknowledgements

TMC would like to thank Sol and Charlie Magory for their fruitful discussion.

## AI Use Declaration

This manuscript was prepared with the assistance of generative artificial intelligence tools. Specifically, ChatGPT (OpenAI, GPT-5.3) and OpenAI Codex (0.118.0) were used for language editing and code troubleshooting only.

---

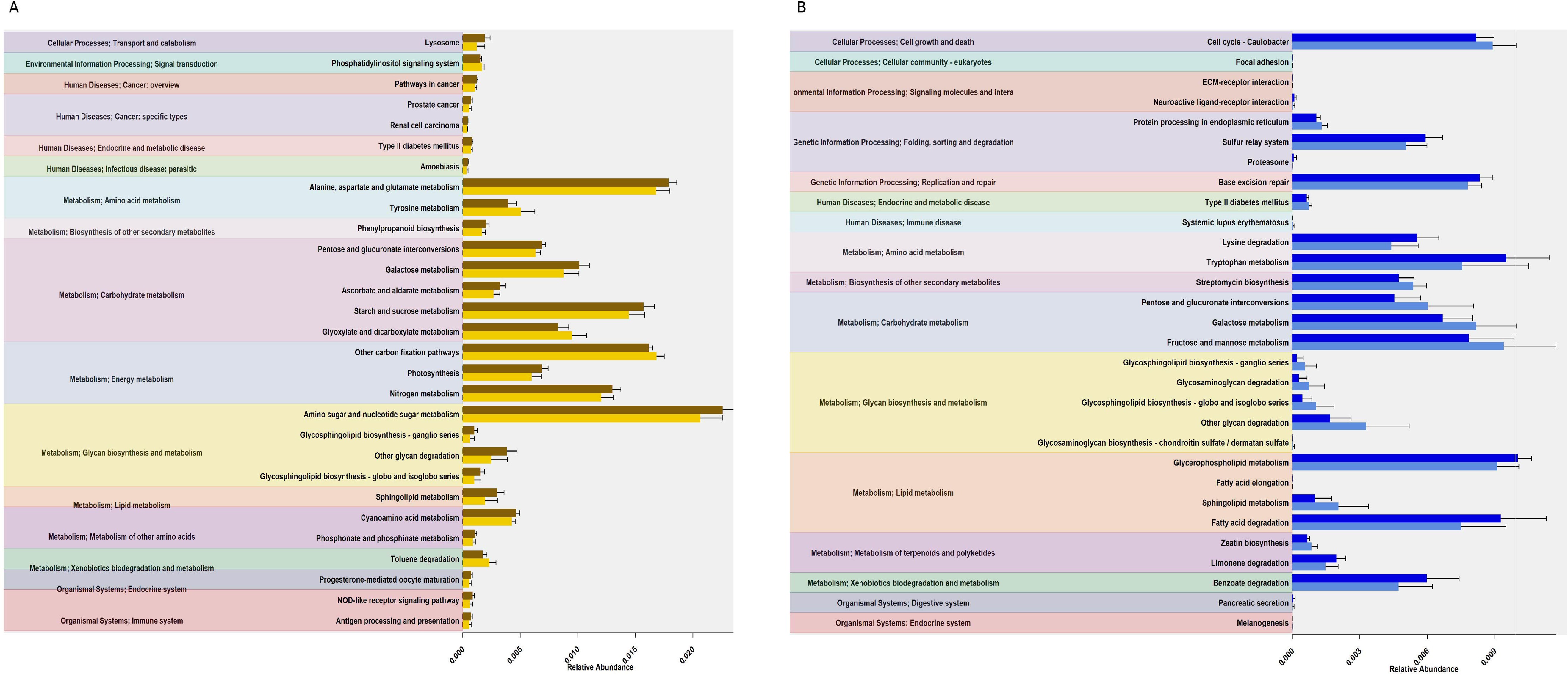

## Notes

https://www.ebi.ac.uk/ena/browser/view/PRJEB95990

https://qiita.ucsd.edu/public/?study_id=16099

### Competing Interest Statement

The authors have declared no competing interest.

