## Supplementary material for "Barking up the microbiome tree: a multi-body-site characterization of maternal postpartum and pup early colonization dynamics": Code: feast_refactor.R

```
library(dplyr)
library(tidyr)
library(ggplot2)
library(openxlsx)
library(forcats)
library(FEAST)

feast_colors <- c(
  "Dam_Milk_Day1" = "#66C2A5",
  "Dam_Milk_Day8" = "#117733",
  "Dam_Rectal_Day1" = "cornflowerblue",
  "Dam_Rectal_Day8" = "blue2",
  "Dam_Vaginal_Day1" = "mediumorchid",
  "Dam_Vaginal_Day8" = "darkorchid4",
  "Dam_Oral_Day1" = "gold1",
  "Dam_Oral_Day8" = "darkgoldenrod4",
  "Unknown" = "grey50"
)

normalize_feast_runs <- function(feast_df) {
  if (is.null(feast_df) || nrow(feast_df) == 0) {
    return(feast_df)
  }

  feast_df %>%
    group_by(Day, PupSample, PupSite, DropSource) %>%
    mutate(
      run_total = sum(Prop, na.rm = TRUE),
      Prop = dplyr::if_else(run_total > 0, Prop / run_total, NA_real_)
    ) %>%
    ungroup() %>%
    select(-run_total)
}

annotate_feast_results <- function(feast_df) {
  if (is.null(feast_df) || nrow(feast_df) == 0) {
    return(feast_df)
  }

  feast_df %>%
    mutate(
      PupID = sub("_(Rectal|Oral|Milk|Vaginal)$", "", PupSample),
      SourceSite = case_when(
        SourceEnv == "Rectal" | grepl("Rectal", SourceEnv) ~ "Rectal",
        SourceEnv == "Vaginal" | grepl("Vaginal", SourceEnv) ~ "Vaginal",
        SourceEnv == "Milk" | grepl("Milk", SourceEnv) ~ "Milk",
        SourceEnv == "Oral" | grepl("Oral", SourceEnv) ~ "Oral",
        SourceEnv == "Unknown" ~ "Unknown",
        TRUE ~ "Other"
      ),
      SourceDay = case_when(
        Day == "Day 1" & SourceSite == "Milk" ~ "Dam_Milk_Day1",
        Day == "Day 8" & SourceSite == "Milk" ~ "Dam_Milk_Day8",
        Day == "Day 1" & SourceSite == "Rectal" ~ "Dam_Rectal_Day1",
        Day == "Day 8" & SourceSite == "Rectal" ~ "Dam_Rectal_Day8",
        Day == "Day 1" & SourceSite == "Vaginal" ~ "Dam_Vaginal_Day1",
        Day == "Day 8" & SourceSite == "Vaginal" ~ "Dam_Vaginal_Day8",
        Day == "Day 1" & SourceSite == "Oral" ~ "Dam_Oral_Day1",
        Day == "Day 8" & SourceSite == "Oral" ~ "Dam_Oral_Day8",
        SourceSite == "Unknown" ~ "Unknown",
        TRUE ~ "Other"
      ),
      SourceDay = factor(SourceDay, levels = c(
        "Dam_Milk_Day1", "Dam_Milk_Day8",
        "Dam_Rectal_Day1", "Dam_Rectal_Day8",
        "Dam_Vaginal_Day1", "Dam_Vaginal_Day8",
        "Dam_Oral_Day1", "Dam_Oral_Day8",
        "Unknown"
      ))
    )
}

add_feast_batch_column <- function(meta_df) {
  meta_df %>%
    mutate(
      batch = case_when(
        Type == "Oral" ~ "2",
        Type %in% c("Rectal", "Vaginal", "Milk") ~ "1",
        TRUE ~ NA_character_
      )
    )
}

detect_batch_column <- function(meta_df, batch_column_candidates) {
  matches <- intersect(batch_column_candidates, colnames(meta_df))
  if (length(matches) == 0) {
    stop(
      paste0(
        "Batch correction requested, but none of these metadata columns were found: ",
        paste(batch_column_candidates, collapse = ", "),
        ". Available columns are: ",
        paste(colnames(meta_df), collapse = ", ")
      )
    )
  }
  matches[1]
}

prepare_feast_counts <- function(count_matrix, target_depth = NULL) {
  count_matrix <- as.matrix(count_matrix)
  count_matrix[is.na(count_matrix)] <- 0
  count_matrix[count_matrix < 0] <- 0

  if (is.null(target_depth)) {
    target_depth <- round(rowSums(count_matrix))
  }

  row_totals <- rowSums(count_matrix)
  nonzero_rows <- row_totals > 0
  count_matrix_rel <- matrix(0, nrow = nrow(count_matrix), ncol = ncol(count_matrix))
  rownames(count_matrix_rel) <- rownames(count_matrix)
  colnames(count_matrix_rel) <- colnames(count_matrix)

  if (any(nonzero_rows)) {
    count_matrix_rel[nonzero_rows, ] <- sweep(
      count_matrix[nonzero_rows, , drop = FALSE],
      1,
      row_totals[nonzero_rows],
      "/"
    )
  }

  target_depth <- pmax(round(target_depth), 1)
  count_matrix_scaled <- sweep(count_matrix_rel, 1, target_depth, "*")
  count_matrix_int <- round(count_matrix_scaled)
  storage.mode(count_matrix_int) <- "integer"

  keep_taxa <- colSums(count_matrix_int, na.rm = TRUE) > 0
  count_matrix_int[, keep_taxa, drop = FALSE]
}

apply_batch_correction_to_counts <- function(count_matrix, batch_info) {
  if (!requireNamespace("limma", quietly = TRUE)) {
    stop("Please install the limma package to perform batch correction.")
  }

  count_matrix <- as.matrix(count_matrix)
  lib_sizes <- pmax(rowSums(count_matrix), 1)
  rel_mat <- sweep(count_matrix, 1, lib_sizes, "/")
  log_mat <- log2(rel_mat * 1e6 + 1)
  corrected_log <- t(limma::removeBatchEffect(t(log_mat), batch = batch_info))
  corrected_log[corrected_log < 0] <- 0

  corrected_abund <- pmax(2^corrected_log - 1, 0)
  corrected_totals <- rowSums(corrected_abund)
  corrected_rel <- matrix(0, nrow = nrow(corrected_abund), ncol = ncol(corrected_abund))
  rownames(corrected_rel) <- rownames(corrected_abund)
  colnames(corrected_rel) <- colnames(corrected_abund)

  keep_rows <- corrected_totals > 0
  if (any(keep_rows)) {
    corrected_rel[keep_rows, ] <- sweep(
      corrected_abund[keep_rows, , drop = FALSE],
      1,
      corrected_totals[keep_rows],
      "/"
    )
  }

  sweep(corrected_rel, 1, lib_sizes, "*")
}

read_feast_output <- function(prop_file, pup_sample, dam_id, pup_site, day_label, drop_source) {
  if (!file.exists(prop_file)) {
    return(NULL)
  }

  prop_df <- read.table(
    prop_file,
    header = TRUE,
    row.names = 1,
    check.names = FALSE
  )

  if (nrow(prop_df) == 0) {
    return(NULL)
  }

  prop_df %>%
    pivot_longer(everything(), names_to = "SourceEnv", values_to = "Prop") %>%
    mutate(
      PupSample = pup_sample,
      DamID = dam_id,
      PupSite = pup_site,
      Day = day_label,
      DropSource = drop_source
    )
}

run_feast_pipeline_day <- function(
  phy_obj_day,
  day_label,
  analysis_label,
  sink_sites,
  source_sites,
  batch_correction = FALSE,
  batch_column_candidates = c("Batch", "batch", "Plate")
) {
  otu_mat <- as.data.frame(t(otu_table(phy_obj_day)))
  meta_df <- data.frame(sample_data(phy_obj_day)) %>%
    mutate(sampleID = paste(Host_ID, Type, sep = "_")) %>%
    filter(Host_new %in% c("Dam", "Pup")) %>%
    mutate(Type = as.character(Type))

  if (batch_correction) {
    meta_df <- add_feast_batch_column(meta_df)
  }

  rownames(otu_mat) <- meta_df$sampleID

  batch_column <- NULL
  if (batch_correction) {
    batch_column <- detect_batch_column(meta_df, batch_column_candidates)
  }

  pups_subset <- meta_df %>%
    filter(Host_new == "Pup", Type %in% sink_sites)

  results_list <- list()
  temp_dir <- file.path(tempdir(), paste0("feast_", analysis_label))
  dir.create(temp_dir, showWarnings = FALSE, recursive = TRUE)

  for (i in seq_len(nrow(pups_subset))) {
    pup_row <- pups_subset[i, ]
    pup_sample <- pup_row$sampleID
    dam_id <- pup_row$Mother
    pup_site <- pup_row$Type

    source_samples <- meta_df %>%
      filter(Host_new == "Dam", Host_ID == dam_id, Type %in% source_sites)

    if (nrow(source_samples) == 0) {
      next
    }

    run_one_model <- function(current_sources, drop_source = "None") {
      samples_use <- intersect(c(pup_sample, current_sources$sampleID), rownames(otu_mat))
      if (length(samples_use) < 2) {
        return(NULL)
      }

      C <- otu_mat[samples_use, , drop = FALSE]
      metadata_feast <- meta_df %>%
        filter(sampleID %in% samples_use) %>%
        mutate(
          Env = ifelse(Host_new == "Pup", "Unknown", Type),
          SourceSink = ifelse(Host_new == "Pup", "Sink", "Source"),
          id = 1
        ) %>%
        select(sampleID, Env, SourceSink, id)

      common_samples <- intersect(rownames(C), metadata_feast$sampleID)
      if (length(common_samples) < 2) {
        return(NULL)
      }

      C <- as.matrix(C[common_samples, , drop = FALSE])
      metadata_feast <- metadata_feast %>%
        filter(sampleID %in% common_samples) %>%
        arrange(match(sampleID, common_samples))
      rownames(metadata_feast) <- metadata_feast$sampleID

      original_depth <- rowSums(C)

      if (batch_correction) {
        batch_info <- meta_df %>%
          filter(sampleID %in% common_samples) %>%
          arrange(match(sampleID, common_samples)) %>%
          pull(all_of(batch_column))

        if (any(is.na(batch_info))) {
          stop("Batch column contains NA values for FEAST samples. Please clean the metadata first.")
        }

        if (length(unique(batch_info)) > 1) {
          C <- apply_batch_correction_to_counts(C, batch_info)
        }
      }

      C_feast <- prepare_feast_counts(C, target_depth = original_depth)
      if (ncol(C_feast) < 2 || any(rowSums(C_feast) == 0)) {
        return(NULL)
      }

      metadata_feast_input <- metadata_feast %>% select(Env, SourceSink, id)
      rownames(metadata_feast_input) <- metadata_feast$sampleID

      outfile <- paste0(
        analysis_label, "_", gsub(" ", "", day_label), "_",
        gsub("[^A-Za-z0-9]", "_", pup_sample), "_", drop_source
      )

      FEAST(
        C = C_feast,
        metadata = metadata_feast_input,
        dir_path = temp_dir,
        different_sources_flag = 1,
        outfile = outfile
      )

      read_feast_output(
        file.path(temp_dir, paste0(outfile, "_source_contributions_matrix.txt")),
        pup_sample = pup_sample,
        dam_id = dam_id,
        pup_site = pup_site,
        day_label = day_label,
        drop_source = drop_source
      )
    }

    results_list[[length(results_list) + 1]] <- run_one_model(source_samples, "None")

    for (drop_source in source_sites) {
      source_samples_drop <- source_samples %>% filter(Type != drop_source)
      if (nrow(source_samples_drop) == 0) {
        next
      }
      results_list[[length(results_list) + 1]] <- run_one_model(source_samples_drop, drop_source)
    }
  }

  bind_rows(results_list)
}

summarize_feast_analysis <- function(feast_results, source_sites) {
  feast_results <- feast_results %>%
    normalize_feast_runs() %>%
    annotate_feast_results()

  expected_sites <- c(source_sites, "Unknown")

  raw_main <- feast_results %>% filter(DropSource == "None")
  raw_drop <- feast_results %>% filter(DropSource != "None")

  per_pup <- raw_main %>%
    group_by(Day, PupID, PupSite, SourceSite) %>%
    summarise(mean_prop = mean(Prop, na.rm = TRUE), .groups = "drop") %>%
    complete(nesting(Day, PupID, PupSite), SourceSite = expected_sites, fill = list(mean_prop = 0))

  population <- per_pup %>%
    group_by(Day, PupSite, SourceSite) %>%
    summarise(
      mean_contribution = mean(mean_prop, na.rm = TRUE),
      median_contribution = median(mean_prop, na.rm = TRUE),
      sd_contribution = sd(mean_prop, na.rm = TRUE),
      n_pups = n(),
      .groups = "drop"
    )

  summary_by_source <- per_pup %>%
    group_by(Day, PupSite, SourceSite) %>%
    summarise(
      mean_prop = mean(mean_prop, na.rm = TRUE),
      median_prop = median(mean_prop, na.rm = TRUE),
      n_pups = n(),
      .groups = "drop"
    ) %>%
    mutate(SourceEnv = SourceSite) %>%
    select(Day, PupSite, SourceEnv, SourceSite, mean_prop, median_prop, n_pups)

  drop_per_pup <- raw_drop %>%
    group_by(Day, PupID, PupSite, SourceSite, DropSource) %>%
    summarise(mean_prop = mean(Prop, na.rm = TRUE), .groups = "drop") %>%
    complete(nesting(Day, PupID, PupSite, DropSource), SourceSite = expected_sites, fill = list(mean_prop = 0))

  drop_population <- drop_per_pup %>%
    group_by(Day, PupSite, SourceSite, DropSource) %>%
    summarise(
      mean_contribution = mean(mean_prop, na.rm = TRUE),
      median_contribution = median(mean_prop, na.rm = TRUE),
      sd_contribution = sd(mean_prop, na.rm = TRUE),
      n_pups = n(),
      .groups = "drop"
    )

  list(
    all = feast_results,
    raw_main = raw_main,
    raw_drop = raw_drop,
    per_pup = per_pup,
    population = population,
    summary_by_source = summary_by_source,
    drop_per_pup = drop_per_pup,
    drop_population = drop_population
  )
}

build_sum_diagnostic <- function(df, group_cols, value_col) {
  if (is.null(df) || nrow(df) == 0) {
    return(tibble(note = "No rows available for this diagnostic."))
  }

  df %>%
    group_by(across(all_of(group_cols))) %>%
    summarise(
      total_prop = sum(.data[[value_col]], na.rm = TRUE),
      n_rows = n(),
      off_by = abs(total_prop - 1),
      .groups = "drop"
    ) %>%
    arrange(desc(off_by))
}

add_diagnostic_sheet <- function(wb, sheet_name, df, group_cols, value_col) {
  addWorksheet(wb, sheet_name)
  writeData(wb, sheet_name, build_sum_diagnostic(df, group_cols, value_col))
}

save_feast_stacked_bar <- function(df, file_name, title_text) {
  plot_df <- annotate_feast_results(df)
  plot_df$PupID <- factor(plot_df$PupID, levels = sort(unique(plot_df$PupID)))
  plot_df$FillKey <- factor(
    as.character(plot_df$SourceDay),
    levels = c(
      "Dam_Milk_Day1", "Dam_Milk_Day8",
      "Dam_Vaginal_Day1", "Dam_Vaginal_Day8",
      "Dam_Rectal_Day1", "Dam_Rectal_Day8",
      "Dam_Oral_Day1", "Dam_Oral_Day8",
      "Unknown"
    )
  )

  fill_values <- c(
    "Dam_Milk_Day1" = unname(feast_colors["Dam_Milk_Day1"]),
    "Dam_Milk_Day8" = unname(feast_colors["Dam_Milk_Day8"]),
    "Dam_Vaginal_Day1" = unname(feast_colors["Dam_Vaginal_Day1"]),
    "Dam_Vaginal_Day8" = unname(feast_colors["Dam_Vaginal_Day8"]),
    "Dam_Rectal_Day1" = unname(feast_colors["Dam_Rectal_Day1"]),
    "Dam_Rectal_Day8" = unname(feast_colors["Dam_Rectal_Day8"]),
    "Dam_Oral_Day1" = unname(feast_colors["Dam_Oral_Day1"]),
    "Dam_Oral_Day8" = unname(feast_colors["Dam_Oral_Day8"]),
    "Unknown" = unname(feast_colors["Unknown"])
  )

  fill_labels <- c(
    "Dam_Milk_Day1" = "Milk (Day 1)",
    "Dam_Milk_Day8" = "Milk (Day 8)",
    "Dam_Vaginal_Day1" = "Vaginal (Day 1)",
    "Dam_Vaginal_Day8" = "Vaginal (Day 8)",
    "Dam_Rectal_Day1" = "Rectal (Day 1)",
    "Dam_Rectal_Day8" = "Rectal (Day 8)",
    "Dam_Oral_Day1" = "Oral (Day 1)",
    "Dam_Oral_Day8" = "Oral (Day 8)",
    "Unknown" = "Unknown"
  )

  p <- plot_df %>%
    ggplot(aes(x = PupID, y = Prop, fill = FillKey)) +
    geom_col(position = "stack") +
    facet_grid(Day ~ PupSite, scales = "free_x", space = "free_x") +
    labs(
      x = "",
      y = "Estimated source proportion",
      fill = "Source",
      title = title_text
    ) +
    scale_fill_manual(
      values = fill_values,
      labels = fill_labels,
      drop = TRUE,
      na.value = "grey50"
    ) +
    theme_bw() +
    theme(
      axis.text.x = element_blank(),
      axis.ticks.x = element_blank(),
      plot.title = element_text(hjust = 0.5)
    )

  dir.create(dirname(file_name), recursive = TRUE, showWarnings = FALSE)
  ggsave(file_name, p, width = 10, height = 6, dpi = 300)
}

save_feast_drop_one_plot <- function(df, file_name, title_text) {
  if (nrow(df) == 0) {
    return(invisible(NULL))
  }

  plot_df <- annotate_feast_results(df) %>%
    filter(SourceEnv != "Unknown") %>%
    mutate(DayPup = factor(paste(Day, PupSite, sep = " ")))

  p <- plot_df %>%
    ggplot(aes(x = SourceSite, y = Prop, fill = SourceDay, color = SourceDay)) +
    geom_boxplot(outlier.shape = NA, alpha = 0.6) +
    geom_jitter(width = 0.15, size = 2.5, alpha = 0.7) +
    facet_grid(DayPup ~ DropSource, scales = "free_y") +
    labs(
      x = "Maternal microbial source",
      y = "Estimated contribution to pup microbiome",
      fill = "Source",
      color = "Source",
      title = title_text
    ) +
    scale_fill_manual(values = feast_colors, na.value = "grey50") +
    scale_color_manual(values = feast_colors, na.value = "grey30") +
    theme_classic() +
    theme(
      panel.border = element_rect(color = "black", fill = NA, linewidth = 0.8),
      axis.text.x = element_text(angle = 45, hjust = 1, size = 10),
      strip.text = element_text(size = 11),
      plot.title = element_text(size = 14, hjust = 0.5),
      legend.position = "right"
    )

  dir.create(dirname(file_name), recursive = TRUE, showWarnings = FALSE)
  ggsave(file_name, p, width = 14, height = 10, dpi = 300)
}

save_feast_boxplot_main <- function(df, file_name, title_text) {
  if (nrow(df) == 0) {
    return(invisible(NULL))
  }

  plot_df <- annotate_feast_results(df) %>%
    mutate(
      SourceSite = factor(SourceSite, levels = c("Milk", "Vaginal", "Rectal", "Oral", "Unknown")),
      Day = factor(Day, levels = c("Day 1", "Day 8"))
    )

  summary_df <- plot_df %>%
    select(Day, PupID, PupSite, SourceSite, Prop) %>%
    complete(
      nesting(Day, PupID, PupSite),
      SourceSite = c("Milk", "Vaginal", "Rectal", "Oral", "Unknown"),
      fill = list(Prop = 0)
    ) %>%
    filter(SourceSite %in% unique(as.character(plot_df$SourceSite))) %>%
    mutate(
      SourceDay = case_when(
        Day == "Day 1" & SourceSite == "Milk" ~ "Dam_Milk_Day1",
        Day == "Day 8" & SourceSite == "Milk" ~ "Dam_Milk_Day8",
        Day == "Day 1" & SourceSite == "Vaginal" ~ "Dam_Vaginal_Day1",
        Day == "Day 8" & SourceSite == "Vaginal" ~ "Dam_Vaginal_Day8",
        Day == "Day 1" & SourceSite == "Rectal" ~ "Dam_Rectal_Day1",
        Day == "Day 8" & SourceSite == "Rectal" ~ "Dam_Rectal_Day8",
        Day == "Day 1" & SourceSite == "Oral" ~ "Dam_Oral_Day1",
        Day == "Day 8" & SourceSite == "Oral" ~ "Dam_Oral_Day8",
        SourceSite == "Unknown" ~ "Unknown",
        TRUE ~ NA_character_
      )
    ) %>%
    group_by(Day, SourceSite, SourceDay) %>%
    summarise(
      mean_prop = mean(Prop, na.rm = TRUE),
      sd_prop = sd(Prop, na.rm = TRUE),
      n = sum(!is.na(Prop)),
      sem_prop = ifelse(n > 1, sd_prop / sqrt(n), NA_real_),
      .groups = "drop"
    )

  p <- plot_df %>%
    ggplot(aes(x = SourceSite, y = Prop, color = SourceDay)) +
    geom_jitter(width = 0.15, size = 2.2, alpha = 0.75) +
    geom_errorbar(
      data = summary_df,
      aes(
        x = SourceSite,
        ymin = pmax(mean_prop - sem_prop, 0),
        ymax = pmin(mean_prop + sem_prop, 1)
      ),
      width = 0.12,
      linewidth = 1.2,
      color = "black",
      inherit.aes = FALSE
    ) +
    geom_point(
      data = summary_df,
      aes(x = SourceSite, y = mean_prop),
      shape = 21,
      size = 5,
      stroke = 1.2,
      fill = "white",
      color = "black",
      inherit.aes = FALSE
    ) +
    facet_wrap(~ Day, ncol = 1) +
    labs(
      x = "Maternal microbial source",
      y = "Estimated contribution to pup microbiome\n(mean +/- SEM)",
      fill = NULL,
      color = NULL,
      title = NULL
    ) +
    scale_fill_manual(values = feast_colors, labels = c(
      "Dam_Milk_Day1" = "Milk (Day 1)",
      "Dam_Milk_Day8" = "Milk (Day 8)",
      "Dam_Vaginal_Day1" = "Vaginal (Day 1)",
      "Dam_Vaginal_Day8" = "Vaginal (Day 8)",
      "Dam_Rectal_Day1" = "Rectal (Day 1)",
      "Dam_Rectal_Day8" = "Rectal (Day 8)",
      "Dam_Oral_Day1" = "Oral (Day 1)",
      "Dam_Oral_Day8" = "Oral (Day 8)",
      "Unknown" = "Unknown"
    ), drop = TRUE, na.value = "grey50") +
    scale_color_manual(values = feast_colors, labels = c(
      "Dam_Milk_Day1" = "Milk (Day 1)",
      "Dam_Milk_Day8" = "Milk (Day 8)",
      "Dam_Vaginal_Day1" = "Vaginal (Day 1)",
      "Dam_Vaginal_Day8" = "Vaginal (Day 8)",
      "Dam_Rectal_Day1" = "Rectal (Day 1)",
      "Dam_Rectal_Day8" = "Rectal (Day 8)",
      "Dam_Oral_Day1" = "Oral (Day 1)",
      "Dam_Oral_Day8" = "Oral (Day 8)",
      "Unknown" = "Unknown"
    ), drop = TRUE, na.value = "grey30") +
    theme_classic() +
    theme(
      panel.border = element_rect(color = "black", fill = NA, linewidth = 0.8),
      axis.text.x = element_text(angle = 45, hjust = 1, size = 14),
      axis.text.y = element_text(size = 14),
      axis.title.x = element_text(size = 16),
      axis.title.y = element_text(size = 16),
      strip.text = element_text(size = 15),
      legend.position = "none"
    )

  dir.create(dirname(file_name), recursive = TRUE, showWarnings = FALSE)
  ggsave(file_name, p, width = 7, height = 5, dpi = 300)
}

run_feast_analysis <- function(
  phyloseq_by_day,
  analysis_label,
  sink_sites,
  source_sites,
  batch_correction = FALSE,
  batch_column_candidates = c("Batch", "batch", "Plate")
) {
  raw_results <- bind_rows(lapply(
    names(phyloseq_by_day),
    function(day_label) {
      run_feast_pipeline_day(
        phy_obj_day = phyloseq_by_day[[day_label]],
        day_label = day_label,
        analysis_label = analysis_label,
        sink_sites = sink_sites,
        source_sites = source_sites,
        batch_correction = batch_correction,
        batch_column_candidates = batch_column_candidates
      )
    }
  ))

  summarize_feast_analysis(raw_results, source_sites = source_sites)
}

export_feast_analysis <- function(
  analysis,
  workbook_path,
  main_plot_path,
  main_plot_title,
  drop_plot_path,
  drop_plot_title,
  drop_plot_filter_sites,
  box_plot_path = NULL,
  box_plot_title = NULL
) {
  wb <- createWorkbook()
  addWorksheet(wb, "raw_main")
  writeData(wb, "raw_main", as.data.frame(analysis$raw_main))
  addWorksheet(wb, "per_pup")
  writeData(wb, "per_pup", as.data.frame(analysis$per_pup))
  addWorksheet(wb, "population")
  writeData(wb, "population", as.data.frame(analysis$population))
  addWorksheet(wb, "summary_by_source")
  writeData(wb, "summary_by_source", as.data.frame(analysis$summary_by_source))

  add_diagnostic_sheet(wb, "diag_raw_main", analysis$raw_main, c("Day", "PupSample", "PupSite"), "Prop")
  add_diagnostic_sheet(wb, "diag_per_pup", analysis$per_pup, c("Day", "PupID", "PupSite"), "mean_prop")
  add_diagnostic_sheet(wb, "diag_population", analysis$population, c("Day", "PupSite"), "mean_contribution")

  if (nrow(analysis$raw_drop) > 0) {
    addWorksheet(wb, "drop_one_raw")
    writeData(wb, "drop_one_raw", as.data.frame(analysis$raw_drop))
    addWorksheet(wb, "drop_one_per_pup")
    writeData(wb, "drop_one_per_pup", as.data.frame(analysis$drop_per_pup))
    addWorksheet(wb, "drop_one_population")
    writeData(wb, "drop_one_population", as.data.frame(analysis$drop_population))

    add_diagnostic_sheet(wb, "diag_drop_one_raw", analysis$raw_drop, c("Day", "PupSample", "PupSite", "DropSource"), "Prop")
    add_diagnostic_sheet(wb, "diag_drop_one_per_pup", analysis$drop_per_pup, c("Day", "PupID", "PupSite", "DropSource"), "mean_prop")
    add_diagnostic_sheet(wb, "diag_drop_one_pop", analysis$drop_population, c("Day", "PupSite", "DropSource"), "mean_contribution")
  }

  saveWorkbook(wb, workbook_path, overwrite = TRUE)

  save_feast_stacked_bar(analysis$raw_main, main_plot_path, main_plot_title)
  if (!is.null(box_plot_path) && !is.null(box_plot_title)) {
    save_feast_boxplot_main(analysis$raw_main, box_plot_path, box_plot_title)
  }
  save_feast_drop_one_plot(
    analysis$raw_drop %>% filter(PupSite %in% drop_plot_filter_sites),
    drop_plot_path,
    drop_plot_title
  )
}
```
