## Supplementary material for "Barking up the microbiome tree: a multi-body-site characterization of maternal postpartum and pup early colonization dynamics": Code: Phyloseq_functions.R

```
###########################################Phyloseq_functions.R#########################
plot_PCA <- function(dds,group, set_name,dds_res,res_dir=result_dir,to_label=T){
  #print(paste0(result_dir,"/plot_deseq_",set_name,"_PCA", ".jpg"))
  vstcounts <- varianceStabilizingTransformation(dds, blind=TRUE)
  p<-DESeq2::plotPCA(vstcounts, intgroup= group) + 
    labs(title=paste('PCA', group, set_name))
  if (to_label)     p<-p+geom_text_repel(aes(label=colnames(dds)),vjust=2,  size=4)
  if (!grepl("efratsh",getwd())){
    res_txt<-capture.output(summary(dds_res))
  } else {
    res_txt<-capture.output(DESeq2::summary(dds_res))
  }
  text <- paste(res_txt,collapse="\n ")
  text<-text_grob(text)
  
  together <- arrangeGrob(p, text, ncol = 2,nrow = 1)
  plot(together)
  dev.copy(jpeg,filename=paste0(res_dir,"/plot_deseq_",set_name,"_PCA", ".jpg"), 
           width=1200, height=600)
  dev.off()
}

plot_dds_heatmap <- function(dds,group, set_name, res_dir=result_dir){
  #rld = rlogTransformation(dds)
  vsd <- varianceStabilizingTransformation (dds)
  distsRL <- dist(t(assay(vsd))) # Calculate distances using transformed &normalized counts
  mat <- as.matrix(distsRL) # convert to matrix
  #mat <- mat[, order(colnames(mat))]
  colors <- colorRampPalette( rev(brewer.pal(9, "Blues")) )(255) # set colors
  print(length(group))
  if (length(group)==1){
    colnames(mat)<-paste0(colData(vsd)[,group],"_",colnames(mat))
    rownames(mat)<-paste0(colData(vsd)[,group],"_",rownames(mat))
    pheatmap(mat, legend = TRUE,
             clustering_distance_rows=distsRL,
             clustering_distance_cols=distsRL,
             col=colors, main = paste(set_name,"Sample-to-sample distances (vsd)"))
    
  } else {
    df = as.data.frame(colData(vsd)[,group]) # dataframe with a column groups
    pheatmap(mat, annotation_col=df, annotation_row = df, legend = TRUE,
             clustering_distance_rows=distsRL,
             clustering_distance_cols=distsRL,
             col=colors, main = paste(set_name,"Sample-to-sample distances (vsd)"))
    
  }
  dev.copy(jpeg,filename=paste0(res_dir,"/heatmap_samples_",set_name, ".jpg"), 
           width=600, height=600)
  dev.off()
  
}

plot_lfc <- function(res_df, set_name, no_phylum=FALSE, res_dir=result_dir){
  print(nrow(res_df))
  
  if (no_phylum){
    print(ggplot(res_df, aes(x=short_title, y=log2FoldChange)) +
            geom_jitter(size=3, width = 0.2) + xlab("Taxa") +
            ggtitle(paste("Taxa log2FoldChange",set_name ,"(Alpha=",alpha,")",nrow(res_df))) +
            theme(axis.text.x = element_text(angle = -90, hjust = 0, vjust=0.5, size=14)) +
            coord_flip())
    
  } else{
    print(ggplot(res_df, aes(x=short_title, y=log2FoldChange, color=Phylum)) +
            geom_jitter(size=3, width = 0.2) + xlab("Taxa") +
            ggtitle(paste("Taxa log2FoldChange",set_name ,"(Alpha=",alpha,")",nrow(res_df))) +
            theme(axis.text.x = element_text(angle = -90, hjust = 0, vjust=0.5, size=14)) +
            coord_flip())
  }
  dev.copy(jpeg,filename=paste0(res_dir,"/LFC_",set_name, "_plot.jpg"), 
             width=1100, height=600)
  dev.off()
}


# phyloseq_to_lefse
# this script convert phyloseq object into lefse recognized file format
phyloseq_to_lefse <- function(physeq, st_field="TimeW", nd_field="PatientID",short=F){
  #physeq<-physeq_rarefied     #physeq_rarefied_scaled
  #st_field="TimeW" #day
  #nd_field="PatientID"   #mouseID
  
  if (!short){
    # aggregate to one level
    ps_lefse <- physeq %>% 
  				tax_glom(taxrank = 'Species', NArm = F)
    
    # extract taxonomic data from phyloseq object and then stored in a vector called lefse_tax
    #lefse_tax <- ps_lefse %>% 
    #  tax_table %>% 
    #  data.frame(stringsAsFactors=FALSE) %>% 
    #  dplyr::select(short_title) %>%
    #  mutate(short_title=ifelse(substr(short_title,1,1)==" ",substr(short_title,2,str_length(short_title)),short_title))
    to_clean<- c("Unassigned" ,"unidentified","uncultured","gut")
    lefse_tax <- ps_lefse %>% 
              tax_table %>% 
              data.frame(stringsAsFactors=FALSE) %>% 
              rownames_to_column("OTU") %>%
              dplyr::select(-short_title,-mark)  %>%
      mutate_all(~replace(.,str_detect(.,paste(to_clean,collapse = "|")),"")) %>%
              unite(col= "lefse_title",  Domain:Species, na.rm=TRUE, sep= "|")  #%>%
              #dplyr::filter(lefse_title!="||||||")  %>%
    #take off the last |
    for (i in 1:7){
      lefse_tax<-lefse_tax %>%
                  mutate(lefse_title=ifelse(substr(lefse_title,str_length(lefse_title),str_length(lefse_title))=="|",
                                substr(lefse_title,1,(str_length(lefse_title)-1)),lefse_title))  #%>%
        
    }
  }else{
    lefse_tax <- physeq %>% 
                  tax_table %>% 
                  data.frame(stringsAsFactors=FALSE) %>% 
                  rownames_to_column("OTU") %>%
                  dplyr::select(OTU,short_title)  %>%
                  rename(lefse_title=short_title)
      
  }
  lefse_tax<-lefse_tax %>%
              mutate(lefse_title=str_replace_all(lefse_title,"-","_"))  %>%
              column_to_rownames("OTU")  
  
  # extract otu abundance matrix from phyloeq object and annotated with tax information
  lefse_abundance <- otu_table(ps_lefse) %>% 
						as.data.frame(stringsAsFactors = F) %>% 
            rownames_to_column %>% 
            right_join(rownames_to_column(lefse_tax),by="rowname")%>% 
            column_to_rownames("rowname") %>% 
						t %>% 
						data.frame(stringsAsFactors = F)

  # extract sample matadata and order sample same in lefse_matrix
  lefse_md <- sample_data(ps_lefse)%>% 
          data.frame(stringsAsFactors = F) %>%
          rownames_to_column("sample_name") %>%
					dplyr::select("sample_name",eval(st_field),eval(nd_field)) #%>%  
					#mutate_if(is.factor,as.character())  %>%
					#column_to_rownames("sample_name") #SampleID
  
  # merge 
  lefse_table <- right_join(lefse_md, rownames_to_column(lefse_abundance,"sample_name"), 
                           by = "sample_name") %>% 
                  column_to_rownames("sample_name") %>% 
                  t%>% 
                  data.frame(stringsAsFactors = F)
  lefse_table<- cbind(lefse_table[,ncol(lefse_table)], lefse_table[,1:ncol(lefse_table)-1],stringsAsFactors = F)
  lefse_table[1,1]<-st_field
  lefse_table[2,1]<-nd_field
  
  return(lefse_table)
}

rarefy <- function(physeq, min_sum){
  #physeq<-physeq_filtered
  set.seed(123)
  # Plot the rarefaction curves using vegan function rarecurve()
  rarecurve(t(otu_table(physeq)), step=50, cex=0.5)
  dev.copy(jpeg,filename=paste0(result_dir,"/unrarefied_",set, ".jpg"), 
           width=1000, height=700)
  dev.off()
  #rarecurve(t(otu_table(physeq_clean)), step=50, cex=0.5)
  
  #Rarefy the samples without replacement. Rarefaction is used to simulate even number of reads per sample. In this example, the rarefaction depth chosen is the 80% of the minimum sample depth in the dataset (in this case ? reads per sample).
  
  sample_sums(physeq)
  # take the same number of speciman from each sample
  if (missing(min_sum)){
    physeq_rarefied <- rarefy_even_depth(physeq, rngseed=123,
                                      sample.size=0.99*min(sample_sums(physeq)), replace=T)
  } else {
    physeq_rarefied <- rarefy_even_depth(physeq, rngseed=123,
                                        sample.size=min_sum, replace=T)
  }
  physeq_rarefied
  sample_sums(physeq_rarefied)
  rarecurve(t(otu_table(physeq_rarefied)), step=50, cex=0.5)
  dev.copy(jpeg,filename=paste0(result_dir,"/rarefied_",set, ".jpg"), 
           width=1000, height=700)
  dev.off()
  
  sample_data(physeq_rarefied)
  rank_names(physeq_rarefied)
  sample_variables(physeq_rarefied)
  
  #physeq_rarefied_data <- psmelt(physeq_rarefied)
  assign("physeq_rarefied_data",psmelt(physeq_rarefied), envir = .GlobalEnv)
  
  str(physeq_rarefied_data)
  
  #openxlsx::write.xlsx (physeq_rarefied_data,
  #                      paste0(metadata_dir,"/physeq_rarefied_data_",set,classif,".xlsx"),
  #                      sheetName ="physeq_rarefied_data" ,col.names = TRUE) 
  
  openxlsx::write.xlsx (otu_table(physeq_rarefied),paste0(metadata_dir,"/physeq_rarefied_otu_table.xlsx"),
                        sheetName ="physeq_rarefied_otu_table.xlsx" ,col.names = TRUE,row.names=TRUE) 
  
  return(physeq_rarefied)
}

prep_taxa <- function(tax,classifier="GG"){
  #tax<-taxaGT
  # split taxon column
  taxa_16S <- data.frame(stringr::str_split_fixed(tax$Taxon,";",7),stringsAsFactors=FALSE)
  colnames(taxa_16S) <- c("Domain", "Phylum", "Class", "Order", "Family", "Genus", "Species")
  rownames(taxa_16S) <- tax$OTU.ID

  #add short title
  if (classifier!="GG"){
    taxa_16S <- taxa_16S %>% 
      rownames_to_column("OTU")  %>% 
      mutate(Species=str_replace_all(Species," ","_"))  %>%
      # add short taxa title=last rank
      mutate(short_title=if_else(!endsWith(Species,"_")& 
                                   Species!=""& 
                                   !str_detect(Species,"uncultured") & 
                                   !str_detect(Species,"_gut") & 
                                   !str_detect(Species,"metagenome") & 
                                   !str_detect(Species,"unidentified"),Species, #only GT
                                 if_else(!endsWith(Genus,"_")& 
                                           !str_detect(Genus,"uncultured") & 
                                           Genus!="",Genus,
                                         if_else(!endsWith(Family,"_")& 
                                                   !str_detect(Family,"uncultured") & 
                                                   Family!="",Family,
                                                 if_else(!endsWith(Order,"_")& 
                                                           !str_detect(Order,"uncultured") & 
                                                           Order!="",Order,
                                                         if_else(!endsWith(Class,"_")& Class!="",Class,
                                                                 if_else(!endsWith(Phylum,"_")& Phylum!="",Phylum,
                                                                         Domain))))))) %>%
      #add mark for level
      mutate(mark=case_when(startsWith(short_title,"k__")~1,
                            startsWith(short_title,"p__")~2,
                            startsWith(short_title,"c__")~3,
                            startsWith(short_title,"o__")~4,
                            startsWith(short_title,"f__")~5,
                            startsWith(short_title,"g__")~6,
                            startsWith(short_title,"s__")~7,
                            TRUE ~  0))   %>%
      column_to_rownames("OTU")
  }else{
    taxa_16S <- taxa_16S %>% 
      rownames_to_column("OTU")  %>% 
      # add short taxa title=last rank
      mutate(short_title=if_else(!endsWith(Species,"_")& 
                                   Species!=""& 
                                   !str_detect(Species,"uncultured") & 
                                   !str_detect(Species,"_gut") & 
                                   !str_detect(Species,"metagenome") & 
                                   !str_detect(Species,"unidentified"),
                                 str_replace(str_replace(paste0(Genus,"_",Species)," s__",""),"g__","s__"),
                                 if_else(!endsWith(Genus,"_")& 
                                           !str_detect(Genus,"uncultured") & 
                                           Genus!="",Genus,
                                         if_else(!endsWith(Family,"_")& 
                                                   !str_detect(Family,"uncultured") & 
                                                   Family!="",Family,
                                                 if_else(!endsWith(Order,"_")& 
                                                           !str_detect(Order,"uncultured") & 
                                                           Order!="",Order,
                                                         if_else(!endsWith(Class,"_")& Class!="",Class,
                                                                 if_else(!endsWith(Phylum,"_")& Phylum!="",Phylum,
                                                                         Domain))))))) %>%
      #add mark for level
      mutate(mark=case_when(startsWith(short_title,"k__")~1,
                            startsWith(short_title," p__")~2,
                            startsWith(short_title," c__")~3,
                            startsWith(short_title," o__")~4,
                            startsWith(short_title," f__")~5,
                            startsWith(short_title," g__")~6,
                            startsWith(short_title," s__")~7,
                            TRUE ~  0))   %>%
      column_to_rownames("OTU")
  }
    #remove the level letter
    #mutate_at(vars(!contains("short") & !contains("OTU")),funs(substring(.,5))) %>%
    #dplyr::filter(short_title!="Unassigned") %>%
							 
  print(paste("nrow taxa:",nrow(taxa_16S)))
  return(taxa_16S)
}

corr_gene_tax <- function(in_corr_p, in_corr_r, in_gene_res_pval_sig , in_res_sig_taxa, 
                          in_wb, in_sheet, in_tl_size=.6) {
  #in_corr_p<-corr_p
  #in_corr_r<-corr_r
  #in_gene_res_pval_sig<-gene_sig_lfc
  #in_res_sig_taxa<-res_sig_taxa
  #in_tl_size=.45
  #in_sheet<-sheet_name
  
  print(paste("sheet:",in_sheet))
  #write sig cor genes per OTU to excel
  corr_p_sig<-subset(in_corr_p,pval<=0.05)
  print(paste("no of cor pval<0.05:",nrow(corr_p_sig)))
  if (nrow(corr_p_sig)>0){
    colno<-1
    for (t in unique(corr_p_sig$taxa)){
      #t<-" s__subflava" 
      corr_r_sig_t<-subset(in_corr_r,taxa==t,select=c(gene,r)) 
      corr_p_sig_t<-subset(corr_p_sig,taxa==t,select=gene) %>%
        left_join(corr_r_sig_t,by="gene")
      if (nrow(corr_p_sig_t)>2){
		  result = tryCatch({
			
			anno_t<-
			  getBM(attributes=c('external_gene_name','description', 'gene_biotype'), 
					filters = 'mgi_symbol', #external_gene_name
					values = corr_p_sig_t, 
					mart = mouse_ensembl,useCache = FALSE)
			#setNames(anno_t$external_gene_name,t)
			data.table::setnames(anno_t, "external_gene_name", t)
			print(colnames(anno_t))
			writeData(in_wb, sheet = in_sheet,anno_t , rowNames = F, startCol = colno)
			setColWidths(in_wb, in_sheet, cols = 1:25, widths = "auto")
		  },
		  error = function (condition) {
			writeData(in_wb, sheet = in_sheet,corr_p_sig_t , rowNames = F, startCol = colno)
			setColWidths(in_wb, in_sheet, cols = 1:30, widths = "auto")
		  })
		  colno=colno+3
		}
    setColWidths(in_wb, in_sheet, cols = 1:colno, widths = "auto")
    
   #for(t in unique(in_corr_r$taxa)){
    # corr_r_4plot<-in_corr_r %>%                   
    #    mutate(gene=as.factor(gene)) %>%
    #    dplyr::filter(taxa==t) %>%
    #    arrange( r)
    #  print(ggplot(corr_r_4plot, aes(x=reorder(gene,r), y=r)) + 
    #          geom_point() +
    #          scale_y_continuous()+
    #          ggtitle(paste(in_sheet," correlation for", t)))
    #}
    
    #re-order gene by LFC
    gene_corr<- in_gene_res_pval_sig %>%
                  rownames_to_column() %>%
                  dplyr::filter(rowname %in% unique(in_corr_p$gene)) %>%
                  dplyr::arrange(log2FoldChange) %>%
                  column_to_rownames()
    nrow(gene_corr)
    taxa_corr<- in_res_sig_taxa %>%
                  rownames_to_column() %>%
                  dplyr::filter(rowname %in% unique(in_corr_p$taxa)) %>%
                  arrange(desc(log2FoldChange))  %>%
                  column_to_rownames()
    nrow(taxa_corr)
    
    #pivot_wider (move rows into columns).
    corr_p_spread<- pivot_wider(in_corr_p, names_from=gene, values_from=pval) %>% 
                      column_to_rownames("taxa") %>%
                      relocate(rownames(gene_corr))  %>%
                      as.matrix()
    nrow(corr_p_spread)
    ncol(corr_p_spread)
    #re-order taxa accordong to LFC
    if (nrow(corr_p_spread)>1) {
		corr_p_spread<-corr_p_spread[rownames(taxa_corr),]
    }
    corr_r_spread<- pivot_wider(in_corr_r, names_from =gene, values_from =r) %>% 
                      column_to_rownames("taxa")  %>%
                      relocate(rownames(gene_corr))  %>%
                      as.matrix()
    nrow(corr_p_spread)
    ncol(corr_p_spread)
    #re-order taxa accordong to LFC
    corr_r_spread<-corr_r_spread[rownames(taxa_corr),]
    
    }
#browser()
    #filter for sig correlations
    corr_p_sig_spread <-corr_p_spread%>%
      as.data.frame%>%
      select_if(~ any(.<=0.05))%>%
      filter_all(any_vars(. <=0.05))%>%
      as.matrix()

    #corr_p_sig_spread <-corr_p_spread[,apply(corr_p_spread<0.05, 2, any) ]
    #corr_p_sig_spread <-corr_p_sig_spread[apply(corr_p_sig_spread<0.05, 1, any) ,]
    nrow(corr_p_sig_spread)
    ncol(corr_p_sig_spread)
    
    corr_r_sig_spread<-corr_r_spread%>%
                        as.data.frame%>%
                        rownames_to_column("taxa") %>%
                        dplyr::filter(taxa %in% rownames(corr_p_sig_spread))%>%
                        dplyr::select(taxa,colnames(corr_p_sig_spread)) %>%
                        column_to_rownames("taxa") %>%
                        as.matrix()
    nrow(corr_r_sig_spread)
    ncol(corr_r_sig_spread)
    
    print(paste("no of cor genes pval<0.05(corr_p_sig_spread):",ncol(corr_p_sig_spread)))
    print(paste("no of cor taxa pval<0.05(corr_p_sig_spread):",nrow(corr_p_sig_spread)))
    
    gene_corr_sig<- gene_corr %>%
                      rownames_to_column() %>%
                      dplyr::filter(rowname %in% colnames(corr_p_sig_spread)) #%>%
                      #column_to_rownames()
 
    taxa_corr_sig<- taxa_corr %>%
                      rownames_to_column() %>%
                      dplyr::filter(rowname %in% rownames(corr_p_sig_spread)) #%>%
                      #column_to_rownames()
    
    #par(mfrow=c(2,1))
    corrplot::corrplot(corr_r_spread, p.mat = corr_p_spread, method = "square", is.corr=FALSE,
                       insig = "label_sig", sig.level = c(.001, .01, .05), 
                       col = colorRampPalette(c("blue","white","red"))(100),
                       title=paste("\n\n\nTaxa-significant genes correlation", in_sheet,
                                   "\nno of genes:",ncol(corr_r_spread),
                                   "\nno of TAXAs:",nrow(corr_r_spread)-1),
                      cl.cex=.7, cl.ratio = 0.08,#cl.pos = 'b', cl.offset = 1,
                       tl.cex=in_tl_size, tl.srt=30, tl.col="black",
                       pch.cex = .7, pch.col = "black") #, order="FPC"
     
    
 
      #corrplot only pval<0.05
      
      corrplot::corrplot(corr_r_sig_spread, p.mat = corr_p_sig_spread, method = "square", is.corr=FALSE,
                         insig = "label_sig", sig.level = c(.001, .01, .05), 
                         col = colorRampPalette(c("blue","white","red"))(100),
                         title=paste("\n\n\nOTU-significant genes correlation", in_sheet,
                                     "\nno of genes:",ncol(corr_r_sig_spread),
                                     "\nno of TAXAs:",nrow(corr_r_sig_spread)),
                         cl.cex=.6, cl.ratio = 0.08,#cl.pos = 'b', cl.offset = 1,
                         tl.cex=in_tl_size, tl.srt=60, tl.col="black",
                         pch.cex = .5, pch.col = "black") #, order="FPC"
      
      #p <- grab_grob()
      
      # genes LFC barplot

      print(ggplot(gene_corr_sig, aes(fill=pvalue, 
                                y=log2FoldChange, 
                                x=reorder(rowname,log2FoldChange)))+
        geom_bar(position="dodge", stat="identity")+
        theme_grey(base_size = 10) +
        xlab("Genes")+
        ylab("log2FoldChange") +
        ggtitle(paste("barplot genes sig lfc.",gene_thresh,in_sheet,set))+
        theme(axis.text.x = element_text(angle = 45))+
        theme(panel.grid.major = element_blank(), panel.grid.minor = element_blank(),
              panel.background = element_blank())+
        theme(plot.margin=unit(c(0,1,0,3),"cm")))
  
      # taxa LFC barplot
      print(ggplot(taxa_corr_sig, aes(fill=pvalue, 
                                    y=log2FoldChange, 
                                    x=reorder(rowname,log2FoldChange)))+
              geom_bar(position="dodge", stat="identity")+
              theme_grey(base_size = 20) +
              ggtitle(paste("barplot taxa sig lfc.",taxa_thresh,in_sheet,set))+
              theme(axis.title.y= element_blank(), panel.grid.minor = element_blank(),
                    panel.background = element_blank())+ 
              coord_flip())
      #save(taxa_corr, gene_corr, corr_r_sig_spread,corr_p_sig_spread,
      #     file=paste0(metadata_dir,"/data_4corr_",set,"_",in_sheet,".Rdata"))
    
  }  
}


merge_plot_taxa <- function(physeq_r_scaled, physeq_r_scaled_data, set_name="all",
                            field,field2,field3,rank_name=c("Species","short_title")){
  #set_name="Israel"
  #physeq_r_scaled<-physeq_rarefied_scaled
  #physeq_r_scaled_data<-physeq_rarefied_scaled_data
  #rank_name<-c("short_title")
  #field= "Control_GDM";  field2= "trimester"
  getPalette = colorRampPalette(brewer.pal(9, "Set1"))
  sampleColor <- getPalette(max(nsamples(physeq_r_scaled)))
  #for (r in rank_names(physeq_r_scaled)){
    for (r in rank_name){
      #The option NArm set to FALSE forces the function to keep the unclassified OTUs at the phylum level. 
    #r<-"short_title"
    print(paste("r:",r))
    #scaled  
    physeq_r_scaled_merged <- tax_glom(physeq_r_scaled, taxrank=r, NArm=T)
    tax_table(physeq_r_scaled_merged)<-
      tax_table(physeq_r_scaled)[,1:which(rank_names(physeq_r_scaled)==r)]
    print(physeq_r_scaled_merged)
    #abundances(physeq_r_scaled_merged)
    
    #cleaner bar plot:
    if (r!="Domain" & r!="Phylum"& r!="Class"){
      #r<-"short_title"
      PhylaPalette <- getPalette(max(nrow(unique(as.data.frame(taxa_16S_df[,r])))))
      
      
      print(phyloseq::plot_bar(physeq_r_scaled_merged, fill=r, 
                               title=paste("Abundance",r, "scaled& merged per",eval(field), "-",set_name))+ 
              scale_fill_manual(values = PhylaPalette, guide = guide_legend(nrow=60)) + 
                                #breaks = unique(fct_reorder(taxa_16S_df[[r]], taxa_16S_df$Phylum,min)),
                                #guide=guide_legend(nrow=50)) + 
              facet_wrap(~get(field), scales="free_x", nrow=1))
      dev.copy(jpeg,filename=paste0(result_dir,"/barplot_",r,"_scaled_merged_",eval(field),"_",set_name, ".jpg"), 
               width=nrow(unique(as.data.frame(taxa_16S_df[,r])))*3+2000, height=1200)
      dev.off()
      if(!missing(field2)) {
        print(phyloseq::plot_bar(physeq_r_scaled_merged, fill=r, 
                                 title=paste("Abundance",r, "scaled& merged per",eval(field), "-",set_name))+ 
                scale_fill_manual(values = PhylaPalette, guide = guide_legend(nrow=60)) + 
                facet_wrap(~get(field2), scales="free_x", nrow=1))
        dev.copy(jpeg,filename=paste0(result_dir,"/barplot_",r,"_scaled_merged_",eval(field2),"_",set_name, ".jpg"), 
                 width=nrow(unique(as.data.frame(taxa_16S_df[,r])))*3+1000, height=1200)
        dev.off()
      }        
      #plot_heatmap(physeq )
      print(plot_heatmap(physeq_r_scaled_merged, taxa.label=r, method = "DPCoA", distance = "bray",
                         title=paste("Abundance",r, "scaled& merged-",set_name)))
      ggsave(paste0(result_dir,"/plot_",r,"_scaled_merged_sample_heatmap","_",set_name, ".jpg"), 
             width=10, height=10)
      
      #dev.copy(jpeg,filename=paste0(result_dir,"/plot_",r,
      #                              "_scaled_merged_sample_heatmap","_",set_name, ".jpg"), 
      #         width=1600, height=1000)
      dev.off()
      
      #remove empty data for the rank
#      nrow(physeq_r_data)
#      physeq_r_data_filled=subset(physeq_r_data,physeq_r_data[,r]!= "")
#      nrow(physeq_r_data_filled)
      
      #Plot Abundance Change sacaled 

      nrow(physeq_r_scaled_data)
      physeq_r_scaled_data_filled<-
        subset(physeq_r_scaled_data,physeq_r_scaled_data[,r]!= "")
      nrow(physeq_r_scaled_data_filled)
      
      print(ggplot(physeq_r_scaled_data_filled, 
                   aes(x = get(field), y = Abundance, factor(get(r)), 
                                                           fill = factor(get(r)))) + 
              ylab(r)+ 
              xlab(eval(field)) +
              ggtitle(paste("Microbiota Composition Scaled" ,r,"-",set_name)) + 
              geom_bar(stat = "identity") + 
              facet_wrap(~get(field), scales = "free_x") + 
              scale_fill_manual(values = PhylaPalette) + 
              theme(legend.text=element_text(size=9)) +
              guides(fill=guide_legend(ncol=4)))
      dev.copy(jpeg,filename=paste0(result_dir,"/plot_",r,"_scaled_",eval(field),"_grouped","_",set_name, ".jpg"), 
               width=1600, height=1000)
      dev.off()
      
      if(!missing(field2)) {
        print("field2")
        print(ggplot(physeq_r_scaled_data_filled, 
                     aes(x = get(field2), y = Abundance, factor(get(r)),
                         fill = factor(get(r)))) + 
                ylab(r)+ 
                xlab(eval(field2)) +
                ggtitle(paste("Microbiota Composition per ",eval(field),"+",eval(field2)," Scaled" ,r,"-",set_name)) + 
                geom_bar(stat = "identity") + 
                facet_wrap(~get(field), scales = "free_x") + 
                scale_fill_manual(values = PhylaPalette) + 
                theme(legend.text=element_text(size=9)) +
                guides(fill=guide_legend(ncol=4)))
        dev.copy(jpeg,filename=paste0(result_dir,"/plot_",r,"_scaled_",eval(field2),"_grouped_",eval(field),"_",set_name, ".jpg"), 
                 width=1600, height=1000)
        dev.off()

        print(ggplot(physeq_r_scaled_data_filled, 
                     aes(x = get(field), y = Abundance, factor(get(r)),
                         fill = factor(get(r)))) + 
                ylab(r)+ 
                xlab(eval(field)) +
                ggtitle(paste("Microbiota Composition per ",eval(field2)," Scaled" ,r,"-",set_name)) + 
                geom_bar(stat = "identity") + 
                facet_wrap(~get(field2), scales = "free_x", nrow=1) + 
                scale_fill_manual(values = PhylaPalette) + 
                theme(legend.text=element_text(size=9)) +
                guides(fill=guide_legend(ncol=4)))
        dev.copy(jpeg,filename=paste0(result_dir,"/plot_",r,"_scaled_",eval(field),"_grouped_",eval(field2),"_",set_name, ".jpg"), 
                 width=1600, height=1000)
        dev.off()
      }
      if(!missing(field3)) {
        print("field3")
        print(ggplot(physeq_r_scaled_data_filled, 
                     aes(x = get(field3), y = Abundance, factor(get(r)),
                         fill = factor(get(r)))) + 
                ylab(r)+ 
                xlab(eval(field3)) +
                ggtitle(paste("Microbiota Composition per ",eval(field),"+",eval(field3)," Scaled" ,r,"-",set_name)) + 
                geom_bar(stat = "identity") + 
                facet_wrap(~get(field), scales = "free_x") + 
                scale_fill_manual(values = PhylaPalette) + 
                theme(legend.text=element_text(size=9)) +
                guides(fill=guide_legend(ncol=4)))
        dev.copy(jpeg,filename=paste0(result_dir,"/plot_",r,"_scaled_",eval(field3),"_grouped_",eval(field),"_",set_name, ".jpg"), 
                 width=1600, height=1000)
        dev.off()
        print(ggplot(physeq_r_scaled_data_filled, 
                     aes(x = get(field), y = Abundance, factor(get(r)),
                         fill = factor(get(r)))) + 
                ylab(r)+ 
                xlab(eval(field)) +
                ggtitle(paste("Microbiota Composition per ",eval(field3)," Scaled" ,r,"-",set_name)) + 
                geom_bar(stat = "identity") + 
                facet_wrap(~get(field3), scales = "free_x", nrow=1) + 
                scale_fill_manual(values = PhylaPalette) + 
                theme(legend.text=element_text(size=9)) +
                guides(fill=guide_legend(ncol=4)))
        dev.copy(jpeg,filename=paste0(result_dir,"/plot_",r,"_scaled_",eval(field),"_grouped_",eval(field3),"_",set_name, ".jpg"), 
                 width=1600, height=1000)
        dev.off()
      }  
    }
  }
}

Abundance_boxplot <- function(physeq_data,res_dir=result_dir, lvl="OTU",
                            field="TimeW",field2="PatientID"){
  #physeq_data<-physeq_anova
  #field="RatType"
  #field2="TimeGroup"
  #res_dir=result_dir
  
  wb <- openxlsx::createWorkbook()
  sheet_name <- "taxa_abundance_stats"
  addWorksheet(wb, sheet_name)
  titStyle <- createStyle(fontColour = "#006100", fgFill = "#C6EFCE")
  st_row<-1
  for (sht in unique(physeq_data$short_title)) {
    #sht<-" f__S24-7" #" g__Prevotella" 
    print(sht)
    physeq_data_p<- subset(physeq_data,short_title==sht, select=-short_title)
    print(paste("no of OTUs:", nrow(physeq_data_p)))
    
    # ANOVA-Test whether the groups, TimeW,  differ significantly from each
    if (nrow(subset(physeq_data_p,Abundance>0))>2 & nrow(physeq_data_p)<400){
      # prevent zero values
      physeq_data_p_trans<-physeq_data_p
      physeq_data_p_trans$Abundance<-sqrt (1+physeq_data_p$Abundance)
      #anova <- adonis(Abundance ~ get(field), data=physeq_data_p_trans)
      anova <- adonis(formula =Abundance ~ get(field)+ get(field2), data=physeq_data_p_trans)
      anova2 <- adonis(formula =Abundance ~ get(field2)+ get(field), data=physeq_data_p_trans)
      print(anova2)
      anova_2way <- aov(Abundance ~  get(field2)+get(field), data=physeq_data_p_trans) #
      tukey<- TukeyHSD(anova_2way)
      tukey_df<-rbind(as.data.frame(tukey$`get(field)`), as.data.frame(tukey$`get(field2)`))
      tukey_df_sig<-subset(tukey_df,tukey_df$`p adj` <0.05)
      anova_2way2 <- aov(Abundance ~  get(field)+get(field2), data=physeq_data_p_trans) #
      tukey2<- TukeyHSD(anova_2way2)
      tukey_df2<-rbind(as.data.frame(tukey$`get(field2)`), as.data.frame(tukey$`get(field)`))
      tukey_df_sig2<-subset(tukey_df,tukey_df$`p adj` <0.05)
      if (!is.na(anova$aov.tab$`Pr(>F)`[1]) | 
          !is.na(anova$aov.tab$`Pr(>F)`[2]) | 
          !is.na(anova$aov.tab$`Pr(>F)`[3]) | 
          nrow(tukey_df_sig)>0) {
        writeData(wb,sheet_name , sht, colNames = T, rowNames= T, startRow= st_row) 
        addStyle(wb, 1, style = titStyle, cols=1:1, rows=st_row:st_row, gridExpand = TRUE)
        writeData(wb,sheet_name , anova$aov.tab, colNames = T, rowNames= T, startRow= st_row+1) 
        writeData(wb,sheet_name , anova2$aov.tab, colNames = T, rowNames= T, startRow= st_row+6) 
        writeData(wb,sheet_name , tukey_df, colNames= T, rowNames= T, startRow= st_row+11) 
        writeData(wb,sheet_name , tukey_df2, colNames= T, rowNames= T, startRow= st_row+15) 
        st_row<-st_row+20
        print(ggplot(physeq_data_p, aes(x = get(field), y = Abundance, fill= get(field2))) +
                geom_point(size=2, position=position_jitter(width=0.2), alpha=0.5) +
                xlab(eval(field))+
                labs(fill = eval(field2))+
                theme(axis.text.x = element_text(angle = 30)) +
                ggtitle(paste(sht,"Abundance of",lvl,"per taxa and",field,"boxplot",
                              "\n no of OTUs:", nrow(physeq_data_p),
                              "\navova",eval(field),anova$aov.tab$`Pr(>F)`[1],
                              eval(field2),anova$aov.tab$`Pr(>F)`[2],
                              "\n",eval(field),":",eval(field2),anova$aov.tab$`Pr(>F)`[3],
                              "\n sig tukey:",paste(rownames(tukey_df_sig),collapse = ";"))) +
                geom_boxplot())
        dev.copy(jpeg,filename=paste0(res_dir,"/taxa_abundance_",field,"_boxplot_",sht,"_",lvl,".jpg"), 
                 width=600, height=800)
        dev.off()
      }
      if (!is.na(anova2$aov.tab$`Pr(>F)`[1]) |
          !is.na(anova2$aov.tab$`Pr(>F)`[2]) |
          !is.na(anova2$aov.tab$`Pr(>F)`[3]) |
          nrow(tukey_df_sig)>0) {
        print(ggplot(physeq_data_p, aes(x = get(field2), y = Abundance, fill= get(field))) +
                geom_point(size=2, position=position_jitter(width=0.2), alpha=0.5) +
                xlab(eval(field2))+
                labs(fill = eval(field))+
                theme(axis.text.x = element_text(angle = 30)) +
                ggtitle(paste(sht,"Abundance of",lvl,"per taxa and ",field2,"boxplot",
                              "\n no of OTUs:", nrow(physeq_data_p),
                              "\navova2",eval(field2),anova2$aov.tab$`Pr(>F)`[1],
                              eval(field),anova2$aov.tab$`Pr(>F)`[2],
                              "\n",eval(field2),":",eval(field),anova2$aov.tab$`Pr(>F)`[3],
                              "\n sig tukey2:",paste(rownames(tukey_df_sig2),collapse = ";"))) +
                geom_boxplot())
        dev.copy(jpeg,filename=paste0(res_dir,"/taxa_abundance_",field2,"_boxplot_",sht,"_",lvl,".jpg"), 
                 width=600, height=800)
        dev.off()
      }
      
    }
  }
  setColWidths(wb, sheet_name, cols = 1:8, widths = "auto")
  openxlsx::saveWorkbook(wb, file=paste0(res_dir,"/taxa_abundance_stats_",lvl,"_",set,classif, ".xlsx"), 
                         overwrite = TRUE)
  
  
}


 taxa_sig_pairs<- function(taxa_thresh, taxa_all_results,set_name,element="taxa",
                           res_dir=result_dir, anno_df=taxa_all_df, 
                           use_padj=FALSE, pthresh=0.05,corr=TRUE, out_format="jpg"){
   #taxa_all_results<- taxa_MW_res # func_all_res  taxa_all_res metabo_all_res
   #taxa_thresh<-1  #0.58
   #res_dir<- result_dir  #result_dir #result_dir_all MB_result_dir
   #set_name<-"" #
   #corr=F ; use_padj=T
   #anno_df=meta_4MA  #taxa_all_df
   print(set_name)
   wb <- openxlsx::createWorkbook()
   res_sig_all<-rownames_to_column(as.data.frame(get(taxa_all_results[1])[0,]),"short_title")
   first_res<-TRUE
   for (res in taxa_all_results){
     #res <- taxa_all_results[2]
     print(res)
     sheet_name<-str_replace(res,"taxa_","")
     if (str_length(sheet_name)>31) sheet_name<-str_split_fixed(sheet_name,"\\_",2)[,2]
     if (str_length(sheet_name)>31) sheet_name<-str_sub(sheet_name,1,31)
     addWorksheet(wb,sheet_name )
     
     compared<-str_split_fixed(res,"\\_",4)[,1]
     res_name<-str_split_fixed(res,"\\_",2)[,2]
     
     res_df<- as.data.frame(get(res))
     print(paste("nrow res_df:",nrow(res_df)))
     ## Volcano plot t
     pv <- plot_volcano(res_df,paste0(set,"_16s_",compared,"_pairs_sig",
                                      ifelse(use_padj,paste0("PADJ.",pthresh), 
                                             paste0("pval lfc.",taxa_thresh)),"_", res,"_",set_name),
                        res_dir = res_dir,
                        lfc=taxa_thresh,
                        padj_thresh=pthresh)
     
     ## calculate sig taxa
     res_padj <- subset(res_df, padj <= pthresh)
     print(paste(res, "padj sig:",nrow(res_padj)))
     
     res_pval <- subset(res_df, pvalue <= pthresh)
     print(paste(res, "pvalue <=:",pthresh,nrow(res_pval)))
     
     if (!use_padj){
       res_sig <- as.data.frame(subset(res_pval, abs(log2FoldChange) >=taxa_thresh))
       print(paste(compared,"pval sig",taxa_thresh,nrow(res_sig)))
       assign(paste0(res,"_pval_sig"),res_sig, envir = .GlobalEnv)
     } else {
#browser()
       res_sig <- as.data.frame(subset(res_padj, abs(log2FoldChange) >=taxa_thresh))
       print(paste(compared,"padj sig",taxa_thresh,nrow(res_sig)))
       assign(paste0(res,"_padj_sig"),res_sig, envir = .GlobalEnv)
     }
     writeData(wb,sheet_name , res_sig, colNames = TRUE, rowNames = TRUE) 
     
     assign(paste0("res_",set_name,"_",res_name),res_sig, envir = .GlobalEnv)
    #4correlation
     if (corr){
       taxa_abund<-get(paste0(element,"_count_",set_name))
       if (!use_padj){
         taxa_abund_all<- taxa_abund  %>%
           rownames_to_column("short_title") %>%
           dplyr::filter(short_title %in% rownames(res_pval)) %>%
           column_to_rownames("short_title") 
       } else {
         taxa_abund_all<- taxa_abund  %>%
           rownames_to_column("short_title") %>%
           dplyr::filter(short_title %in% rownames(res_padj)) %>%
           column_to_rownames("short_title") 
       }
       #print(rownames(res_sig))
       taxa_abund_sig<- taxa_abund  %>%
         rownames_to_column("short_title") %>%
         dplyr::filter(short_title %in% rownames(res_sig)) %>%
         column_to_rownames("short_title") 
       #print(rownames(taxa_abund_sig))
       
       #4corr
       res_sig_taxa<-res_sig
       res_taxa_pval<-res_pval
#browser()
       save(res_sig_taxa, taxa_abund_sig, res_taxa_pval,taxa_abund_all,
            file=paste0(metadata_dir,"/sig.",taxa_thresh,"_",element,"_abund_4corr_",res,"_",set,"_",set_name, ".Rdata"))
       
     } 
     res_sig_all<-res_sig  %>%
         rownames_to_column("short_title") %>%
         mutate(pair=res) %>%
         bind_rows(res_sig_all)
     if (nrow(res_sig)>1){
		 # LFC barplot
		 tit<-paste("barplot",compared,"sig",ifelse(use_padj,"PADJ", paste0("lfc.",taxa_thresh)),res,set,set_name)
#browser()
		 print(ggplot(rownames_to_column(res_sig), aes(fill=pvalue, 
															y=log2FoldChange, 
															x=reorder(substr(rowname,1,60),log2FoldChange)))+
				 geom_bar(position="dodge", stat="identity")+
				 #theme_grey(base_size = 20) +
				 theme(panel.grid.major = element_blank(), 
					   panel.grid.minor = element_blank(),
					   panel.background = element_blank(), 
					   axis.line = element_line(colour = "black"),
					   text=element_text(size = 20,  family="sans") )+ 
				 ggtitle(tit)+
				 theme(axis.title.y= element_blank())+#, 
				 coord_flip())
		 if (out_format=="jpg"){
		   dev.copy(jpeg,filename=paste0(res_dir,"/barplot_",compared,"_sig_",
										 ifelse(use_padj,"PADJ", paste0("lfc.",taxa_thresh)),"_",res,"_",set,"_",set_name, ".jpg"), 
					width=max((max(res_sig$log2FoldChange)-min(res_sig$log2FoldChange))*20,str_length(tit)*25), 
					height=nrow(res_sig)*45)
		 } else {
		   ggsave(paste0(res_dir,"/barplot_",compared,"_sig_",
						 ifelse(use_padj,"PADJ", paste0("lfc.",taxa_thresh)),"_",res,"_",set,".pdf"), width = 5, height = 5, device = "pdf", dpi=300, units="in")
		 }
		 dev.off()
		 #openxlsx::write.xlsx(res_sig,paste0(res_dir,"/",compared,"_sig_lfc.",taxa_thresh,"_",res,"_",set,".xlsx"),
		#                      row.names = TRUE, col.names = TRUE)
     
     #plot sample-taxa heatmap
     # only the relevant cols
     print(str_split(res,"_",simplify = T))
     ana_type<-ncol(str_split(res,"_",simplify = T))
     if ((ana_type==4 | ana_type==6) & !str_detect(res,"_y_")){
       vst_pair_df<-vst_all
       cond_df<-anno_df
     }else{
       vst_pair_df<-get (str_replace(res,compared,"vst"))
       # only the relevant rows
       cond_df<-get (str_replace(res,compared,"anno"))
     }
     
     # heatmap for all base on the first result
    vst_sig<-vst_pair_df[rownames(res_sig),] %>%
                rownames_to_column() %>%
                mutate(rowname=substr(rowname,1,60)) %>%
                column_to_rownames()
        print(paste("vst_sig",nrow(vst_sig)))
#browser()       
    if (nrow(vst_sig)>1){
       
       result = tryCatch({
         plot_genes_heatmap(vst_sig,
                            cond_df,
                            y_name=compared,
                            paste0(set,"_",res,"_sig",ifelse(use_padj,"PADJ", paste0("lfc.",taxa_thresh)),"_",set_name), scl="row",
                            res_dir=res_dir, 
                            plot_title = paste(set,res,"sig",ifelse(use_padj,"PADJ", paste0("lfc.",taxa_thresh)),set_name,"\nscale=row") )
       },
       error = function (condition) {
         plot_genes_heatmap(vst_sig,
                            cond_df,
                            y_name=compared,
                            paste0(set,"_",res,"_sig",ifelse(use_padj,"PADJ", paste0("lfc.",taxa_thresh)),"_",set_name), 
                            scl="none",
                            res_dir=res_dir, 
                            plot_title = paste(set,,res,"sig",ifelse(use_padj,"PADJ", paste0("lfc.",taxa_thresh)),set_name,"\nscale=none") )
         
       })
       
       }
     }
    openxlsx::saveWorkbook(wb, file=paste0(res_dir,"/",proj,"_",compared,"_sig_",ifelse(use_padj,"PADJ", paste0("lfc.",taxa_thresh)),
                                           set,classif,"_",set_name, ".xlsx"), 
                           overwrite = TRUE)
    
   } 
   #print("after loop")
   #print(res_sig_taxa_all)
   if (nrow(res_sig_all)>1){
     print(ggplot(res_sig_all, aes(fill=pvalue, 
                                  y=log2FoldChange, 
                                  x=reorder(substr(short_title,1,60),log2FoldChange)))+
       geom_bar(position="dodge", stat="identity")+
       theme_grey(base_size = 32) +
       ggtitle(paste("barplot",compared,"sig",ifelse(use_padj,"PADJ", paste("lfc.",taxa_thresh)),set,set_name))+
       theme(axis.title.y= element_blank())+#, 
       facet_wrap(~pair, strip.position = "top",drop=FALSE, scales = "fixed",nrow = 1)+
       coord_flip())
  #browser()   
     w<-max(max((max(res_sig_all$log2FoldChange)-min(res_sig_all$log2FoldChange))*
                  length(unique(res_sig_all$pair)) *30,str_length(tit)*25),
            max(str_length(unique(substr(res_sig_all$short_title,1,60)))))
     print(w)
     dev.copy(jpeg,filename=paste0(res_dir,"/barplot_",compared,"_sig_",
                                   ifelse(use_padj,"PADJ", paste0("lfc.",taxa_thresh)),"_",set,"_",set_name, ".jpg"), 
              width=w, 
              height=nrow(res_sig_all)*30)
     dev.off()
   }
   return(res_sig_all)
 }
 
 
 genus_contaminantion_check<- function(genus2check){
   contaminant_Alpha_proteobacteria<-c("Afipia", "Aquabacterium", "Asticcacaulis", "Aurantimonas", "Beijerinckia", "Bosea", "Bradyrhizobium", "Brevundimonas" , "Caulobacter", "Craurococcus", "Devosia", "Hoeflea", "Mesorhizobium", "Methylobacterium", "Novosphingobium", "Ochrobactrum", "Paracoccus", "Pedomicrobium", "Phyllobacterium", "Rhizobium", "Roseomonas", "Sphingobium", "Sphingomonas", "Sphingopyxis")
   contaminant_Beta_proteobacteria<-c("Acidovorax", "Azoarcus", "Azospira", "Burkholderia", "Comamonas", "Cupriavidus", "Curvibacter", "Delftia", "Duganella", "Herbaspirillum", "Janthinobacterium", "Kingella", "Leptothrix", "Limnobacter", "Massilia", "Methylophilus", "Methyloversatilis", "Oxalobacter", "Pelomonas", "Polaromonas", "Ralstonia", "Schlegelella", "Sulfuritalea", "Undibacterium", "Variovorax")
   contaminant_Gamma_proteobacteria<-c("Acinetobacter" , "Enhydrobacter", "Enterobacter", "Escherichia", "Nevskia", "Pseudomonas", "Pseudoxanthomonas", "Psychrobacter", "Stenotrophomonas", "Xanthomonas")
   contaminant_Actinobacteria<-c("Aeromicrobium", "Arthrobacter", "Beutenbergia", "Brevibacterium", "Corynebacterium", "Curtobacterium", "Dietzia", "Geodermatophilus", "Janibacter", "Kocuria", "Microbacterium", "Micrococcus", "Microlunatus", "Patulibacter", "Propionibacterium", "Rhodococcus", "Tsukamurella")
   contaminant_other<-c("Abiotrophia", "Bacillus", "Brevibacillus", "Brochothrix", "Facklamia", "Paenibacillus", "Streptococcus","Chryseobacterium", "Dyadobacter", "Flavobacterium", "Hydrotalea", "Niastella", "Olivibacter", "Pedobacter", "Wautersiella", "Deinococcus","Acidobacteria")
   if(any(genus2check$Genus %in% contaminant_Alpha_proteobacteria, na.rm = FALSE)){
     ifelse(genus2check$Genus %in% contaminant_Alpha_proteobacteria,taxa_16S_df$Genus,NA)
   } else print("contaminant_Alpha_proteobacteria not contaminanted")
   if(any(genus2check$Genus %in% contaminant_Beta_proteobacteria, na.rm = FALSE)){
     ifelse(genus2check$Genus %in% contaminant_Beta_proteobacteria,taxa_16S_df$Genus,NA)
   } else print("contaminant_Beta_proteobacteria not contaminanted")
   if(any(genus2check$Genus %in% contaminant_Gamma_proteobacteria, na.rm = FALSE)){
     ifelse(genus2check$Genus %in% contaminant_Gamma_proteobacteria,taxa_16S_df$Genus,NA)
   }else print("contaminant_Gamma_proteobacteria not contaminanted")
   if(any(genus2check$Genus %in% contaminant_Actinobacteria, na.rm = FALSE)){
     ifelse(genus2check$Genus %in% contaminant_Actinobacteria,taxa_16S_df$Genus,NA)
   }else print("contaminant_Actinobacteria not contaminanted")
   if(any(genus2check$Genus %in% contaminant_other, na.rm = FALSE)){
     ifelse(genus2check$Genus %in% contaminant_other,taxa_16S_df$Genus,NA)
   }else print("contaminant_other not contaminanted")
 }
 
 Bray_dist<- function(count,md,group,set_name="all", res_dir = result_dir){
   #count<-otu_table(physeq_rarefied_LV)
   #md<-as.data.frame(sample_data(physeq_rarefied_LV)$Condition)
   #group<-"sample_data(physeq_rarefied_LV)$Condition"

   ## Bray-Curtis distances between samples
   dis <- vegdist(t(count))
   ## Calculate multivariate dispersions. Both permdisp and betadisper accept a single "grouping" factor 
   mod <- betadisper(dis, md[[group]]) #, type = "median", bias.adjust=TRUE
   print(mod)
   
   # tests if centroid distances are significantly different from each other
   anova_mod<-anova(mod)
   ## Permutation test for F
   #densityplot(permustats(permutest(mod, pairwise = TRUE, permutations = 99)), scales = list(x = list(relation = "free")))
   
   ## Tukey's Honest Significant Differences
   mod_tukey <- TukeyHSD(mod)
   tukey_df<-as.data.frame(mod_tukey$group)
   tukey_df_sig<-subset(tukey_df,tukey_df$`p adj` <0.1)
   ##Differences in mean levels of the group
   plot(mod_tukey)
#browser()
   plot(mod, ellipse = TRUE, hull = FALSE, conf = 0.90, xlab=group, ylab="Bray-Curtis distance to centroid",
        main=paste("Bray-Curtis distances between samples",set_name,
                   "\nAnova",round(anova_mod$`Pr(>F)`[1],digits = 2),
                   paste(" sig tukey:",paste0(rownames(tukey_df_sig)," (",round(tukey_df_sig$`p adj`,digits = 2),") ", collapse = ";")))) # 90% data ellipse
   text( mod$vectors[,1:2],labels = rownames(mod$vectors), cex = 0.5,pos = 1)
   
   dev.copy(jpeg,filename=paste0(res_dir,"/Dist_samples_",group,"_",set,"_",set_name, ".jpg"))
   dev.off()
   boxplot(mod,xlab=group, ylab="Bray-Curtis distance to centroid",las=2,cex.lab=1,cex.axis=.7,
           main=paste("Distances between samples",set_name,                 
                      "\nAnova",round(anova_mod$`Pr(>F)`[1],digits = 2),
                      paste(" sig tukey:",paste0(rownames(tukey_df_sig)," (",round(tukey_df_sig$`p adj`,digits = 2),") ", collapse = ";")))) 
   dev.copy(jpeg,filename=paste0(res_dir,"/Dist_samples_",group,"_boxplot_",set,"_",set_name, ".jpg"))
   dev.off()
 }
 
 Adiver<- function(wb_in=wb, phy_rarefied,field1,field2,field3,color_field2=TRUE,fill_field3=TRUE, plus_f3=T,
                   set_name="all",PD=TRUE,sheet_a= NULL){
   #phy_rarefied<-physeq_rarefied
   #field1<- "Control_GDM" #RatType TimeW Condition
   #field2<- "trimester" #TimeGroup PatientID Wasp
   #field3<- "woman" #RatID batch Stage
   #color_field2=T ;fill_field3=F ; plus_f3=T
   print(field1)
   set.seed(42)
   posStyle <- createStyle(fontColour = "#006100", fgFill = "#C6EFCE")
   s <- createStyle(numFmt = "$ #,##0.00")

   #m<-"Observed"
   titStyle <- createStyle(fontColour = "#006100", fgFill = "#C6EFCE")
   #sheet_a_divers <- paste0("Adivers ",sheet_a)
#   sheet_wilcox <- "alpha_pairwise_wilcox"
   sheet_a_anova <- paste0("Adivers_anova ",sheet_a)
   #addWorksheet(wb_in, sheet_a_divers)
#   addWorksheet(wb_in, sheet_wilcox)
   addWorksheet(wb_in, sheet_a_anova)
   
   #-------------faithPD- species richness (SR) and phylogenetic diversity (PD) ------
   if (PD){
     alpha_diversity_pd<-estimate_pd(phy_rarefied)
     pairwise.wilcox.test_PD <- NULL
     pairwise.wilcox.test_SR <- NULL
     PD_tukey_df_sig <-NULL
     SR_tukey_df_sig <-NULL
     
     if(!missing(field3)) {
       alpha_divers_pd_meta<- alpha_diversity_pd %>%
         rownames_to_column("rowname") %>%
         left_join(rownames_to_column(meta_16S,"rowname"),by="rowname") %>%
         dplyr::select(rowname,PD,SR,eval(field1),eval(field2),eval(field3)) %>%
         mutate_if(is.integer,as.factor)  %>%
         column_to_rownames("rowname")
       if (plus_f3){
         print("plus_f3")
          anova_PD <- adonis(formula =PD ~ get(field1)*get(field2)+get(field3), data=alpha_divers_pd_meta)
          anova_PD_call<-paste("PD ~ ",field1,"*",field2,"+",field3)
          anova_2way <- aov(PD ~ get(field1)*get(field2)+get(field3), data=alpha_divers_pd_meta) #
          tukey<- TukeyHSD(anova_2way)
          tukey_df<-as.data.frame(rbind(tukey$`get(field1)`,
                                        tukey$`get(field2)`,
                                        tukey$`get(field3)`,
                                        tukey$`get(field1):get(field2)`,
                                        tukey$`get(field2):get(field3)`))
          PD_tukey_df_sig<-subset(tukey_df,tukey_df$`p adj` <0.05)
          anova_SR <- adonis(formula =SR ~ get(field1)*get(field2)+get(field3), data=alpha_divers_pd_meta)
          anova_SR_call<-paste("SR ~ ",field1,"*",field2,"+",field3)
          anova_2way <- aov(PD ~ get(field1)*get(field2)+get(field3), data=alpha_divers_pd_meta) #
          tukey<- TukeyHSD(anova_2way)
          tukey_df<-as.data.frame(rbind(tukey$`get(field1)`,
                                        tukey$`get(field2)`,
                                        tukey$`get(field3)`,
                                        tukey$`get(field1):get(field2)`,
                                        tukey$`get(field2):get(field3)`))
          SR_tukey_df_sig<-subset(tukey_df,tukey_df$`p adj` <0.05)
          print(SR_tukey_df_sig)
          
       } else {
         anova_PD <- adonis(formula =PD ~ get(field1)*get(field2)*get(field3), data=alpha_divers_pd_meta)
         anova_PD_call<-paste("PD ~ ",field1,"*",field2,"*",field3)
         anova_2way <- aov(PD ~ get(field1)*get(field2)*get(field3), data=alpha_divers_pd_meta) #
         tukey<- TukeyHSD(anova_2way)
         tukey_df<-as.data.frame(rbind(tukey$`get(field1)`,
                                       tukey$`get(field2)`,
                                       tukey$`get(field3)`,
                                       tukey$`get(field1):get(field2)`,
                                       tukey$`get(field2):get(field3)`,
                                       tukey$`get(field1):get(field2):get(field3)`))
         PD_tukey_df_sig<-subset(tukey_df,tukey_df$`p adj` <0.05)
         anova_SR <- adonis(formula =SR ~ get(field1)*get(field2)*get(field3), data=alpha_divers_pd_meta)
         anova_SR_call<-paste("SR ~ ",field1,"*",field2,"*",field3)
         print(PD_tukey_df_sig)
         anova_2way <- aov(SR ~ get(field1)*get(field2)*get(field3), data=alpha_divers_pd_meta) #
         tukey<- TukeyHSD(anova_2way)
         tukey_df<-as.data.frame(rbind(tukey$`get(field1)`,
                                       tukey$`get(field2)`,
                                       tukey$`get(field3)`,
                                       tukey$`get(field1):get(field2)`,
                                       tukey$`get(field2):get(field3)`,
                                       tukey$`get(field1):get(field2):get(field3)`))
         SR_tukey_df_sig<-subset(tukey_df,tukey_df$`p adj` <0.05)
       }
       
       print(anova_PD)
       print(anova_SR)
       if(color_field2) {
         if(fill_field3) {
           g<-ggplot(alpha_divers_pd_meta, aes(x=get(field1), y=log(PD), color=get(field2), fill=get(field3)))+
            scale_color_discrete(name = eval(field2)) +
            scale_fill_manual(name = eval(field3),values = rainbow(length(levels(alpha_divers_pd_meta[[eval(field3)]]))))
           g2<-ggplot(alpha_divers_pd_meta, aes(x=get(field1), y=log(SR), color=get(field2), fill=get(field3)))+
            scale_color_discrete(name = eval(field2)) +
            scale_fill_manual(name = eval(field3),values = rainbow(length(levels(alpha_divers_pd_meta[[eval(field3)]]))))
         } else{
           g<-ggplot(alpha_divers_pd_meta, aes(x=get(field1), y=log(PD), color=get(field2)))+
             scale_color_discrete(name = eval(field2) )
           g2<-ggplot(alpha_divers_pd_meta, aes(x=get(field1), y=log(SR), color=get(field2)))+
             scale_color_discrete(name = eval(field2)) 
          }
        } else{
          print("1-field1 only")
           g<-ggplot(alpha_divers_pd_meta, aes(x=get(field1), y=log(PD))+
                       stat_compare_means(method = "anova"))
         g2<-ggplot(alpha_divers_pd_meta, aes(x=get(field1), y=log(SR))+
                      stat_compare_means(method = "anova"))
       }
       if (plus_f3){
         print("plus_f3 plot")
#browser()         
         g<- g+ggtitle(paste("FaithPD-phylogenetic diversity (PD) (log)", sheet_a,
                             "\nAnova",eval(field1), anova_PD$aov.tab$`Pr(>F)`[1],
                             "\nAnova",eval(field2),anova_PD$aov.tab$`Pr(>F)`[2],
                             "\nAnova",eval(field3),anova_PD$aov.tab$`Pr(>F)`[3],
                             "\nAnova",eval(field1),":",eval(field2),anova_PD$aov.tab$`Pr(>F)`[4],
                             "\nTukey sig",paste(rownames(PD_tukey_df_sig), collapse = ";") ))
         g2<- g2+ggtitle(paste("FaithPD-species richness (SR) (log)", sheet_a,
                               "\nAnova",eval(field1), anova_SR$aov.tab$`Pr(>F)`[1],
                               "\nAnova",eval(field2),anova_SR$aov.tab$`Pr(>F)`[2],
                               "\nAnova",eval(field3),anova_SR$aov.tab$`Pr(>F)`[3],
                               "\nAnova",eval(field1),":",eval(field2),anova_SR$aov.tab$`Pr(>F)`[4],
                               "\nTukey sig",paste(rownames(SR_tukey_df_sig), collapse = ";") ))
       }else{
         g<- g+ggtitle(paste("FaithPD-phylogenetic diversity (PD) (log)", sheet_a,
                             "\nAnova",eval(field1), anova_PD$aov.tab$`Pr(>F)`[1],
                             "\nAnova",eval(field2),anova_PD$aov.tab$`Pr(>F)`[2],
                             "\nAnova",eval(field3),anova_PD$aov.tab$`Pr(>F)`[3],
                             "\nAnova",eval(field1),":",eval(field2),anova_PD$aov.tab$`Pr(>F)`[4],
                             "\nAnova",eval(field1),":",eval(field3),anova_SR$aov.tab$`Pr(>F)`[5],
                             "\nAnova",eval(field1),":",eval(field2),":",eval(field3),anova_SR$aov.tab$`Pr(>F)`[6],
                             "\nTukey sig",paste(rownames(PD_tukey_df_sig), collapse = ";")  ))
         g2<- g2+ggtitle(paste("FaithPD-species richness (SR) (log)", sheet_a,
                               "\nAnova",eval(field1), anova_SR$aov.tab$`Pr(>F)`[1],
                               "\nAnova",eval(field2),anova_SR$aov.tab$`Pr(>F)`[2],
                               "\nAnova",eval(field3),anova_SR$aov.tab$`Pr(>F)`[3],
                               "\nAnova",eval(field1),":",eval(field2),anova_SR$aov.tab$`Pr(>F)`[4],
                               "\nAnova",eval(field1),":",eval(field3),anova_SR$aov.tab$`Pr(>F)`[5],
                               "\nAnova",eval(field1),":",eval(field2),":",eval(field3),anova_SR$aov.tab$`Pr(>F)`[6],
                               "\nTukey sig",paste(rownames(SR_tukey_df_sig), collapse = ";")  ))
       }
       print(g+ 
               xlab(eval(field1))+
               geom_boxplot()+
               theme_bw() + 
               theme(axis.text.x = element_text(angle = 90),
                     text = element_text(size = 12))+
               stat_compare_means(method = "anova")) #, axis.text.x = element_blank()))
       ggsave(paste0(result_dir,"/FaithPD_PD_",eval(field1),"_boxplot_",proj,"_",set,"_",sheet_a,".jpg"), 
              width=7, height=5)
       dev.off()
       #print(g2+ 
      #         xlab(eval(field1))+
      #         geom_boxplot()+
      #         theme_bw() + 
      #         theme(axis.text.x = element_text(angle = 90),
      #               text = element_text(size = 12))) #, axis.text.x = element_blank()))
      # ggsave(paste0(result_dir,"/FaithPD_SR_",eval(field1),"_boxplot_",proj,"_",set,"_",sheet_a,".jpg"), 
      #        width=7, height=5)
      # dev.off()
     } else if(!missing(field2)){
     print("field2")
#browser()
     alpha_divers_pd_meta<- alpha_diversity_pd %>%
       rownames_to_column("rowname") %>%
       left_join(rownames_to_column(meta_16S,"rowname"),by="rowname") %>%
       dplyr::select(rowname,PD,SR,eval(field1),eval(field2)) %>%
       dplyr::filter( str_length(get(field1))>0  & !is.na(get(field1)) & str_length(get(field2))>0  & !is.na(get(field2))) %>%
       column_to_rownames("rowname")
     
     anova_PD <- adonis(formula =PD ~ get(field1)*get(field2), data=alpha_divers_pd_meta)
     anova_PD_call<-paste("PD ~ ",field1,"*",field2)
     print(anova_PD)
     print(anova_PD_call)
     anova_2way <- aov(PD ~ get(field1)*get(field2), data=alpha_divers_pd_meta) #
     tukey<- TukeyHSD(anova_2way)
     tukey_df<-as.data.frame(rbind(tukey$`get(field1)`,
                                   tukey$`get(field2)`,
                                   tukey$`get(field1):get(field2)`))
     PD_tukey_df_sig<-subset(tukey_df,tukey_df$`p adj` <0.05)
     print(PD_tukey_df_sig)
     anova_SR <- adonis(formula =SR ~ get(field1)*get(field2), data=alpha_divers_pd_meta)
     anova_SR_call<-paste("SR ~ ",field1,"*",field2)
     anova_SR
     anova_2way <- aov(SR ~ get(field1)*get(field2), data=alpha_divers_pd_meta) #
     tukey<- TukeyHSD(anova_2way)
     tukey_df<-as.data.frame(rbind(tukey$`get(field1)`,
                                   tukey$`get(field2)`,
                                   tukey$`get(field1):get(field2)`))
     SR_tukey_df_sig<-subset(tukey_df,tukey_df$`p adj` <0.05)
     
     print(ggplot(alpha_divers_pd_meta, aes(x=get(field1), y=log(PD), color=get(field2))) +
             ggtitle(paste("FaithPD-phylogenetic diversity (PD) (log)", sheet_a,
                           "\nAnova",eval(field1), anova_PD$aov.tab$`Pr(>F)`[1],
                           "\nAnova",eval(field2),anova_PD$aov.tab$`Pr(>F)`[2],
                           "\nAnova",eval(field1),":",eval(field2),anova_PD$aov.tab$`Pr(>F)`[3],
                           "\nTukey sig",paste(rownames(PD_tukey_df_sig), collapse = ";") )) +  
             geom_boxplot()+
             theme_bw() + 
             theme(axis.text.x = element_text(angle = 90),
                   text = element_text(size = 12))+
             stat_compare_means(method = "anova")) #, axis.text.x = element_blank()))
     ggsave(paste0(result_dir,"/FaithPD_PD_",eval(field1),"_boxplot_",proj,"_",set,"_",sheet_a,".jpg"), 
            width=7, height=5)
     dev.off()
     print(ggplot(alpha_divers_pd_meta, aes(x=get(field1), y=log(SR), color=get(field2))) +
             ggtitle(paste("FaithPD-species richness (SR) (log)", sheet_a,
                           "\nAnova",eval(field1), anova_SR$aov.tab$`Pr(>F)`[1],
                           "\nAnova",eval(field2),anova_SR$aov.tab$`Pr(>F)`[2],
                           "\nAnova",eval(field1),":",eval(field2),anova_SR$aov.tab$`Pr(>F)`[3],
                           "\nTukey sig",paste(rownames(SR_tukey_df_sig), collapse = ";") )) +  
             geom_boxplot()+
             theme_bw() + 
             theme(axis.text.x = element_text(angle = 90),
                   text = element_text(size = 12))+
             stat_compare_means(method = "anova")) #, axis.text.x = element_blank()))
     ggsave(paste0(result_dir,"/FaithPD_SR_",eval(field1),"_boxplot_",proj,"_",set,"_",sheet_a,".jpg"), 
            width=7, height=5)
     dev.off()
     }else{
      print("2-field1 only")
      alpha_divers_pd_meta<- alpha_diversity_pd %>%
        rownames_to_column("rowname") %>%
        left_join(rownames_to_column(meta_16S,"rowname"),by="rowname") %>%
        dplyr::select(rowname,PD,SR,eval(field1)) %>%
        dplyr::filter( str_length(get(field1))>0  & !is.na(get(field1)) ) %>%
        column_to_rownames("rowname")
      print(field1)
      anova_PD <- adonis(formula =PD ~ get(field1), data=alpha_divers_pd_meta)
      anova_PD_call<-paste("PD ~ ",field1)
      anova_PD
      print(anova_PD)
      print(anova_PD_call)
      anova_2way <- aov(PD ~ get(field1), data=alpha_divers_pd_meta) #
      tukey<- TukeyHSD(anova_2way)
      tukey_df<-as.data.frame(rbind(tukey$`get(field1)`))
      PD_tukey_df_sig<-subset(tukey_df,tukey_df$`p adj` <0.05)
      print(PD_tukey_df_sig)
      anova_SR <- adonis(formula =SR ~ get(field1), data=alpha_divers_pd_meta)
      anova_SR_call<-paste("SR ~ ",field1) 
      print(anova_SR)
      anova_2way <- aov(SR ~ get(field1), data=alpha_divers_pd_meta) #
      tukey<- TukeyHSD(anova_2way)
      tukey_df<-as.data.frame(rbind(tukey$`get(field1)`))
      SR_tukey_df_sig<-subset(tukey_df,tukey_df$`p adj` <0.05)
      
      print("pairwise.wilcox.test_PD")
      pairwise.wilcox.test_PD <- pairwise.wilcox.test(alpha_divers_pd_meta$PD,alpha_divers_pd_meta[[field1]],p.adjust.method="none" )
      pairwise.wilcox.test_SR <- pairwise.wilcox.test(alpha_divers_pd_meta$SR,alpha_divers_pd_meta[[field1]],p.adjust.method="none" )
      print(pairwise.wilcox.test_PD)

      print(ggplot(alpha_divers_pd_meta, aes(x=get(field1), y=log(PD))) +
               ggtitle(paste("FaithPD-phylogenetic diversity (PD) (log)", sheet_a,
                             "\nAnova",eval(field1), anova_PD$aov.tab$`Pr(>F)`[1],
                             "\nTukey sig",paste(rownames(PD_tukey_df_sig), collapse = ";"))) +  
               geom_boxplot()+
               theme_bw() + 
              theme(axis.text.x = element_text(angle = 90),
                    text = element_text(size = 12))+
              stat_compare_means(method = "anova")) #, axis.text.x = element_blank()))
      ggsave(paste0(result_dir,"/FaithPD_PD_",eval(field1),"_boxplot_",proj,"_",set,"_",sheet_a,".jpg"), 
              width=7, height=5)
      dev.off()
      print(ggplot(alpha_divers_pd_meta, aes(x=get(field1), y=log(SR))) +
               ggtitle(paste("FaithPD-species richness (SR) (log)", sheet_a,
                             "\nAnova",eval(field1), anova_SR$aov.tab$`Pr(>F)`[1],
                             "\nTukey sig",paste(rownames(SR_tukey_df_sig), collapse = ";"))) +  
               geom_boxplot()+
               theme_bw() + 
              theme(axis.text.x = element_text(angle = 90),
                    text = element_text(size = 12))+
              stat_compare_means(method = "anova")) #, axis.text.x = element_blank()))
      ggsave(paste0(result_dir,"/FaithPD_SR_",eval(field1),"_boxplot_",proj,"_",set,"_",sheet_a,".jpg"), 
              width=7, height=5)
       dev.off()
     }
#browser()
     writeData(wb_in,sheet_a_anova,anova_PD_call,
               colNames=T, rowNames=T, startRow= 1) 
     addStyle(wb_in, sheet_a_anova, style = titStyle, cols=1:1, rows=1:1, gridExpand = TRUE)
     writeData(wb_in,sheet_a_anova, anova_PD$aov.tab, colNames = T, rowNames= T, startRow= 2) 
     if (length(PD_tukey_df_sig)>0){
       writeData(wb_in,sheet_a_anova,paste(pairwise.wilcox.test_PD$method,pairwise.wilcox.test_PD$data.name),
                 colNames=T, rowNames=T, startRow= 1,startCol= 10) 
       writeData(wb_in,sheet_a_anova, pairwise.wilcox.test_PD$p.value, colNames = T, rowNames= T, 
               startRow= 2, startCol= 10) 
     } else {
       writeData(wb_in,sheet_a_anova,"PD_tukey_df_sig",
                 colNames=T, rowNames=T, startRow= 1,startCol= 10) 
       writeData(wb_in,sheet_a_anova, PD_tukey_df_sig, colNames = T, rowNames= T, 
                 startRow= 2, startCol= 10) 
     }
#browser()
     st_row<-max(12,nrow(PD_tukey_df_sig))+2
     writeData(wb_in,sheet_a_anova,anova_SR_call,
               colNames=T, rowNames=T, startRow= (st_row-1))
     if (length(SR_tukey_df_sig)>0){
       writeData(wb_in,sheet_a_anova,paste(pairwise.wilcox.test_SR$method,pairwise.wilcox.test_SR$data.name),
                colNames=T, rowNames=T, startRow= (st_row-1),startCol= 10) 
       writeData(wb_in,sheet_a_anova, pairwise.wilcox.test_SR$p.value, colNames = T, rowNames= T, 
                 startRow= st_row, startCol= 10) 
     } else {
       writeData(wb_in,sheet_a_anova,"SR_tukey_df_sig",
                 colNames=T, rowNames=T, startRow= (st_row-1),startCol= 10) 
       writeData(wb_in,sheet_a_anova, SR_tukey_df_sig, colNames = T, rowNames= T, 
                 startRow= st_row, startCol= 10) 
     }
     addStyle(wb_in, sheet_a_anova, style = titStyle, cols=1:1, rows=(st_row-1):(st_row-1), gridExpand = TRUE)
     writeData(wb_in,sheet_a_anova, anova_SR$aov.tab, colNames = T, rowNames= T, startRow= st_row) 
 } 
   

  ###------------------------alpha_diversity----------------------------------
   print("alpha_diversity")
   ##Test whether the observed number of ASVs differs significantly between groups
#browser()
   measures <- c("Observed",  "Shannon")

   alpha_diversity = estimate_richness(phy_rarefied,measures=measures) 
   pairwise.wilcox.test_measures <- NULL

   #alpha_diversity$ShannonEvenness = rich$Shannon / log(rich$Observed)
   #rownames(alpha_diversity) = gsub("X", "", rownames(alpha_diversity))
   #writeData(wb_in,sheet_a_divers , alpha_diversity, colNames = TRUE) 
   #setColWidths(wb_in, sheet_a_divers, cols = 1:8, widths = "auto")
   
   #measures <- c("Observed", "Chao1", "ACE", "Shannon", "Simpson", "InvSimpson", "Fisher")
   print(measures)
   if (missing(field2)){
     print(plot_richness(phy_rarefied, x=eval(field1), measures=measures) + 
             xlab(eval(field1))+
             geom_boxplot()+
             ggtitle(paste("Richness box per",eval(field1),set_name))+ 
             geom_point(position = position_dodge(width = 0.5))+
             stat_compare_means(method = "anova"))
     dev.copy(jpeg,filename=paste0(result_dir,"/Alpha_",sheet_a_anova,"_boxplot_",set_name, ".jpg"), 
              width=2000, height=1000)
     dev.off()
   } else{
     print(plot_richness(phy_rarefied, x=eval(field1), measures=measures, color=eval(field2)) + 
             xlab(eval(field1))+
             geom_boxplot()+
             ggtitle(paste("Richness box per",eval(field1),set_name))+ 
             geom_point(position = position_dodge(width = 0.5))+
             stat_compare_means(method = "anova"))
     dev.copy(jpeg,filename=paste0(result_dir,"/Alpha_",sheet_a_anova,"_boxplot_",set_name, ".jpg"), 
              width=2000, height=1000)
     dev.off()
     print(plot_richness(phy_rarefied, x=eval(field2), measures=measures, color=eval(field1)) + 
             xlab(eval(field2))+
             geom_boxplot()+
             ggtitle(paste("Richness box per",eval(field2),set_name))+ 
             geom_point(position = position_dodge(width = 0.5))+
             stat_compare_means(method = "anova"))
     dev.copy(jpeg,filename=paste0(result_dir,"/Alpha_",sheet_a_anova,"_boxplot_",set_name, ".jpg"), 
              width=2000, height=1000)
     dev.off()
     if(!missing(field3)){
       print(field3)
       print(plot_richness(phy_rarefied, x=eval(field1), measures=measures, color=eval(field3)) + 
               xlab(eval(field2))+
               geom_boxplot()+
               ggtitle(paste("Richness box per",eval(field2),set_name))+ 
               geom_point(position = position_dodge(width = 0.5))+
               stat_compare_means(method = "anova"))
       dev.copy(jpeg,filename=paste0(result_dir,"/Alpha_",eval(field1),"_",eval(field3),"_boxplot_",set_name, ".jpg"), 
                width=2000, height=1000)
       dev.off()
       print(plot_richness(phy_rarefied, x=eval(field2), measures=measures, color=eval(field3)) + 
               xlab(eval(field2))+
               geom_boxplot()+
               ggtitle(paste("Richness box per",eval(field2),set_name))+ 
               geom_point(position = position_dodge(width = 0.5))+
               stat_compare_means(method = "anova"))
       dev.copy(jpeg,filename=paste0(result_dir,"/Alpha_",sheet_a_anova,"_boxplot_",set_name, ".jpg"), 
                width=2000, height=1000)
       dev.off()
     }
   }
   st_row<-max(12,(nrow(PD_tukey_df_sig)+nrow(SR_tukey_df_sig)))+2
   md<-sample_data(phy_rarefied)
   alpha_divers_meta<- alpha_diversity %>%
                         rownames_to_column("rowname") %>%
                         mutate(rowname=str_replace_all(rowname,"\\.","\\-")) %>%
                         mutate(rowname=str_replace(rowname,"^X","")) %>%
                         left_join(rownames_to_column(meta_16S,"rowname"),by="rowname") %>%
                         column_to_rownames("rowname")
   print("measures loop")
#   wilcox <- data.frame(measure=character(),field1_pv=double(), field2_pv=double())
   for (m in measures){
     #m<-"Shannon"
#     wilcox_field1 <- pairwise.wilcox.test(alpha_divers_meta[[m]], alpha_divers_meta[[field1]] )

     if(missing(field2)) {
       alpha_divers_meta_m<- subset(alpha_divers_meta,select=(c(get(m),get(field1)))) %>%
                               dplyr::filter( str_length(get(field1))>0  & !is.na(get(field1))) #%>%
       #anova <- adonis(formula =alpha_diversity[[m]] ~ md[[field1]])
       anova <- adonis(formula =alpha_divers_meta_m[[m]] ~ get(field1), data=alpha_divers_meta_m)
       anova_call<-paste(m,"~",field1)
       pairwise.wilcox.test_measures <- pairwise.wilcox.test(alpha_diversity[[m]],md[[field1]],p.adjust.method="none" )
     }else{
        if(missing(field3)) {
           alpha_divers_meta_m<- subset(alpha_divers_meta,select=(c(get(m),get(field1),get(field2)))) %>%
                                    dplyr::filter( str_length(get(field1))>0  & !is.na(get(field1)) & str_length(get(field2))>0  & !is.na(get(field2))) #%>%
#browser()
           
           anova <- adonis(formula =alpha_divers_meta_m[[m]] ~ get(field1)*get(field2), data=alpha_divers_meta_m)
           #anova <- adonis(formula =alpha_diversity[[m]] ~ md[[field1]]*md[[field2]])
           anova_call<-paste(m,"~",field1,"*",field2)
       } else {
         alpha_divers_meta_m<- subset(alpha_divers_meta,select=(c(get(m),get(field1),get(field2),get(field3))))%>%
           dplyr::filter( str_length(get(field1))>0  & !is.na(get(field1)) & str_length(get(field2))>0  & !is.na(get(field2)) & str_length(get(field3))>0  & !is.na(get(field3))) #%>%
         if (plus_f3){
           anova <- adonis(formula =get(m) ~ get(field1)*get(field2)+get(field3), data=alpha_divers_meta_m)
           #anova <- adonis(formula =alpha_diversity[[m]] ~ md[[field1]]*md[[field2]]+md[[field3]])
           anova_call<-paste(m,"~",field1,"*",field2,"+",field3)
         } else {
           anova <- adonis(formula =alpha_divers_meta_m[[m]] ~ get(field1)*get(field2)*get(field3), data=alpha_divers_meta_m)
           #anova <- adonis(formula =alpha_diversity[[m]] ~ md[[field1]]*md[[field2]]*md[[field3]])
           anova_call<-paste(m,"~",field1,"*",field2,"*",field3)
         }
       }
     }
      anova
#     wilcox <- rbind(wilcox, cbind(Mmeasure = m, 
#                                   field1_pv= as.double(wilcox_field1$p.value), 
#                                   field2_pv= as.double(wilcox_field2$p.value)
     
     writeData(wb_in,sheet_a_anova,anova_call,
               colNames=T, rowNames=T, startRow= st_row) 
     addStyle(wb_in, sheet_a_anova, style = titStyle, cols=1:1, rows=st_row:st_row, gridExpand = TRUE)
     writeData(wb_in,sheet_a_anova,paste(pairwise.wilcox.test_measures$method,pairwise.wilcox.test_measures$data.name),
               colNames=T, rowNames=T, startRow= st_row,startCol= 10) 
     writeData(wb_in,sheet_a_anova, anova$aov.tab, colNames = T, rowNames= T, startRow= st_row+1) 
     writeData(wb_in,sheet_a_anova, pairwise.wilcox.test_measures$p.value, colNames = T, rowNames= T, 
               startRow= st_row, startCol= 10) 
     st_row<-st_row+10
     
   }
#   wilcox<-wilcox %>%
#     data.table::setnames(old=c("field1_pv","field2_pv"),
#                          new=c(paste0("pv_",eval(field1)),paste0("pv_",eval(field2))) )
   
#   addStyle(wb_in, sheet_wilcox, style = s, cols=2:5, rows=2:nrow(wilcox), gridExpand = TRUE)
   ## set a default number format for numeric columns of data.frames
#   options("openxlsx.numFmt" = "0.00")
#   writeData(wb_in,sheet_wilcox , "pairwise.wilcox.test" , colNames = TRUE) 
#   writeData(wb_in,sheet_wilcox , wilcox, colNames = TRUE,startRow = 2) 
#   conditionalFormatting(wb_in, sheet_wilcox, cols=2:5, rows=2:nrow(wilcox),rule="B2<0.05", style = posStyle)
#   setColWidths(wb_in, sheet_wilcox, cols = 1:8, widths = "auto")
 }

 
 Bdiver<- function(wb_in=wb, phy_rarefied,field1,field2,field3, plus_f3=T,Bdiver_dist=T,anova,
                   set_name="all",   dst= "unifrac" ,weight=T ,sheet_b= NULL){
   #phy_rarefied<-physeq_rarefied_MW
   #field1<- "Condition" #RatType TimeW Condition
   #field2<- "Wasp" #TimeGroup PatientID Wasp
   #field3<- "Stage" #RatID batch Stage
   # plus_f3=F
   #dst<- "unifrac" #bray unifrac
   set.seed(42)
   
   sheet_b_divers <- paste(set_name,sheet_b)
   addWorksheet(wb_in, sheet_b_divers)
   st_row<-1
   print(sheet_b_divers)
   titStyle <- createStyle(fontColour = "#006100", fgFill = "#C6EFCE")
   
   # PCoA plot using the weighted/un UniFrac/bray as distance
   theme_set(theme_bw())
   # methods for distance
   #str(phy_tree(phy_rarefied))
#browser()
   phy_tree(phy_rarefied) <- ape::multi2di(phy_tree(phy_rarefied))
   #dist_methods <- c("bray")
   ##  distances between samples
   if (missing(anova))     dist <- phyloseq::distance(phy_rarefied, method=dst, weighted=weight) #
   
   # ANOVA-Test whether the groups, Condition,  differ significantly from each
   if (missing(field2)){
#browser()

     if (missing(anova))     anova <- adonis2(dist ~   sample_data(phy_rarefied)[[field1]])
     anova_call<-paste(sheet_b_divers,": dist ~ ",field1)
     dist_df<- as.data.frame(melt(as.matrix(dist))) %>%
                left_join(rownames_to_column(dplyr::select(meta_16S,field1),"Var1"), by="Var1") %>%
                left_join(rownames_to_column(dplyr::select(meta_16S,field1),"Var2"), by="Var2") %>%
                mutate(cond1=get(paste0(field1,".x"))) %>%
                mutate(cond2=get(paste0(field1,".y"))) 
   }  else if (missing(field3)){
#browser()     
     if (missing(anova))    anova <- adonis2(dist ~ sample_data(phy_rarefied)[[field1]] *
                                                    sample_data(phy_rarefied)[[field2]] )
     anova_call<-paste(sheet_b_divers,": dist ~ ",field1,"*",field2)
     dist_df<- as.data.frame(melt(as.matrix(dist))) %>%
       left_join(rownames_to_column(dplyr::select(meta_16S,field1,field2),"Var1"), by="Var1") %>%
       left_join(rownames_to_column(dplyr::select(meta_16S,field1,field2),"Var2"), by="Var2") %>%
       mutate(cond1=paste0(get(paste0(field1,".x")),"_",get(paste0(field2,".x")))) %>%
       mutate(cond2=paste0(get(paste0(field1,".y")),"_",get(paste0(field2,".y")))) 
   } else {
     if (plus_f3){
       print("plus_f3")
       if (missing(anova))    anova <- adonis2(dist ~ sample_data(phy_rarefied)[[field1]] *
                                         sample_data(phy_rarefied)[[field2]] +
                                         sample_data(phy_rarefied)[[field3]])
       anova_call<-paste(sheet_b_divers,": dist ~ ",field1,"*",field2,"+",field3)
     } else {
       if (missing(anova))    anova <- adonis2(dist ~ sample_data(phy_rarefied)[[field1]] *
                                         sample_data(phy_rarefied)[[field2]] *
                                         sample_data(phy_rarefied)[[field3]])
       anova_call<-paste(sheet_b_divers,": dist ~ ",field1,"*",field2,"*",field3)
      }
     dist_df<- as.data.frame(melt(as.matrix(dist))) %>%
       left_join(rownames_to_column(dplyr::select(meta_16S,field1,field2,field3),"Var1"), by="Var1") %>%
       left_join(rownames_to_column(dplyr::select(meta_16S,field1,field2,field3),"Var2"), by="Var2") %>%
       mutate(cond1=paste0(get(paste0(field1,".x")),"_",get(paste0(field2,".x")),"_",get(paste0(field3,".x")))) %>%
       mutate(cond2=paste0(get(paste0(field1,".y")),"_",get(paste0(field2,".y")),"_",get(paste0(field3,".y")))) 
   }
   print(anova)
 #browser()  
   writeData(wb_in,sheet_b_divers,startRow = st_row,
             #rownames_to_column(anova$aov.tab,anova_call),
             rownames_to_column(as.data.frame(anova),anova_call),
             colNames = TRUE) 
   
   # beta diversity boxplots-distribution of distances
   st_row<-st_row+8

   anova3_2way <- aov(value ~  cond1+cond2, data=dist_df) #
   tukey<- TukeyHSD(anova3_2way)
   tukey_df<-as.data.frame(tukey$cond2)
   tukey_df_sig<-subset(tukey_df,tukey_df$`p adj` <0.05)
   print(ggplot(dist_df, aes(x=as.factor(cond1), y=value, color=as.factor(cond2))) +
           geom_boxplot()+
           xlab(eval(field1)) +
           theme_bw()+
           #geom_jitter(shape=16, position=position_jitter(0.2))+
           theme(text = element_text(size = 12),axis.text.x = element_text(angle = 45, hjust = 1))+
           ggtitle(paste(sheet_b_divers, "Group significance plot-",dst,"distance ",
                        # "\nAnova2way",anova2$aov.tab$`Pr(>F)`[1],
                         "\n sig tukey:",paste(rownames(tukey_df_sig), collapse = ";")))+
           stat_compare_means(method = "anova"))
   ggsave(paste0(result_dir,"/boxplot_dist_",dst,"_",proj,"_",set,"_",sheet_b_divers,".jpg"), 
          width=7, height=5)
   dev.off()
   
   if (Bdiver_dist){
     for (cnd in unique(dist_df$cond1)){
       #cnd<-"G_MW_CTRL-G"
       dist_cnd<-subset(dist_df,cond1==cnd ) #& cond1!=cond2
       #deal with zeros
       dist_cnd$value<-sqrt (1+dist_cnd$value)
       
       anova2 <- adonis(value ~ cond2, data = dist_cnd)
       anova_2way <- aov(value ~  cond2, data=dist_cnd) #
       tukey<- TukeyHSD(anova_2way)
       tukey_df<-as.data.frame(tukey$cond2)
       tukey_df_sig<-subset(tukey_df,tukey_df$`p adj` <0.05)
       
       writeData(wb_in,sheet_b_divers,paste(cnd,as.character(anova2$call[2])),
                 colNames=T, rowNames=T, startRow= st_row) 
       addStyle(wb_in, sheet_b_divers, style = titStyle, cols=1:1, rows=st_row:st_row, gridExpand = TRUE)
       writeData(wb_in,sheet_b_divers, anova2$aov.tab, colNames = T, rowNames= T, startRow= st_row+1) 
       writeData(wb_in,sheet_b_divers, paste(rownames(tukey_df_sig), collapse = ";  "),
                 colNames = T, rowNames= T, startRow= st_row+5) 
       st_row<-st_row+8
       
     }
   }
   setColWidths(wb_in, sheet_b_divers, cols = 1:10, widths = "auto")
   
   #plot 
   ord_meths = c( "CCA", "RDA", "DPCoA", "NMDS", "PCoA") #"DCA", "MDS",
   #ord_meths =  "CCA"
   
   for (mth in ord_meths){
     #  for (dst in dist_methods){
     #mth<- "PCoA"
     
     print(paste("method:", mth, "dist method: ", dst))
#browser() 
     if (mth!="DPCoA") ord <- ordinate(phy_rarefied, method=mth, distance=dst, weighted=weight) else ord <- ordinate(phy_rarefied, method=mth, distance=dst)
     
     #plots
     result = tryCatch({
       if (!missing(field2)){
         print(plot_ordination(phy_rarefied, ord, color=eval(field1), shape = eval(field2)) + #, label="SampleID"
               theme(aspect.ratio=1,plot.title = element_text(size = 16, face = "bold")) +
               geom_point(size = 5) +
               ggtitle(paste( dst, "Distance Method,",mth,"Ordination Method",sheet_b_divers))+ 
               #scale_shape_manual(values = c(20,18))+
               stat_ellipse() +coord_fixed())#+ #geom = "circle"
       #        geom_path() +
       
       dev.copy(jpeg,filename=paste0(result_dir,"/Beta_",mth,"_",sheet_b_divers, "_plot.jpg"), 
                width=600, height=600)
       dev.off()
       } else{
         print("only field1")
         print(plot_ordination(phy_rarefied, ord, color=eval(field1)) + #, label="SampleID"
                 theme(aspect.ratio=1,plot.title = element_text(size = 16, face = "bold")) +
                 geom_point(size = 5) +
                 ggtitle(paste( dst, "Distance Method,",mth,"Ordination Method",sheet_b_divers))+ 
                 scale_shape_manual(values = c(20,18))+
                 stat_ellipse() +coord_fixed())#+ #geom = "circle"
         dev.copy(jpeg,filename=paste0(result_dir,"/Beta_",mth,"_",eval(field1),"_",sheet_b_divers, "_plot.jpg"), 
                  width=600, height=600)
         dev.off()
       }
       if (!missing(field3)){
         print(plot_ordination(physeq_rarefied, ord, color=eval(field1), shape = eval(field3)) + #, label="SampleID"
                 theme(aspect.ratio=1,plot.title = element_text(size = 16, face = "bold")) +
                 geom_point(size = 5) +
                 ggtitle(paste( dst, "Distance Method,",mth,"Ordination Method",sheet_b_divers))+ 
                 scale_shape_manual(values = c(20,18))+
                 stat_ellipse() +coord_fixed())#+ #geom = "circle"
         #        geom_path() +
         dev.copy(jpeg,filename=paste0(result_dir,"/Beta_",mth,"_",sheet_b_divers, "_plot.jpg"), 
                  width=600, height=600)
         dev.off()
       }
     },
     error = function (condition) {
       print(paste("ERROR in method:", mth, "dist method: ", dst))
     })
   }
   sig_anova<-as.data.frame(subset(anova,`Pr(>F)`<0.05))
   print(paste("no of sig anova:",nrow(sig_anova)))
   if (nrow(sig_anova)>0){return(TRUE)}else{return(FALSE)}
 }
 
 
 deseq_part<- function(md,taxa_count,field,value, to_inc=T,dds_design,res_for,res_for2,set_name,exact=F,
                       res_dir=result_dir, element="taxa",to_label=T,filter_id=F){
   #value  <-"GDM"   # "CTRL-G"
   #field<-"Control_GDM"   #Condition"
   #dds_design<-" ~ woman + trimester"   #formula(" ~ Stage") 
   #res_for<-"trimester"   #Stage"
   #set_name<-GDM_only
#browser()   
   
   if (field!="dummy"){
     if (to_inc)  {
       if (!exact) md_part<-md %>%
                   dplyr::filter(str_detect(get(field),value) & str_length(md[[res_for]])>0  & !is.na(get(res_for)) & get(res_for)!="na" & get(res_for)!="NA") %>%
                   dplyr::select(-eval(field)) 
        else md_part<-md %>%
            dplyr::filter(get(field)==value & str_length(md[[res_for]])>0  & !is.na(get(res_for)) & get(res_for)!="na" & get(res_for)!="NA") %>%
            dplyr::select(-eval(field))
     } else {
       md_part<-md %>%
                   dplyr::filter(!str_detect(get(field),value)) 
       md_part[[field]]<-factor(md_part[[field]],levels = unique(md_part[[field]]))
     }
   }else{
     md_part<-md #%>%
   }
   md_part[[res_for]]<-factor(md_part[[res_for]],levels = unique(md_part[[res_for]]))
#browser()   
   if (missing(res_for2)){
     print(table(md_part[[res_for]]))
   } else {
     print(table(md_part[[res_for]],md_part[[res_for2]]))
   }
   
   assign(paste0("md_",set_name), md_part, envir = .GlobalEnv)
   taxa_count_part<-taxa_count %>%
     dplyr::select(rownames(md_part)) #%>% # filter zero rows
   #select_if(purrr::negate(function(col) is.numeric(col) && sum(col) >0))
#browser()   
   #sum the abundnce per taxa
   assign(paste0(element,"_count_",set_name), taxa_count_part, envir = .GlobalEnv)

   dds_taxa_part <- DESeqDataSetFromMatrix(countData = taxa_count_part,
                                         colData = md_part,
                                         design=dds_design)
   
   dds_taxa_part = DESeq(dds_taxa_part)
   print(resultsNames(dds_taxa_part))
   assign("dds_count_df",as.data.frame(counts(dds_taxa_part, normalized=TRUE)), envir = .GlobalEnv)
   assign(paste0(element,"_norm_abund_",set_name), dds_count_df, envir = .GlobalEnv)
   
   
#browser()
   list1<-rev(levels(md_part[[res_for]]))
   print(list1)
   list2<-list1
   res_all<-character()
   res_all_df<-character()
   for (r1 in list1){
     list2<-list2[list2 != r1]
     for (r2 in list2){
       if (r1!=r2){
         #r1<-"CTRL-G"; r2<-"CTRL-O"
         #The factor level given last is the base level for the comparison.
         #print(paste("res_for,r1,r2",res_for,r1,r2))
#browser()
         taxa_res_part<-results(dds_taxa_part,contrast=c(res_for,r1,r2))
         if (filter_id) taxa_res_part<-subset(taxa_res_part,!startsWith(rownames(taxa_res_part),"id_"))
         DESeq2::summary(taxa_res_part)
         md_part2<-md_part %>%
                  dplyr::filter(get(res_for)==r1 | get(res_for)==r2)
#browser()
         vst_part<-varianceStabilizingTransformation (dds_taxa_part, blind=FALSE) %>%
                     assay() %>%
                     as.data.frame(stringsAsFactors = FALSE) %>%
                     #rownames_to_column %>%
                     #mutate(rowname=substr(rowname,1,75)) %>%
                     #column_to_rownames %>%
                     dplyr::select(rownames(md_part2))
         if (to_inc) {name<-paste0("_",value,"_",r1,"_vs_",r2)} else{
          name<-paste0("_exclude-",value,"_",r1,"_vs_",r2)
         }
         assign(paste0(element,name),taxa_res_part, envir = .GlobalEnv)
         assign(paste0("vst",name), vst_part, envir = .GlobalEnv)
         assign(paste0("anno",name), subset(md_part2,select=eval(res_for)), envir = .GlobalEnv)
         res_all<-c(res_all,paste0(element,name))
         print(paste("res_all:",res_all))
         taxa_res<-as.data.frame(taxa_res_part) %>%
                    rownames_to_column("Gene_name") %>%
                    mutate(Comparison=name)
         print(name)
         res_all_df<-rbind(res_all_df,taxa_res)
       }
     }
   }

   boxplot(log10(assays(dds_taxa_part)[["cooks"]]), range=0, las=2,
           main=paste("boxplot",element," Cook’s distances - test for outliers",
           "\n",set_name),
           ylab="Cook’s distances")
   dev.copy(jpeg,filename=paste0(res_dir,"/plot_",element,"_Cooks_distances_",set_name, ".jpg"), 
            width=600, height=600)
   dev.off()
   
   pc<- plot_PCA(dds_taxa_part,group=res_for, set_name=paste0("res_",element,"_",set_name,"_for_",res_for),
                 dds_res=DESeq2::summary(taxa_res_part), res_dir = res_dir,to_label = to_label)
   ph<- plot_dds_heatmap(dds_taxa_part,group=res_for, res_dir= res_dir,
                         set_name=paste0("res_",element,"_",set_name,"_for_",res_for))
   ## Bray-Curtis distances between samples
   norm_count<-as.data.frame(counts(dds_taxa_part, normalized=TRUE))
   openxlsx::write.xlsx(norm_count,paste0(res_dir,"/Norm_count_",element,"_",set_name,".xlsx"),row.names=T)
   bd<- Bray_dist(norm_count,md_part,res_for,set_name = set_name, res_dir= res_dir)
   
   
   ## Bray-Curtis distances between samples
   #bd<- Bdiver(taxa_count_MW,md_MW,"Wasp")
   return(res_all)
 }
 
 
 prep_count_levels<- function(phy_rarefied_data,taxa_level=c("short_title", "Family", "Genus")){
   #phy_rarefied_data<-physeq_rarefied_data                      
   
   for (tl in taxa_level){
     #tl<-"short_title"
     print(tl)
     taxa_count<-phy_rarefied_data %>%
                   dplyr::select(Sample,Abundance,eval(tl)) %>%
                   # only samples that has RNAseq
                   dplyr::filter(Sample %in% rownames(md)) %>%
                   dplyr::group_by(Sample,get(tl)) %>% 
                   dplyr::summarise(total_abund = sum(Abundance))  %>%
                   pivot_wider(names_from =Sample, values_from =total_abund) %>%
                   dplyr::rename("taxa"='get(tl)') %>%
                   dplyr::filter(taxa!="k__Bacteria" & 
                                   taxa!="d__Bacteria" & 
                                   str_length(taxa)>0 &
                                   !is.na(taxa)) %>%
                   replace(is.na(.), 0) %>% 
                   column_to_rownames("taxa")
    # filter zero rows
     taxa_count <- taxa_count[, colSums(taxa_count != 0) > 0]
     assign(paste0("taxa_count_",eval(tl)),taxa_count,envir = .GlobalEnv)
   }
   return(taxa_level)
   
 } 
 

 
 metaphlanToPhyloseq <- function(
   tax,
   metadat=NULL,
   simplenames=TRUE,
   roundtointeger=FALSE,
   split="|"){
   ## tax is a matrix or data.frame with the table of taxonomic abundances, rows are taxa, columns are samples
   ## metadat is an optional data.frame of specimen metadata, rows are samples, columns are variables
   ## if simplenames=TRUE, use only the most detailed level of taxa names in the final object
   ## if roundtointeger=TRUE, values will be rounded to the nearest integer
   xnames = rownames(tax)
   shortnames = gsub(paste0(".+\\", split), "", xnames)
   if(simplenames){
     rownames(tax) = shortnames
   }
   if(roundtointeger){
     tax = round(tax * 1e4)
   }
   x2 = strsplit(xnames, split=split, fixed=TRUE)
   taxmat = matrix(NA, ncol=max(sapply(x2, length)), nrow=length(x2))
   colnames(taxmat) = c("Kingdom", "Phylum", "Class", "Order", "Family", "Genus", "Species", "Strain")[1:ncol(taxmat)]
   rownames(taxmat) = rownames(tax)
   for (i in 1:nrow(taxmat)){
     taxmat[i, 1:length(x2[[i]])] <- x2[[i]]
   }
   #taxmat = gsub("[a-z]__", "", taxmat)
   #taxmat = phyloseq::tax_table(taxmat)
   #otutab = phyloseq::otu_table(tax, taxa_are_rows=TRUE)
   colnames(tax)[match(metadat$BarcodeID, str_split_fixed(colnames(tax),"_",3)[,2])] = rownames(metadat)
   assign("Taxa" , taxmat,envir = .GlobalEnv)
   
#browser()
   TAX = tax_table(as.matrix(taxmat))
   OTU = otu_table(as.matrix(tax), taxa_are_rows = TRUE)
   META <- sample_data(metadat)
   head(sample_names(OTU))
   head(sample_names(META))
   
   if(is.null(metadat)){
     res = phyloseq::phyloseq(taxmat, otutab)
   }else{
     res = phyloseq::phyloseq(TAX, OTU, META)
   }
   return(res)
 }
```
