## Supplementary material for "Barking up the microbiome tree: a multi-body-site characterization of maternal postpartum and pup early colonization dynamics": Code: Puppies_2026_publication.Rmd

````
---
title: "Puppies_2025"
author: "Tali Magory"
date: "2025-04-28"
output: html_document
---

Clear the environment

```{r}
rm(list = ls())
```

# Load libraries

```{r, include=FALSE}
library(phyloseq)
library(ggplot2)
library("dplyr")
library("tidyr")
library("tibble")
library("RColorBrewer")
library("pheatmap")
library(ape)
library(stringr)
library(vegan)
library(btools)
library(ggpubr)
library(microbiome)
library(Maaslin2)
library(openxlsx)
library(rstatix)
library(biomformat)
library(Biostrings)
library(FEAST)
library(forcats)

source("Phyloseq_functions.R")

```

# Define variables

```{r}
pwd <- getwd()
results_dir <- file.path(pwd, "results")

# Define a consistent color palette

TypeHostNew_colors <- c(
  "HealthyPupRectal1" = "cornflowerblue",
  "HealthyPupRectal8" = "blue2",
  "HealthyPupOral1" = "gold1",
  "HealthyPupOral8" = "darkgoldenrod4",
  "DamMilk1" = "#66C2A5",
  "DamMilk8" = "#117733",
  "DamRectal1" = "cornflowerblue",
  "DamRectal8" = "blue2",
  "DamVaginal1" = "mediumorchid",
  "DamVaginal8" = "darkorchid4",
  "DamOral1" = "gold1",
  "DamOral8" = "darkgoldenrod4"
)

TypeHostNew_shapes <- c(
  "HealthyPupRectal1" = 21,
  "HealthyPupRectal8" = 21,
  "HealthyPupOral1" = 21,
  "HealthyPupOral8" = 21,
  "DamMilk1" = 16,
  "DamMilk8" = 16,
  "DamRectal1" = 16,
  "DamRectal8" = 16,
  "DamVaginal1" = 16,
  "DamVaginal8" = 16,
  "DamOral1" = 16,
  "DamOral8" = 16
)

```

# Load data

```{r}
# merged ok175 and ok180
otu_mat <- read.table("otu_2025.tsv", header = TRUE, sep = "\t", stringsAsFactors = FALSE)
tax_mat <- read.table("taxonomy_2025.tsv", header = TRUE, sep = "\t", stringsAsFactors = FALSE)
samples_df <- read.csv("metadata_merge_new.csv")
tree <- read.tree("tree_2025.nwk")
seqs <- readDNAStringSet("aligned-dna-sequences.fasta")

```

# Prepare data

otu_mat

```{r}
# Remove the leading 'X' from the column names that start with it
colnames(otu_mat) <- gsub("^X", "", colnames(otu_mat))

# Make the column names the row names
otu_mat1 <-  otu_mat %>% column_to_rownames("OTU.ID")
as.data.frame(otu_mat1)

```

tax_mat

```{r}
# split taxon column and create short_title, and get the best from 2 classifiers
taxa_16S_df<-prep_taxa(tax_mat) 
nrow(taxa_16S_df)

# Set the row names to be from the "Feature.ID" column
  rownames(taxa_16S_df) <- tax_mat$Feature.ID

# Define the values to remove
remove_values <- c("s__", "f__", "c__", "o__", "g__", "p__")

# Define the columns to process
columns_to_process <- c("Phylum", "Class", "Order", "Family", "Genus", "Species")

# Create a copy of the data frame to modify
taxa_16S_clean <- taxa_16S_df

# Loop through each column and replace matching values with NA
for (col in columns_to_process) {
  taxa_16S_clean[[col]] <- ifelse(
    str_trim(taxa_16S_clean[[col]]) %in% remove_values, 
    "", 
    str_trim(taxa_16S_clean[[col]])
  )
}
```

metadata

```{r}
# make samples the rownames
samples_df <- samples_df %>% 
    tibble::column_to_rownames("SampleID") 
```

## Make a phyloseq object

```{r}
OTU <- otu_table(otu_mat1, taxa_are_rows = TRUE)
TAX <- tax_table(as.matrix(taxa_16S_clean))
SAMPLE <- sample_data(samples_df)
TREE <- read_tree(tree)

```

Check if the taxa/OTU names match

```{r}
# Check OTU names in the OTU table
otu_names <- taxa_names(OTU)

# Check OTU names in the taxonomic table
tax_names <- taxa_names(TAX)

# Compare
identical(otu_names, tax_names)  # Should return TRUE
```

Check if sample names match

```{r}
otu_s_names <- sample_names(OTU_175)
sample_names <- sample_names(SAMPLE_175)
```

make a phyloseq object

```{r}
physeq <- phyloseq(OTU,TAX, SAMPLE, TREE)
physeq
```

Resolve tree topology

```{r}
data_in <- physeq

# Check if binary (dichotomy) & multifurcating (polytomy) trees
sprintf("Is tree binary: %s", is.binary(TREE))
#> [1] "Is tree binary: FALSE"

# If FASLE, randomly resolve polytomies and replace tree in "ps"
phy_tree(physeq) = multi2di(TREE)
sprintf("Is tree binary: %s", is.binary(phy_tree(physeq)))
#> [1] "Is tree binary: TRUE"
```

## Subsetting

```{r}
# Define the samples 
neg_list <- c("Kit3", "MOCK1", "MOCK2")

non_study <- c("Kit3", "MOCK1", "MOCK2")

# Subset the phyloseq object to exclude negative control samples
phy_noneg <- subset_samples(physeq, !(sample_names(physeq) %in% neg_list))

# Subset the phyloseq object to exclude negative and positive control samples
phy_study <- subset_samples(physeq, 
                                 !(sample_names(physeq) %in% non_study) & !(sample_data(physeq)$Host == "FadingPup"))
```

return OTU names to phyloseq object

```{r}
otu_names <- rownames(OTU)
taxa_names(phy_study) <- paste(otu_names)
taxa_names(phy_noneg) <- paste(otu_names)
```

Count samples

```{r}
df_input_metadata <- as(sample_data(phy_study), "data.frame")
count_table <- df_input_metadata  %>%
    dplyr::count(Type, Host)
print(count_table)
```

Subset according to group we are testing

```{r}
# Only mothers - milk
milk <- subset_samples(phy_study, Type =="Milk")

# Only mothers - vaginal
vaginal <- subset_samples(phy_study, Type =="Vaginal")

# Only mothers - rectal
dams_rectal <- subset_samples(phy_study, Host == "Dam" & Type == "Rectal")

# Only pups - rectal
pups_rectal <- subset_samples(phy_study, Host %in% c("HealthyPup") & Type == c( "Rectal"))

# Only mothers - oral
dams_oral <- subset_samples(phy_study, Host == "Dam" & Type == "Oral")

# Only pups - oral
pups_oral <- subset_samples(phy_study, Host %in% c("HealthyPup") & Type == c( "Oral"))

# All pups
pups_all <- subset_samples(phy_study, Host %in% c("HealthyPup") )

# All dams
dams_all <- subset_samples(phy_study, Host == "Dam")

# All rectal
rectal_all <- subset_samples(phy_study, Type =="Rectal")

# All oral
oral_all <- subset_samples(phy_study, Type =="Oral")

# All Day 1
all_day1 <- subset_samples(phy_study, Day =="1")

# All Day 8
all_day8 <- subset_samples(phy_study, Day =="8")

# Oral & rectal Day 1 pups & dams
oral_rectal_day1 <- subset_samples(phy_study, Day == "1" & Type == c("Rectal","Oral"))

# Oral & rectal Day 8 pups & dams
oral_rectal_day8 <- subset_samples(phy_study, Day == "8" & Type == c("Rectal","Oral"))

# Oral Day 1 pups & dams
oral_day1 <- subset_samples(phy_study, Day == "1" & Type == c("Oral"))

# Oral Day 8 pups & dams
oral_day8 <- subset_samples(phy_study, Day == "8" & Type == c("Oral"))

# Rectal Day 1  & dams
rectal_day1 <- subset_samples(phy_study, Day == "1" & Type == c("Rectal"))

# Rectal Day 8  & dams
rectal_day8 <- subset_samples(phy_study, Day == "8" & Type == c("Rectal"))

# Oral Day 1 pups
pup_oral_day1 <- subset_samples(phy_study, Day == "1" & Type == c("Oral") & Host %in% c("HealthyPup"))

# Oral Day 8 pups
pup_oral_day8 <- subset_samples(phy_study, Day == "8" & Type == c("Oral") & Host %in% c("HealthyPup"))

# Rectal Day 1 pups
pup_rectal_day1 <- subset_samples(phy_study, Day == "1" & Type == c("Rectal") & Host %in% c("HealthyPup"))

# Rectal Day 8 pups
pup_rectal_day8 <- subset_samples(phy_study, Day == "8" & Type == c("Rectal") & Host %in% c("HealthyPup"))

# Oral Day 1 dams
dam_oral_day1 <- subset_samples(phy_study, Day == "1" & Type == c("Oral") & Host %in% c("Dam"))

# Oral Day 8 dams
dam_oral_day8 <- subset_samples(phy_study, Day == "8" & Type == c("Oral") & Host %in% c("Dam"))

# Rectal Day 1 dams
dam_rectal_day1 <- subset_samples(phy_study, Day == "1" & Type == c("Rectal") & Host %in% c("Dam"))

# Rectal Day 8 dams
dam_rectal_day8 <- subset_samples(phy_study, Day == "8" & Type == c("Rectal") & Host %in% c("Dam"))

# Milk Day 1 dams
dam_milk_day1 <- subset_samples(phy_study, Day == "1" & Type == c("Milk") & Host %in% c("Dam"))

# Milk Day 8 dams
dam_milk_day8 <- subset_samples(phy_study, Day == "8" & Type == c("Milk") & Host %in% c("Dam"))

# Vaginal Day 1 dams
dam_vaginal_day1 <- subset_samples(phy_study, Day == "1" & Type == c("Vaginal") & Host %in% c("Dam"))

# Vaginal Day 8 dams
dam_vaginal_day8 <- subset_samples(phy_study, Day == "8" & Type == c("Vaginal") & Host %in% c("Dam"))

datasets <- list(
  ORAL = oral_all,
  RECTAL = rectal_all,
  VAGINAL = vaginal,
  MILK = milk,
  PUPS = pups_all,
  DAMS = dams_all,
  PUPS_oral= pups_oral,
  PUPS_rectal = pups_rectal,
  DAMS_oral = dams_oral,
  DAMS_rectal = dams_rectal,
  ALL = phy_study,
  ALL_day1 = all_day1,
  ALL_day8 = all_day8,
  O_R_day1 = oral_rectal_day1,
  O_R_day8 = oral_rectal_day8,
  ORAL_day1 = oral_day1,
  ORAL_day8 = oral_day8,
  RECTAL_day1 = rectal_day1,
  RECTAL_day8 = rectal_day8,
  PUP_ORAL_day1 = pup_oral_day1,
  PUP_ORAL_day8 = pup_oral_day8,
  PUP_RECTAL_day1 = pup_rectal_day1,
  PUP_RECTAL_day8 = pup_rectal_day8,
  DAM_ORAL_day1 = dam_oral_day1,
  DAM_ORAL_day8 = dam_oral_day8,
  DAM_RECTAL_day1 = dam_rectal_day1,
  DAM_RECTAL_day8 = dam_rectal_day8,
  DAM_MILK_day1 = dam_milk_day1,
  DAM_MILK_day8 = dam_milk_day8,
  DAM_VAGINAL_day1 = dam_vaginal_day1,
  DAM_VAGINAL_day8 = dam_vaginal_day8)

# Create workbook
wb <- createWorkbook()

# Loop over datasets
for (name in names(datasets)) {
  phy_nonfiltered <- datasets[[name]]
  
  # Create metadata df with total read counts
  metadata_nonfiltered <- data.frame(sample_data(phy_nonfiltered))
  metadata_nonfiltered$total_reads <- sample_sums(phy_nonfiltered)
  
  # Add as a sheet in Excel workbook
  addWorksheet(wb, sheetName = name)
  writeData(wb, sheet = name, x = metadata_nonfiltered)
  
}

# Save Excel workbook
saveWorkbook(wb, file = paste0(results_dir, "tables/nonfiltered_metadata_with_total_reads.xlsx"), overwrite = TRUE)

```

## Filtering

Choose the minimum number of reads based on rarefaction curve per dataset (and count tables)

```{r}
# Loop over each dataset
for (name in names(datasets)) {
  obj <- datasets[[name]]
  
  # Calculate summary statistics for read counts
  read_counts <- sample_sums(obj)
  read_counts_df <- as.data.frame(read_counts)
  write.csv(read_counts_df, paste0(results_dir, "tables/rarefaction/rarefaction_", name,".csv"))
  
  # Plot rarefaction curves
  tab <- otu_table(obj)
  class(tab) <- "matrix"
  tab <- t(tab)
  
  # --- Rarefaction curve ---
  jpeg(filename = paste0(results_dir,"plots/rarefaction/Rarefaction_", name, ".jpeg"), width = 1200, height = 800)
  rarecurve(tab, step = 50, lwd = 2, ylab = "OTU", label = TRUE,
            main = paste("Rarefaction Curve:", name))
  dev.off()

  # --- Histogram of read counts ---
  jpeg(filename = paste0(results_dir,"plots/rarefaction/ReadCounts_Histogram_", name, ".jpeg"), width = 1200, height = 800)
  hist(read_counts, main = paste("Histogram: Read Counts -", name),
       xlab = "Total Reads", border = "blue", col = "green", las = 1,
       breaks = 40, axes = FALSE)
  axis(1, at = seq(0, max(read_counts), by = 1000), las = 2)
  axis(2)
  dev.off()
}
```

We decided on the following filters: 
ORAL = 2000
  RECTAL = 2000
  VAGINAL = 1000
  MILK = 1000
  PUPS = 2000 
  DAMS = 1000
  PUPS_oral= 2000
  PUPS_rectal = 2000
  DAMS_oral = 5000
  DAMS_rectal = 2000
  ALL = 2000
  ALL_day1 = 1000
  ALL_day8 = 2000
  O_R_day1 = 1000
  O_R_day8 = 2000
  ORAL_day1 = 2000
  ORAL_day8 = 5000
  RECTAL_day1 = 5000
  RECTAL_day8 = 5000
  PUP_ORAL_day1 = 2000
  PUP_ORAL_day8 = 2000
  PUP_RECTAL_day1 = 2000
  PUP_RECTAL_day8 = 2000
  DAM_ORAL_day1 = 5000
  DAM_ORAL_day8 = 5000
  DAM_RECTAL_day1 = 2000
  DAM_RECTAL_day8 = 2000
  DAM_MILK_day1 = 1000
  DAM_MILK_day8 = 1000
  DAM_VAGINAL_day1 = 1000
  DAM_VAGINAL_day8 = 1000

```{r filter}
# Filter out samples that have fewer than 1000 reads
ORAL_fil <- prune_samples(sample_sums(oral_all) > 2000, oral_all)
RECTAL_fil <- prune_samples(sample_sums(rectal_all) > 2000, rectal_all)
VAGINAL_fil <- prune_samples(sample_sums(vaginal) > 1000, vaginal)
MILK_fil <- prune_samples(sample_sums(milk) > 1000, milk)
PUPS_fil <- prune_samples(sample_sums(pups_all) > 2000, pups_all)
DAMS_fil <- prune_samples(sample_sums(dams_all) > 1000, dams_all)
PUPS_oral_fil <- prune_samples(sample_sums(pups_oral) > 2000, pups_oral)
PUPS_rectal_fil <- prune_samples(sample_sums(pups_rectal) > 2000, pups_rectal)
DAMS_oral_fil <- prune_samples(sample_sums(dams_oral) > 5000, dams_oral)
DAMS_rectal_fil <- prune_samples(sample_sums(dams_rectal) > 2000, dams_rectal)
ALL_fil <- prune_samples(sample_sums(phy_study) > 2000, phy_study)
ALL_day1_fil <- prune_samples(sample_sums(all_day1) > 1000, all_day1)
ALL_day8_fil <- prune_samples(sample_sums(all_day8) > 2000, all_day8)
O_R_day1_fil <- prune_samples(sample_sums(oral_rectal_day1) > 1000, oral_rectal_day1)
O_R_day8_fil <- prune_samples(sample_sums(oral_rectal_day8) > 1000, oral_rectal_day8)
ORAL_day1_fil <- prune_samples(sample_sums(oral_day1) > 2000, oral_day1)
ORAL_day8_fil <- prune_samples(sample_sums(oral_day8) > 5000, oral_day8)
RECTAL_day1_fil <- prune_samples(sample_sums(rectal_day1) > 5000, rectal_day1)
RECTAL_day8_fil <- prune_samples(sample_sums(rectal_day8) > 5000, rectal_day8)
PUP_ORAL_day1_fil <- prune_samples(sample_sums(pup_oral_day1) > 2000, pup_oral_day1)
PUP_ORAL_day8_fil <- prune_samples(sample_sums(pup_oral_day8) > 2000, pup_oral_day8)
PUP_RECTAL_day1_fil <- prune_samples(sample_sums(pup_rectal_day1) > 2000, pup_rectal_day1)
PUP_RECTAL_day8_fil <- prune_samples(sample_sums(pup_rectal_day8) > 2000, pup_rectal_day8)
DAM_ORAL_day1_fil <- prune_samples(sample_sums(dam_oral_day1) > 5000, dam_oral_day1)
DAM_ORAL_day8_fil <- prune_samples(sample_sums(dam_oral_day8) > 5000, dam_oral_day8)
DAM_RECTAL_day1_fil <- prune_samples(sample_sums(dam_rectal_day1) > 2000, dam_rectal_day1)
DAM_RECTAL_day8_fil <- prune_samples(sample_sums(dam_rectal_day8) > 2000, dam_rectal_day8)
DAM_MILK_day1_fil <- prune_samples(sample_sums(dam_milk_day1) > 1000, dam_milk_day1)
DAM_MILK_day8_fil <- prune_samples(sample_sums(dam_milk_day8) > 1000, dam_milk_day8)
DAM_VAGINAL_day1_fil <- prune_samples(sample_sums(dam_vaginal_day1) > 1000, dam_vaginal_day1)
DAM_VAGINAL_day8_fil <- prune_samples(sample_sums(dam_vaginal_day8) > 1000, dam_vaginal_day8)

datasets_fil <- list(
  ORAL_fil = ORAL_fil,
  RECTAL_fil = RECTAL_fil,
  VAGINAL_fil = VAGINAL_fil,
  MILK_fil = MILK_fil,
  PUPS_fil = PUPS_fil,
  DAMS_fil = DAMS_fil,
  PUPS_oral_fil = PUPS_oral_fil,
  PUPS_rectal_fil = PUPS_rectal_fil,
  DAMS_oral_fil = DAMS_oral_fil,
  DAMS_rectal_fil = DAMS_rectal_fil,
  ALL_fil = ALL_fil,
  ALL_day1_fil = ALL_day1_fil,
  ALL_day8_fil = ALL_day8_fil,
  O_R_day1_fil = O_R_day1_fil,
  O_R_day8_fil = O_R_day8_fil,
  ORAL_day1_fil = ORAL_day1_fil,
  ORAL_day8_fil = ORAL_day8_fil,
  RECTAL_day1_fil = RECTAL_day1_fil,
  RECTAL_day8_fil = RECTAL_day8_fil,
  PUP_ORAL_day1_fil = PUP_ORAL_day1_fil,
  PUP_ORAL_day8_fil = PUP_ORAL_day8_fil,
  PUP_RECTAL_day1_fil = PUP_RECTAL_day1_fil,
  PUP_RECTAL_day8_fil = PUP_RECTAL_day8_fil,
  DAM_ORAL_day1_fil = DAM_ORAL_day1_fil,
  DAM_ORAL_day8_fil = DAM_ORAL_day8_fil,
  DAM_RECTAL_day1_fil = DAM_RECTAL_day1_fil,
  DAM_RECTAL_day8_fil = DAM_RECTAL_day8_fil,
  DAM_MILK_day1_fil = DAM_MILK_day1_fil,
  DAM_MILK_day8_fil = DAM_MILK_day8_fil,
  DAM_VAGINAL_day1_fil = DAM_VAGINAL_day1_fil,
  DAM_VAGINAL_day8_fil = DAM_VAGINAL_day8_fil

)

# Create workbook
wb <- createWorkbook()

# Loop over datasets
for (name in names(datasets_fil)) {
  phy_filtered <- datasets_fil[[name]]
  
  # Create metadata df with total read counts
  metadata_filtered <- data.frame(sample_data(phy_filtered))
  metadata_filtered$total_reads <- sample_sums(phy_filtered)
  
  # Add as a sheet in Excel workbook
  addWorksheet(wb, sheetName = name)
  writeData(wb, sheet = name, x = metadata_filtered)
} 
# Save Excel workbook
saveWorkbook(wb, file = paste0(results_dir, "tables/filtered_metadata_with_total_reads.xlsx"), overwrite = TRUE)

```

## Normalization 

## Rarefication

Normalize number of reads in each sample using median sequencing depth. == RAREFICATION

We chose to rarefy for each group separately

```{r}
# Set global seed for reproducibility
set.seed(1)

# List of datasets and their corresponding names
datasets_fil <- list(
  ORAL_fil = ORAL_fil,
  RECTAL_fil = RECTAL_fil,
  VAGINAL_fil = VAGINAL_fil,
  MILK_fil = MILK_fil,
  PUPS_fil = PUPS_fil,
  DAMS_fil = DAMS_fil,
  PUPS_oral_fil = PUPS_oral_fil,
  PUPS_rectal_fil = PUPS_rectal_fil,
  DAMS_oral_fil = DAMS_oral_fil,
  DAMS_rectal_fil = DAMS_rectal_fil,
  ALL_fil = ALL_fil,
  ALL_day1_fil = ALL_day1_fil,
  ALL_day8_fil = ALL_day8_fil,
  O_R_day1_fil = O_R_day1_fil,
  O_R_day8_fil = O_R_day8_fil,
  ORAL_day1_fil = ORAL_day1_fil,
  ORAL_day8_fil = ORAL_day8_fil,
  RECTAL_day1_fil = RECTAL_day1_fil,
  RECTAL_day8_fil = RECTAL_day8_fil,
  PUP_ORAL_day1_fil = PUP_ORAL_day1_fil,
  PUP_ORAL_day8_fil = PUP_ORAL_day8_fil,
  PUP_RECTAL_day1_fil = PUP_RECTAL_day1_fil,
  PUP_RECTAL_day8_fil = PUP_RECTAL_day8_fil,
  DAM_ORAL_day1_fil = DAM_ORAL_day1_fil,
  DAM_ORAL_day8_fil = DAM_ORAL_day8_fil,
  DAM_RECTAL_day1_fil = DAM_RECTAL_day1_fil,
  DAM_RECTAL_day8_fil = DAM_RECTAL_day8_fil,
  DAM_MILK_day1_fil = DAM_MILK_day1_fil,
  DAM_MILK_day8_fil = DAM_MILK_day8_fil,
  DAM_VAGINAL_day1_fil = DAM_VAGINAL_day1_fil,
  DAM_VAGINAL_day8_fil = DAM_VAGINAL_day8_fil
  )

dataset_names <- names(datasets_fil)

# Loop through each dataset
for (i in 1:length(datasets_fil)) {
  # Set current dataset and its name
  rar_data <- datasets_fil[[i]]
  rar_data_n <- dataset_names[i]

  # Rarefy the data
  phy_study_rar <- rarefy_even_depth(rar_data, rngseed=1, sample.size=0.9*min(sample_sums(rar_data)), replace=F)
  
  # Save the rarefied phyloseq object if needed
  assign(paste0("rar_", rar_data_n), phy_study_rar)
}

datasets_fil_rar <- list(
  rar_ORAL_fil = rar_ORAL_fil,
  rar_RECTAL_fil = rar_RECTAL_fil,
  rar_VAGINAL_fil = rar_VAGINAL_fil,
  rar_MILK_fil = rar_MILK_fil,
  rar_PUPS_fil = rar_PUPS_fil,
  rar_DAMS_fil = rar_DAMS_fil,
  rar_PUPS_oral_fil = rar_PUPS_oral_fil,
  rar_PUPS_rectal_fil = rar_PUPS_rectal_fil,
  rar_DAMS_oral_fil = rar_DAMS_oral_fil,
  rar_DAMS_rectal_fil = rar_DAMS_rectal_fil,
  rar_ALL_fil = rar_ALL_fil,
  rar_ALL_day1_fil = rar_ALL_day1_fil,
  rar_ALL_day8_fil = rar_ALL_day8_fil,
  rar_O_R_day1_fil = rar_O_R_day1_fil,
  rar_O_R_day8_fil = rar_O_R_day8_fil,
  rar_ORAL_day1_fil = rar_ORAL_day1_fil,
  rar_ORAL_day8_fil = rar_ORAL_day8_fil,
  rar_RECTAL_day1_fil = rar_RECTAL_day1_fil,
  rar_RECTAL_day8_fil = rar_RECTAL_day8_fil,
  rar_PUP_ORAL_day1_fil = rar_PUP_ORAL_day1_fil,
  rar_PUP_ORAL_day8_fil = rar_PUP_ORAL_day8_fil,
  rar_PUP_RECTAL_day1_fil = rar_PUP_RECTAL_day1_fil,
  rar_PUP_RECTAL_day8_fil = rar_PUP_RECTAL_day8_fil,
  rar_DAM_ORAL_day1_fil = rar_DAM_ORAL_day1_fil,
  rar_DAM_ORAL_day8_fil = rar_DAM_ORAL_day8_fil,
  rar_DAM_RECTAL_day1_fil = rar_DAM_RECTAL_day1_fil,
  rar_DAM_RECTAL_day8_fil = rar_DAM_RECTAL_day8_fil,
  rar_DAM_MILK_day1_fil = rar_DAM_MILK_day1_fil,
  rar_DAM_MILK_day8_fil = rar_DAM_MILK_day8_fil,
  rar_DAM_VAGINAL_day1_fil = rar_DAM_VAGINAL_day1_fil,
  rar_DAM_VAGINAL_day8_fil = rar_DAM_VAGINAL_day8_fil
  )
```

## Relative Abundance
Transform to Relative Abundance and generate a bar plot 
(transform the original object, not the rarified object)

```{r, include=FALSE}
# Get dataset names
dataset_names <- names(datasets_fil)

# Loop through each dataset
for (i in seq_along(datasets_fil)) {
  data <- datasets_fil[[i]]
  data_n <- dataset_names[i]

  # Modify CategoricalDay levels
  data@sam_data$CategoricalDay <- factor(data@sam_data$CategoricalDay, 
                                         levels = c("1'", "8'"), 
                                         labels = c("Day 1", "Day 8"))
  
# Scale to relative abundance (percent)
  physeq_scaled <- transform_sample_counts(data, function(y) 100 * y / sum(y))
  
  # Create object in the environment with name like "ORAL_fil_scaled"
  assign(paste0(data_n, "_scaled"), physeq_scaled)
}
```

Generate bar plots of relative abundance

```{r, include=FALSE}
tax_ranks <- c("Class", "Family", "Genus", "Species")
topN <- 15

plot_dir <- file.path(results_dir, "plots/relative_bar_plots")
dir.create(plot_dir, recursive = TRUE, showWarnings = FALSE)

datasets_fil_scaled <- list(
  ORAL_fil_scaled = ORAL_fil_scaled,
  RECTAL_fil_scaled = RECTAL_fil_scaled,
  VAGINAL_fil_scaled = VAGINAL_fil_scaled,
  MILK_fil_scaled = MILK_fil_scaled,
  PUPS_fil_scaled = PUPS_fil_scaled,
  DAMS_fil_scaled = DAMS_fil_scaled,
  PUPS_oral_fil_scaled = PUPS_oral_fil_scaled,
  PUPS_rectal_fil_scaled = PUPS_rectal_fil_scaled,
  DAMS_oral_fil_scaled = DAMS_oral_fil_scaled,
  DAMS_rectal_fil_scaled = DAMS_rectal_fil_scaled,
  ALL_fil_scaled = ALL_fil_scaled,
  ALL_day1_fil_scaled = ALL_day1_fil_scaled,
  ALL_day8_fil_scaled = ALL_day8_fil_scaled,
  O_R_day1_fil_scaled = O_R_day1_fil_scaled,
  O_R_day8_fil_scaled = O_R_day8_fil_scaled
)

dataset_names <- names(datasets_fil_scaled)

# ---- LOOP THROUGH DATASETS ----

for (i in seq_along(datasets_fil_scaled)) {
  
  data <- datasets_fil_scaled[[i]]
  data_n <- dataset_names[i]
  message("\n============================")
  message("Processing dataset: ", data_n)
  message("============================")

# ---- LOOP THROUGH TAXONOMIC RANKS FOR THIS DATASET ----
  
  for (rank in tax_ranks) {
    message("  Rank: ", rank)
    
    # Try to agglomerate
    phy_rank <- tryCatch(tax_glom(data, taxrank = rank), error = function(e) NULL)
    
    if (is.null(phy_rank) || ntaxa(phy_rank) == 0) {
      message("    Skipping: no taxa for rank ", rank)
      next
    }
    
    # Convert to relative abundance
    phy_rank <- transform_sample_counts(phy_rank, function(x) x / sum(x))
    
    # ---- clean taxonomy ----
    tt <- as.data.frame(tax_table(phy_rank))
    if (rank %in% colnames(tt)) {
      tt[[rank]][is.na(tt[[rank]]) | tt[[rank]] == ""] <- "Unclassified"
    } else {
      tt[[rank]] <- "Unclassified"
    }
    tax_table(phy_rank) <- as.matrix(tt)
    
    # ---- compute top N ----
    taxa_sums_rank <- taxa_sums(phy_rank)
    top_taxa <- names(sort(taxa_sums_rank, decreasing = TRUE))[1:min(topN, length(taxa_sums_rank))]
    
    # collapse non-top taxa
    tax_table(phy_rank)[!taxa_names(phy_rank) %in% top_taxa, rank] <- "Other"
    
    # ---- save plot ----
    plot_path <- file.path(
      plot_dir,
      paste0(data_n, "_rarefied_relative_abundance_barplot_", rank, ".tiff")
    )
    
    tiff(filename = plot_path, width = 10, height = 4, units = "in", res = 300)
    
    p <- plot_bar(phy_rank, fill = rank) +
      geom_bar(aes(fill = !!sym(rank), color = !!sym(rank)), 
               stat = "identity", position = "stack") +
      facet_grid(~ CategoricalDay, scales = "free", space = "free") +
      guides(fill = guide_legend(ncol = 2)) +
      ggtitle(paste(data_n, "- Relative Abundance at", rank, "Level"))
    
    print(p)
    dev.off()
  }
}
```

# Statistics

## Alpha diversity

Matched dam-litter comparisons within day and site

```{r alpha Mann-Whitney}
# Create the directories
dir.create(file.path(results_dir, "tables/alpha_diversity/"), recursive = TRUE, showWarnings = FALSE)
dir.create(file.path(results_dir, "plots/alpha_diversity/EPS/"), recursive = TRUE, showWarnings = FALSE)

# Create a new workbook
wb <- createWorkbook()

# Define datasets to compare dams vs pups within the same day and site
datasets_fil_rar_alpha <- list(
  ORAL_day1 = rar_ORAL_day1_fil,
  ORAL_day8 = rar_ORAL_day8_fil,
  RECTAL_day1 = rar_RECTAL_day1_fil,
  RECTAL_day8 = rar_RECTAL_day8_fil
)

dataset_names <- names(datasets_fil_rar_alpha)

# Prepare colors for Faith's PD plots
TypeHostNew_colors_custom <- TypeHostNew_colors
groups_to_clear <- c("HealthyPupRectal1", "HealthyPupRectal8", "HealthyPupOral1", "HealthyPupOral8")
TypeHostNew_colors_custom[groups_to_clear] <- "white"

for (i in seq_along(datasets_fil_rar_alpha)) {
  rar_data <- datasets_fil_rar_alpha[[i]]
  rar_data_n <- dataset_names[i]

  # Plot alpha diversity distributions
  tiff(filename = paste0(results_dir, "plots/alpha_diversity/", rar_data_n, "_alpha_diversity.tiff"), width = 10, height = 4, units = "in", res = 300)
  alpha_plot <- plot_richness(rar_data, x = "HostTypeDay", measures = c("Observed", "Chao1", "Shannon")) + 
    geom_boxplot() +
    scale_x_discrete(labels = c("1'" = "Day 1", "8'" = "Day 8"))
  print(alpha_plot)
  dev.off()

  # Calculate alpha diversity indices
  rar_richness <- estimate_richness(rar_data, measures = c("Observed", "Chao1", "Shannon"))
  alpha_diversity_pd <- estimate_pd(rar_data)
  rar_richness_all <- cbind(sample_data(rar_data), rar_richness, alpha_diversity_pd)
  rar_richness_all$HostTypeDay <- factor(
    rar_richness_all$HostTypeDay,
    levels = c("DamMilk1", "DamMilk8", "HealthyPupOral1", "HealthyPupOral8", "DamOral1", "DamOral8", "HealthyPupRectal1", "HealthyPupRectal8", "DamRectal1", "DamRectal8", "DamVaginal1", "DamVaginal8")
  )

  # Faith's PD figure
  tiff(filename = paste0(results_dir, "plots/alpha_diversity/", rar_data_n, "_alpha_diversity_Faith PD.tiff"), width = 10, height = 4, units = "in", res = 300)
  alpha_plot2 <- rar_richness_all %>%
    ggplot(aes(x = HostTypeDay, y = PD, fill = HostTypeDay, color = HostTypeDay)) +
    geom_boxplot(outlier.shape = NA, alpha = 0.5) +
    geom_jitter(aes(shape = HostTypeDay), position = position_jitter(width = 0.2), size = 2) +
    ggtitle("") + xlab("") + ylab("Faith's Phylogenetic Diversity") +
    scale_fill_manual(values = TypeHostNew_colors_custom) +
    scale_color_manual(values = TypeHostNew_colors) +
    scale_shape_manual(values = TypeHostNew_shapes) +
    theme_classic() +
    theme(
      legend.position = "none",
      axis.title.y = element_text(size = 15),
      axis.text = element_text(size = 15),
      axis.title = element_text(size = 20)
    ) +
    scale_x_discrete(labels = c("1'" = "Day 1", "8'" = "Day 8"))
  print(alpha_plot2)
  dev.off()

  ggsave(
    filename = paste0(results_dir, "plots/alpha_diversity/EPS/", rar_data_n, "_alpha_diversity_Faith PD.eps"),
    plot = alpha_plot2,
    device = cairo_ps,
    width = 7,
    height = 4,
    dpi = 600
  )

  # Build matched dam-litter pairs for testing
  df <- as.data.frame(rar_richness_all) %>%
    mutate(
      PairID = case_when(
        grepl("Pup", HostTypeDay) ~ as.character(Mother),
        TRUE ~ as.character(Host_ID)
      ),
      HostGroup = ifelse(grepl("Pup", HostTypeDay), "Pup", "Dam")
    )

  pup_litter_summary <- df %>%
    filter(HostGroup == "Pup") %>%
    group_by(PairID, HostTypeDay) %>%
    summarise(
      n_pups = n(),
      Observed_pup = median(Observed, na.rm = TRUE),
      Chao1_pup = median(Chao1, na.rm = TRUE),
      Shannon_pup = median(Shannon, na.rm = TRUE),
      PD_pup = median(PD, na.rm = TRUE),
      .groups = "drop"
    )

  dam_summary <- df %>%
    filter(HostGroup == "Dam") %>%
    group_by(PairID, HostTypeDay) %>%
    summarise(
      n_dams = n(),
      Observed_dam = median(Observed, na.rm = TRUE),
      Chao1_dam = median(Chao1, na.rm = TRUE),
      Shannon_dam = median(Shannon, na.rm = TRUE),
      PD_dam = median(PD, na.rm = TRUE),
      .groups = "drop"
    )

  paired_alpha <- inner_join(dam_summary, pup_litter_summary, by = "PairID") %>%
    mutate(
      Dataset = rar_data_n,
      DamHostTypeDay = HostTypeDay.x,
      PupHostTypeDay = HostTypeDay.y
    ) %>%
    select(
      Dataset, PairID, DamHostTypeDay, PupHostTypeDay, n_dams, n_pups,
      Observed_dam, Observed_pup,
      Chao1_dam, Chao1_pup,
      Shannon_dam, Shannon_pup,
      PD_dam, PD_pup
    )

  metrics <- c("Observed", "Chao1", "Shannon", "PD")
  paired_results <- lapply(metrics, function(metric_name) {
    dam_vals <- paired_alpha[[paste0(metric_name, "_dam")]]
    pup_vals <- paired_alpha[[paste0(metric_name, "_pup")]]
    keep <- complete.cases(dam_vals, pup_vals)

    if (sum(keep) == 0) {
      return(tibble(
        Dataset = rar_data_n,
        Metric = metric_name,
        n_pairs = 0,
        statistic = NA_real_,
        p.value = NA_real_,
        dam_median = NA_real_,
        pup_median = NA_real_,
        median_difference = NA_real_
      ))
    }

    test_res <- wilcox.test(dam_vals[keep], pup_vals[keep], paired = TRUE, exact = FALSE)

    tibble(
      Dataset = rar_data_n,
      Metric = metric_name,
      n_pairs = sum(keep),
      statistic = unname(test_res$statistic),
      p.value = test_res$p.value,
      dam_median = median(dam_vals[keep], na.rm = TRUE),
      pup_median = median(pup_vals[keep], na.rm = TRUE),
      median_difference = median(dam_vals[keep] - pup_vals[keep], na.rm = TRUE)
    )
  }) %>%
    bind_rows() %>%
    mutate(p.adjust = p.adjust(p.value, method = "BH"))

  addWorksheet(wb, paste0(rar_data_n, "_paired_input"))
  writeData(wb, sheet = paste0(rar_data_n, "_paired_input"), x = paired_alpha)

  addWorksheet(wb, paste0(rar_data_n, "_paired_wilcox"))
  writeData(wb, sheet = paste0(rar_data_n, "_paired_wilcox"), x = paired_results)
}

saveWorkbook(wb, file = paste0(results_dir, "tables/alpha_diversity/alpha_diversity_matched_dam_pup.xlsx"), overwrite = TRUE)
```

Paired (comparing day 1 and day 8)

```{r}
# Create the directories
dir.create(file.path(results_dir, "tables/alpha_diversity/"), recursive = TRUE, showWarnings = FALSE)
dir.create(file.path(results_dir, "plots/alpha_diversity/EPS/"), recursive = TRUE, showWarnings = FALSE)

# Create a new workbook
wb <- createWorkbook()

# Define a dataset
datasets_fil_rar_alpha <- list(
  rar_VAGINAL_fil = rar_VAGINAL_fil,
  rar_MILK_fil = rar_MILK_fil,
  rar_DAMS_oral_fil = rar_DAMS_oral_fil,
  rar_DAMS_rectal_fil = rar_DAMS_rectal_fil,
  rar_PUPS_oral_fil = rar_PUPS_oral_fil,
  rar_PUPS_rectal_fil = rar_PUPS_rectal_fil
  )

# Get dataset names
dataset_names <- names(datasets_fil_rar_alpha)

# Loop through each dataset
for (i in 1:length(datasets_fil_rar_alpha)) {
  # Set current dataset and its name
  rar_data <- datasets_fil_rar_alpha[[i]]
  rar_data_n <- dataset_names[i]

# Prepare new colors for boxplots
# Copy original colors and override selected groups with white fill
TypeHostNew_colors_custom <- TypeHostNew_colors
groups_to_clear <- c("HealthyPupRectal1", "HealthyPupRectal8", "HealthyPupOral1", "HealthyPupOral8")
TypeHostNew_colors_custom[groups_to_clear] <- "white"  # or "transparent" for invisible fill

#boxplot
tiff(filename = paste0(results_dir,"plots/alpha_diversity/",rar_data_n,"_alpha_diversity.tiff"), width=10, height=4, units='in', res=300)
alpha_plot <- plot_richness(rar_data, x="HostTypeDay", measures=c("Observed", "Chao1", "Shannon")) + geom_boxplot() +
    scale_x_discrete(labels=c("1'" = "Day 1", "8'" = "Day 8"))
print(alpha_plot)  # Explicitly print the plot
dev.off()

# Calculate alpha diversity indices
rar_richness <- estimate_richness(rar_data, measures = c("Observed", "Chao1", "Shannon"))
# also: Faith's PD
alpha_diversity_pd <- estimate_pd(rar_data) #Faith
# Join the different indices 
rar_richness_all <- cbind(sample_data(rar_data), rar_richness, alpha_diversity_pd)
# Order by Type for the figures level
rar_richness_all$HostTypeDay <- factor(rar_richness_all$HostTypeDay,
  levels = c("DamMilk1", "DamMilk8", "HealthyPupOral1", "HealthyPupOral8", "DamOral1", "DamOral8", "HealthyPupRectal1", "HealthyPupRectal8", "DamRectal1", "DamRectal8", "DamVaginal1", "DamVaginal8"))


tiff(filename = paste0(results_dir,"plots/alpha_diversity/",rar_data_n,"_alpha_diversity_Faith PD.tiff"), width=10, height=4, units='in', res=300)
alpha_plot2 <- rar_richness_all %>%
ggplot( aes(x=HostTypeDay, y=(PD), fill= HostTypeDay,  color =  HostTypeDay)) +
    geom_boxplot(outlier.shape = NA, alpha = 0.5) +
    geom_jitter(aes(shape = HostTypeDay), position = position_jitter(width = 0.2), size = 2) +
    ggtitle("") + xlab("") + ylab("Faith's Phylogenetic Diversity") +
    scale_fill_manual(values = TypeHostNew_colors_custom) +  # modified for white fill
    scale_color_manual(values = TypeHostNew_colors ) +
    scale_shape_manual(values = TypeHostNew_shapes) +
    theme_classic() +
    theme(legend.position = "none",
          axis.title.y = element_text(size = 15),  # specific to y-axis title
    ) +
    theme(axis.text = element_text(size = 15), axis.title = element_text(size = 20))  +
    scale_x_discrete(labels=c("1'" = "Day 1", "8'" = "Day 8"))
print(alpha_plot2)  # Explicitly print the plot
dev.off()

# Save the plot as a EPS file
ggsave(
  filename = paste0(results_dir, "plots/alpha_diversity/EPS/",rar_data_n,"_alpha_diversity_Faith PD.eps"),
  plot = alpha_plot2,
  device = cairo_ps,
  width = 7,
  height = 4,
  dpi = 600
)
print(alpha_plot2)  # Save the plot to the file
dev.off()


# Kruskal-Wallis test
  kw_results <- rar_richness_all %>%
    select(HostTypeDay, Observed, Chao1, Shannon, PD) %>%
    summarise(
      Observed_KW = kruskal.test(Observed ~ HostTypeDay)$p.value,
      Chao1_KW = kruskal.test(Chao1 ~ HostTypeDay)$p.value,
      Shannon_KW = kruskal.test(Shannon ~ HostTypeDay)$p.value,
      PD_KW = kruskal.test(PD ~ HostTypeDay)$p.value
    )

  # Dunn's test
  dunn_results <- rar_richness_all %>%
    select(HostTypeDay, Observed, Chao1, Shannon, PD) %>%
    pivot_longer(cols = c(Observed, Chao1, Shannon, PD), names_to = "Metric", values_to = "Value") %>%
    group_by(Metric) %>%
    dunn_test(Value ~ HostTypeDay, p.adjust.method = "bonferroni", detailed = TRUE) %>%
    mutate(Dataset = rar_data_n) # Add dataset name

# Paired Wilcoxon tests (Day 1 vs Day 8 within each host/site)

paired_alpha <- rar_richness_all %>%
  select(Host_new, Host_ID, Day, Observed, Chao1, Shannon, PD)

# wide format
paired_alpha_wide <- paired_alpha %>%
  pivot_wider(id_cols = c(Host_new, Host_ID),
              names_from = Day,
              values_from = c(Observed, Chao1, Shannon, PD))

wilcox_paired_results <- tibble(
  Metric = c("Observed","Chao1","Shannon","PD"),
  p.value = NA_real_
)

for (m in wilcox_paired_results$Metric) {

  day1_col <- paste0(m, "_1")
  day8_col <- paste0(m, "_8")

  # skip if dataset has only one day
  if(!(day1_col %in% names(paired_alpha_wide) &
       day8_col %in% names(paired_alpha_wide))) {
    next
  }

  v_day1 <- paired_alpha_wide[[day1_col]]
  v_day8 <- paired_alpha_wide[[day8_col]]

  keep <- !(is.na(v_day1) | is.na(v_day8))

  if(sum(keep) > 1){
    test_res <- wilcox.test(v_day1[keep], v_day8[keep], paired = TRUE)

    wilcox_paired_results$p.value[
      wilcox_paired_results$Metric == m
    ] <- test_res$p.value
  }
}

wilcox_paired_results <- wilcox_paired_results %>%
  mutate(
    p.adj = p.adjust(p.value, method = "bonferroni"),
    Dataset = rar_data_n
  )
  
# Add each to the workbook as a new sheet
  addWorksheet(wb, paste0(rar_data_n, "_KW"))
  writeData(wb, sheet = paste0(rar_data_n, "_KW"),
            x = kw_results)

  addWorksheet(wb, paste0(rar_data_n, "_Dunn"))
  writeData(wb, sheet = paste0(rar_data_n, "_Dunn"), x = dunn_results)

  addWorksheet(wb, paste0(rar_data_n, "_PairedWX"))
  writeData(wb,sheet = paste0(rar_data_n, "_PairedWX"),
            x = wilcox_paired_results)
}

# Save final workbook with all sheets
saveWorkbook(wb, file = paste0(results_dir, "tables/alpha_diversity/alpha_diversity_paired.xlsx"), overwrite = TRUE)
```

## Beta diversity

PCoA plot using the weighted UniFrac as distance

```{r}
# Get dataset names
dataset_names <- names(datasets_fil_rar)

# Loop through each dataset
for (i in 1:length(datasets_fil_rar)) {
  # Set current dataset and its name
  rar_data <- datasets_fil_rar[[i]]
  rar_data_n <- dataset_names[i]

# Modify the levels of CategoricalDay directly
rar_data@sam_data$CategoricalDay <- factor(rar_data@sam_data$CategoricalDay, 
                                           levels = c("1'", "8'"), 
                                           labels = c("Day 1", "Day 8"))


wunifrac_dist <- phyloseq::distance(rar_data, method="unifrac", weighted=T)
ordination <- ordinate(rar_data, method="PCoA", distance=wunifrac_dist)

tiff(filename = paste0(results_dir,"plots/beta_diversity/weighted_unifraq/",rar_data_n,"_beta_diversity_weighted_unifraq.tiff"), width=6, height=4, units='in', res=300)
beta_w_unifraq <- plot_ordination(rar_data, ordination, color="HostTypeDay") + 
  theme(aspect.ratio=1) +
  stat_ellipse(aes(group = HostTypeDay), linewidth = 1) +  # Add 95% ellipses
  geom_point(aes(shape = HostTypeDay), size = 4,  fill = "white") +  
  scale_color_manual(values = TypeHostNew_colors) +
  scale_shape_manual(values = TypeHostNew_shapes) +
  ggtitle(paste0("Beta Diversity - ",rar_data_n)) +
  theme_classic() +
  theme(axis.text=element_text(size=15),
        axis.title=element_text(size=15),
        legend.title = element_blank())  # Remove the legend title

print(beta_w_unifraq)  # Explicitly print the plot
dev.off()

tiff(filename = paste0(results_dir,"plots/beta_diversity/weighted_unifraq/",rar_data_n,"_beta_diversity_weighted_unifraq_no_ellipse.tiff"), width=6, height=4, units='in', res=300)
beta_w_unifraq2 <- plot_ordination(rar_data, ordination, color="HostTypeDay") + 
  theme(aspect.ratio=1) +
  geom_point(aes(shape = HostTypeDay), size = 4,  fill = "white") +  
  scale_color_manual(values = TypeHostNew_colors) +
  scale_shape_manual(values = TypeHostNew_shapes) +
  ggtitle(paste0("Beta Diversity - ",rar_data_n)) +
  theme_classic() +
  theme(axis.text=element_text(size=15),
        axis.title=element_text(size=15),
        legend.title = element_blank())  # Remove the legend title

print(beta_w_unifraq2)  # Explicitly print the plot
dev.off()

tiff(filename = paste0(results_dir,"plots/beta_diversity/weighted_unifraq/",rar_data_n,"_beta_diversity_weighted_unifraq_no_ellipse_no_legend.tiff"), width=6, height=4, units='in', res=300)
beta_w_unifraq3 <- plot_ordination(rar_data, ordination, color="HostTypeDay") + 
  theme(aspect.ratio=1) +
  geom_point(aes(shape = HostTypeDay), 
             size = 4, 
             fill = "white", 
             stroke = 2) +   # increase thickness of marker borders
  scale_color_manual(values = TypeHostNew_colors) +
  scale_shape_manual(values = TypeHostNew_shapes) +
  ggtitle(paste0("Beta Diversity - ", rar_data_n)) +
  theme_classic() +
  theme(axis.text=element_text(size=15),
        axis.title=element_text(size=15),
        legend.title = element_blank(),
        legend.position = "none" )  
print(beta_w_unifraq3)  # Explicitly print the plot
dev.off()


tiff(filename = paste0(results_dir,"plots/beta_diversity/weighted_unifraq/",rar_data_n,"_beta_diversity_weighted_unifraq_no_ellipse_connected_points.tiff"), width=6, height=4, units='in', res=300)
beta_w_unifraq4 <- plot_ordination(rar_data, ordination, color = "HostTypeDay") + 
  theme(aspect.ratio = 1) +
  geom_point(aes(shape = HostTypeDay), size = 4, fill = "white") +  
  geom_line(aes(group = Host_ID, color = HostTypeDay), linewidth = 0.5) +  # Add lines connecting points by Host_ID
  scale_color_manual(values = TypeHostNew_colors) +
  scale_shape_manual(values = TypeHostNew_shapes) +
  ggtitle(paste0("Beta Diversity - ", rar_data_n)) +
  theme_classic() +
  theme(
    axis.text = element_text(size = 15),
    axis.title = element_text(size = 15),
    legend.title = element_blank()
  )
print(beta_w_unifraq4)  # Explicitly print the plot
dev.off()

# Save the plot as a EPS file
ggsave(
  filename = paste0(results_dir, "plots/beta_diversity/weighted_unifraq/EPS/",rar_data_n,"_beta_diversity_weighted_unifraq_no_ellipse.eps"),
  plot = beta_w_unifraq3,
  device = cairo_ps,
  width = 7,
  height = 4,
  dpi = 600
)
print(beta_w_unifraq3)  # Save the plot to the file
dev.off()

}
```

For Pups&Dams only

```{r}
# Define data set
datasets_OR <- list(rar_O_R_day1_fil = rar_O_R_day1_fil,
                      rar_O_R_day8_fil = rar_O_R_day8_fil)

# Get dataset names
dataset_names <- names(datasets_OR)

# pre-loop code to extract global limits
# Initialize vectors to store all Axis.1 and Axis.2 values
all_axis1 <- c()
all_axis2 <- c()

# First pass: extract ordination values across all datasets
for (i in 1:length(datasets_OR)) {
  rar_data <- datasets_OR[[i]]
  wunifrac_dist <- phyloseq::distance(rar_data, method="unifrac", weighted=T)
  ordination <- ordinate(rar_data, method="PCoA", distance=wunifrac_dist)
  ord_df <- as.data.frame(ordination$vectors)
  
  all_axis1 <- c(all_axis1, ord_df[, 1])
  all_axis2 <- c(all_axis2, ord_df[, 2])
}

# Define global limits
xlims <- range(all_axis1)
ylims <- range(all_axis2)

# Loop through each dataset
for (i in 1:length(datasets_OR)) {
  # Set current dataset and its name
  rar_data <- datasets_OR[[i]]
  rar_data_n <- dataset_names[i]

# Modify the levels of CategoricalDay directly
rar_data@sam_data$CategoricalDay <- factor(rar_data@sam_data$CategoricalDay, 
                                           levels = c("1'", "8'"), 
                                           labels = c("Day 1", "Day 8"))

wunifrac_dist <- phyloseq::distance(rar_data, method="unifrac", weighted=T)
ordination <- ordinate(rar_data, method="PCoA", distance=wunifrac_dist)

tiff(filename = paste0(results_dir,"plots/beta_diversity/weighted_unifraq/",rar_data_n,"_beta_diversity_weighted_unifraq.tiff"), width=6, height=4, units='in', res=300)
beta_w_unifraq <- plot_ordination(rar_data, ordination, color="HostTypeDay") + 
  xlim(xlims) + ylim(ylims) +
  theme(aspect.ratio=1) +
  stat_ellipse(aes(group = HostTypeDay), linewidth = 1) +  # Add 95% ellipses
  geom_point(aes(shape = HostTypeDay), size = 4,  fill = "white") +  
  scale_color_manual(values = TypeHostNew_colors) +
  scale_shape_manual(values = TypeHostNew_shapes) +
  ggtitle(paste0("Beta Diversity - ",rar_data_n)) +
  theme_classic() +
  theme(axis.text=element_text(size=15),
        axis.title=element_text(size=15),
        legend.title = element_blank())  # Remove the legend title

print(beta_w_unifraq)  # Explicitly print the plot
dev.off()

tiff(filename = paste0(results_dir,"plots/beta_diversity/weighted_unifraq/",rar_data_n,"_beta_diversity_weighted_unifraq_no_ellipse.tiff"), width=6, height=4, units='in', res=300)
beta_w_unifraq2 <- plot_ordination(rar_data, ordination, color="HostTypeDay") + 
  xlim(xlims) + ylim(ylims) +
  theme(aspect.ratio=1) +
  geom_point(aes(shape = HostTypeDay), size = 4,  fill = "white") +  
  scale_color_manual(values = TypeHostNew_colors) +
  scale_shape_manual(values = TypeHostNew_shapes) +
  ggtitle(paste0("Beta Diversity - ",rar_data_n)) +
  theme_classic() +
  theme(axis.text=element_text(size=15),
        axis.title=element_text(size=15),
        legend.title = element_blank())  # Remove the legend title

print(beta_w_unifraq2)  # Explicitly print the plot
dev.off()

tiff(filename = paste0(results_dir,"plots/beta_diversity/weighted_unifraq/",rar_data_n,"_beta_diversity_weighted_unifraq_no_ellipse_no_legend.tiff"), width=6, height=4, units='in', res=300)
beta_w_unifraq3 <- plot_ordination(rar_data, ordination, color="HostTypeDay") + 
  xlim(xlims) + ylim(ylims) +
  theme(aspect.ratio=1) +
  geom_point(aes(shape = HostTypeDay), size = 4,  fill = "white") +  
  scale_color_manual(values = TypeHostNew_colors) +
  scale_shape_manual(values = TypeHostNew_shapes) +
  ggtitle(paste0("Beta Diversity - ",rar_data_n)) +
  theme_classic() +
  theme(axis.text=element_text(size=15),
        axis.title=element_text(size=15),
        legend.title = element_blank(),
        legend.position = "none" )  
print(beta_w_unifraq3)  # Explicitly print the plot
dev.off()


tiff(filename = paste0(results_dir,"plots/beta_diversity/weighted_unifraq/",rar_data_n,"_beta_diversity_weighted_unifraq_no_ellipse_connected_points.tiff"), width=6, height=4, units='in', res=300)
beta_w_unifraq4 <- plot_ordination(rar_data, ordination, color = "HostTypeDay") + 
  xlim(xlims) + ylim(ylims) +
  theme(aspect.ratio = 1) +
  geom_point(aes(shape = HostTypeDay), size = 4, fill = "white") +  
  geom_line(aes(group = Host_ID, color = HostTypeDay), linewidth = 0.5) +  # Add lines connecting points by Host_ID
  scale_color_manual(values = TypeHostNew_colors) +
  scale_shape_manual(values = TypeHostNew_shapes) +
  ggtitle(paste0("Beta Diversity - ", rar_data_n)) +
  theme_classic() +
  theme(
    axis.text = element_text(size = 15),
    axis.title = element_text(size = 15),
    legend.title = element_blank()
  )
print(beta_w_unifraq4)  # Explicitly print the plot
dev.off()


# Save the plot as a EPS file
ggsave(
  filename = paste0(results_dir, "plots/beta_diversity/weighted_unifraq/EPS/",rar_data_n,"_beta_diversity_weighted_unifraq_no_ellipse.eps"),
  plot = beta_w_unifraq3,
  device = cairo_ps,
  width = 7,
  height = 4,
  dpi = 600
)
print(beta_w_unifraq3)  # Save the plot to the file
dev.off()

}
```

## Dispersion

```{r}
# Get dataset names
dataset_names <- names(datasets_fil_rar)

# Loop through each dataset
for (i in 1:length(datasets_fil_rar)) {
  # Set current dataset and its name
  rar_data <- datasets_fil_rar[[i]]
  rar_data_n <- dataset_names[i]
  
  # Compute distance matrix
  dist_matrix <- phyloseq::distance(rar_data, method = "unifrac", weighted=T)
  
  # Extract metadata
  metadata_ad <- data.frame(sample_data(rar_data))
  
# Run betadisper (homogeneity of dispersion)
  dispersion_test <- betadisper(dist_matrix, metadata_ad$HostTypeDay)
  dispersion_perm <- permutest(dispersion_test, permutations = 9999)
    
  print(dispersion_perm)
    
  # Save permutation test results
  dispersion_perm_df <- as.data.frame(dispersion_perm$tab)
  write.csv(dispersion_perm_df, 
              file = paste0(results_dir, "tables/dispersion/", rar_data_n, "_dispersion_test.csv"), 
              row.names = TRUE)
    
   # Pairwise Wilcoxon test for dispersion differences
   dispersion_df <- data.frame(
      Group = metadata_ad$HostTypeDay,
      distances = dispersion_test$distances
    )
    
    pairwise_results <- pairwise_wilcox_test(dispersion_df, distances ~ Group, p.adjust.method = "BH")
    write.csv(pairwise_results, 
              file = paste0(results_dir, "tables/dispersion/", rar_data_n, "_dispersion_wilcox.csv"), 
              row.names = TRUE)
    
    # Generate boxplot for dispersion
    tiff(filename = paste0(results_dir, "plots/dispersion/", rar_data_n, "_dispersion.tiff"), 
         width = 6, height = 4, units = 'in', res = 300)
    
    dispersion_plot <- ggboxplot(dispersion_df, 
                                 x = "Group", 
                                 y = "distances", 
                                 fill = "Group") +
      geom_jitter(width = 0.2, size = 2, alpha = 0.7) +
      stat_compare_means(method = "wilcox.test", label = "p.signif") +
      theme_classic() +
      scale_fill_manual(values = TypeHostNew_colors ) +
      scale_color_manual(values = TypeHostNew_colors ) +
      scale_shape_manual(values = TypeHostNew_shapes) +
      labs(x = NULL, y = "Distance from Centroid") +
      theme(axis.text = element_text(size = 14),
            axis.title = element_text(size = 15),
            legend.position = "none")
    
    print(dispersion_plot)
    dev.off()
    
    print(paste("Dispersion analysis completed for:", rar_data_n))
  }

```

## Differential abundance

## (1) PERMANOVA (adonis2)

With rarefied Pups data sets

```{r}
# Define data set
datasets_pups <- list(rar_PUPS_oral_fil = rar_PUPS_oral_fil,
                      rar_PUPS_rectal_fil = rar_PUPS_rectal_fil)

# Get dataset names
dataset_names <- names(datasets_pups)

# Create a new workbook
wb <- createWorkbook()

# Loop through each dataset
for (i in 1:length(datasets_pups)) {
  # Set current dataset and its name
  rar_data <- datasets_pups[[i]]
  rar_data_n <- dataset_names[i]
  
# Compute distances and metadata
wunifrac_dist <- phyloseq::distance(rar_data, method="unifrac", weighted=T)
metadata1 <- data.frame(sample_data(rar_data))
metadata1$Temperature[metadata1$Temperature == "low"] <- NA  # Replace 'low' with NA
metadata1$Temperature <- as.numeric(metadata1$Temperature)    # Convert the column to numeric

# Run adonis2 
test.adonis <- adonis2(wunifrac_dist ~  HostTypeDay, data = metadata1, perm = 9999, by = "margin", strata = metadata1$Mother)
test.adonis
test.adonis_df <- as.data.frame(test.adonis)
test.adonis_df$variable <- rownames(test.adonis_df)


# Add sheets to the workbook
  addWorksheet(wb, sheetName = paste0(rar_data_n, ""))
  writeData(wb, sheet = paste0(rar_data_n), test.adonis_df)

}

# Save the workbook
saveWorkbook(wb, file = paste0(results_dir, "tables/adonis2/Pups_adonis2_results.xlsx"), overwrite = TRUE)
```

Dams

```{r}
# Define data set
datasets_dams <- list(rar_DAMS_oral_fil = rar_DAMS_oral_fil,
                      rar_DAMS_rectal_fil = rar_DAMS_rectal_fil,
                      rar_VAGINAL_fil = rar_VAGINAL_fil,
                      rar_MILK_fil = rar_MILK_fil)

# Get dataset names
dataset_names <- names(datasets_dams)

# Create a new workbook
wb <- createWorkbook()

# Loop through each dataset
for (i in 1:length(datasets_dams)) {
  # Set current dataset and its name
  rar_data <- datasets_dams[[i]]
  rar_data_n <- dataset_names[i]

# Compute distances and metadata
wunifrac_dist <- phyloseq::distance(rar_data, method="unifrac", weighted=T)
metadata1 <- data.frame(sample_data(rar_data))
# Remove rows with NA in the explanatory variables (e.g., 'Temperature')
cleaned_data <- metadata1 %>% filter(across(c(Litter_size), ~ !is.na(.)))


# Run adonis2 - with HostTypeDay
test.adonis <- adonis2(wunifrac_dist ~  HostTypeDay, data = metadata1, perm = 9999, by = "margin")
test.adonis
test.adonis_df <- as.data.frame(test.adonis)
test.adonis_df$variable <- rownames(test.adonis_df)


# Add sheets to the workbook
  addWorksheet(wb, sheetName = paste0(rar_data_n))
  writeData(wb, sheet = paste0(rar_data_n), test.adonis_df)

}

# Save the workbook
saveWorkbook(wb, file = paste0(results_dir, "tables/adonis2/Dams_adonis2_results.xlsx"), overwrite = TRUE)


```
Pups & Dams

```{r}
# Define data set
datasets_dams <- list(rar_O_R_day1_fil = rar_O_R_day1_fil,
                      rar_O_R_day8_fil = rar_O_R_day8_fil,
                      rar_ALL_day1_fil = rar_ALL_day1_fil,
                      rar_ALL_day8_fil = rar_ALL_day8_fil)

# Get dataset names
dataset_names <- names(datasets_dams)

# Create a new workbook
wb <- createWorkbook()

# Loop through each dataset
for (i in 1:length(datasets_dams)) {
  # Set current dataset and its name
  rar_data <- datasets_dams[[i]]
  rar_data_n <- dataset_names[i]

# Compute distances and metadata
wunifrac_dist <- phyloseq::distance(rar_data, method="unifrac", weighted=T)
metadata1 <- data.frame(sample_data(rar_data))
# Remove rows with NA in the explanatory variables (e.g., 'Temperature')
cleaned_data <- metadata1 %>% filter(across(c(Litter_size), ~ !is.na(.)))


# Run adonis2 - with HostTypeDay
test.adonis <- adonis2(wunifrac_dist ~  HostTypeDay, data = metadata1, perm = 9999, by = "margin")
test.adonis
test.adonis_df <- as.data.frame(test.adonis)
test.adonis_df$variable <- rownames(test.adonis_df)


# Add sheets to the workbook
  addWorksheet(wb, sheetName = paste0(rar_data_n))
  writeData(wb, sheet = paste0(rar_data_n), test.adonis_df)

}

# Save the workbook
saveWorkbook(wb, file = paste0(results_dir, "tables/adonis2/Pups&Dams_adonis2_results.xlsx"), overwrite = TRUE)


```

## (2) MaAsLin

With filtered (non-rarefied) Pups data sets

```{r, include=FALSE}
# Define data set
datasets_pups_fil <- list(PUPS_oral_fil = PUPS_oral_fil,
                      PUPS_rectal_fil = PUPS_rectal_fil)

# Get dataset names
dataset_names <- names(datasets_pups_fil)

# Loop through each dataset
for (i in 1:length(datasets_pups_fil)) {
  # Set current dataset and its name
  data <- datasets_pups_fil[[i]]
  data_n <- dataset_names[i]

# Modify the levels of CategoricalDay directly
data@sam_data$CategoricalDay <- factor(data@sam_data$CategoricalDay, 
                                           levels = c("1'", "8'"), 
                                           labels = c("Day 1", "Day 8"))


# Prepare metadata
df_input_metadata <- as(sample_data(data), "data.frame")

# Aggregate all levels
all_data <- aggregate_taxa(data, "short_title")
OTU1 <- as(otu_table(all_data), "matrix")
if (taxa_are_rows(all_data)) { OTU1 <- t(OTU1) }
df_input_data_a <- as.data.frame(OTU1)


# Run MaAsLin2 for all levels - short title
Maaslin2(
    input_data = df_input_data_a, 
    input_metadata = df_input_metadata, 
    min_prevalence = 0.1,
    min_abundance = 0.001,
    max_significance = 0.05,
    output = paste0(results_dir, "maaslin2/Maaslin2_results_",data_n,""),
    fixed_effects = c("CategoricalDay"),
    random_effects = c("Mother"),
    reference = c("CategoricalDay,Day 1")
)
}
```

With filtered (non-rarefied) Dams data sets
only CategoricalDay as fixed factor because:"Breed","Pregnancy","Food" because most breeds/pregnancies/food were represented by a single dam/kennel/diet -we could not include any of these as fixed or random factors. 

```{r, include=FALSE}
# Define data set
datasets_dams_fil <- list(DAMS_oral_fil = DAMS_oral_fil,
                      DAMS_rectal_fil = DAMS_rectal_fil,
                      VAGINAL_fil = VAGINAL_fil,
                      MILK_fil = MILK_fil)

# Get dataset names
dataset_names <- names(datasets_dams_fil)

# Loop through each dataset
for (i in 1:length(datasets_dams_fil)) {
  # Set current dataset and its name
  data <- datasets_dams_fil[[i]]
  data_n <- dataset_names[i]

# Modify the levels of CategoricalDay directly
data@sam_data$CategoricalDay <- factor(data@sam_data$CategoricalDay, 
                                           levels = c("1'", "8'"), 
                                           labels = c("Day 1", "Day 8"))


# Prepare metadata
df_input_metadata <- as(sample_data(data), "data.frame")

# Aggregate all levels
all_data <- aggregate_taxa(data, "short_title")
OTU1 <- as(otu_table(all_data), "matrix")
if (taxa_are_rows(all_data)) { OTU1 <- t(OTU1) }
df_input_data_a <- as.data.frame(OTU1)


# Run MaAsLin2 for all levels - short title
Maaslin2(
    input_data = df_input_data_a, 
    input_metadata = df_input_metadata, 
    min_prevalence = 0.1,
    min_abundance = 0.001,
    max_significance = 0.05,
    output = paste0(results_dir, "maaslin2/Maaslin2_results_", data_n,""),
    fixed_effects = c("CategoricalDay"),
    reference = c("CategoricalDay,Day 1")
)
}
```

With filtered (non-rarefied) combined pups/dams data sets

```{r, include=FALSE}
# Define data set
datasets_dams_fil <- list(O_R_day1_fil = O_R_day1_fil,
                      O_R_day8_fil = O_R_day8_fil)

# Get dataset names
dataset_names <- names(datasets_dams_fil)

# Loop through each dataset
for (i in 1:length(datasets_dams_fil)) {
  # Set current dataset and its name
  data <- datasets_dams_fil[[i]]
  data_n <- dataset_names[i]

# Modify the levels of CategoricalDay directly
data@sam_data$CategoricalDay <- factor(data@sam_data$CategoricalDay, 
                                           levels = c("1'", "8'"), 
                                           labels = c("Day 1", "Day 8"))


# Prepare metadata
df_input_metadata <- as(sample_data(data), "data.frame")

# Aggregate all levels
all_data <- aggregate_taxa(data, "short_title")
OTU1 <- as(otu_table(all_data), "matrix")
if (taxa_are_rows(all_data)) { OTU1 <- t(OTU1) }
df_input_data_a <- as.data.frame(OTU1)


# Run MaAsLin2 for all levels - short title
Maaslin2(
    input_data = df_input_data_a, 
    input_metadata = df_input_metadata, 
    min_prevalence = 0.1,
    min_abundance = 0.001,
    max_significance = 0.05,
    output = paste0(results_dir, "maaslin2/Maaslin2_results_", data_n,""),
    fixed_effects = c("Host"),
    reference = c("Host,Dam")
)
}
```

With filtered (non-rarefied) combined pups/dams data sets, all/separate days - RECTAL

```{r, include=FALSE}
# Define data set
datasets_sub_fil <- list(RECTAL_fil = RECTAL_fil,
                         RECTAL_day1_fil = RECTAL_day1_fil)

# Get dataset names
dataset_names <- names(datasets_sub_fil)

# Loop through each dataset
for (i in 1:length(datasets_sub_fil)) {
  # Set current dataset and its name
  data <- datasets_sub_fil[[i]]
  data_n <- dataset_names[i]

# Modify the levels of CategoricalDay directly
data@sam_data$CategoricalDay <- factor(data@sam_data$CategoricalDay, 
                                           levels = c("1'", "8'"), 
                                           labels = c("Day 1", "Day 8"))


# Prepare metadata
df_input_metadata <- as(sample_data(data), "data.frame")

# Aggregate all levels
all_data <- aggregate_taxa(data, "short_title")
OTU1 <- as(otu_table(all_data), "matrix")
if (taxa_are_rows(all_data)) { OTU1 <- t(OTU1) }
df_input_data_a <- as.data.frame(OTU1)


# Run MaAsLin2 for all levels - short title
Maaslin2(
    input_data = df_input_data_a, 
    input_metadata = df_input_metadata, 
    min_prevalence = 0.1,
    min_abundance = 0.001,
    max_significance = 0.05,
    output = paste0(results_dir, "maaslin2/Maaslin2_results_", data_n,""),
    fixed_effects = c("HostTypeDay"),
    reference = c("HostTypeDay,HealthyPupRectal1")
)
}

# Day 8
# Define data set
datasets_sub_fil <- list(RECTAL_day8_fil = RECTAL_day8_fil)

# Get dataset names
dataset_names <- names(datasets_sub_fil)

# Loop through each dataset
for (i in 1:length(datasets_sub_fil)) {
  # Set current dataset and its name
  data <- datasets_sub_fil[[i]]
  data_n <- dataset_names[i]

# Modify the levels of CategoricalDay directly
data@sam_data$CategoricalDay <- factor(data@sam_data$CategoricalDay, 
                                           levels = c("1'", "8'"), 
                                           labels = c("Day 1", "Day 8"))


# Prepare metadata
df_input_metadata <- as(sample_data(data), "data.frame")

# Aggregate all levels
all_data <- aggregate_taxa(data, "short_title")
OTU1 <- as(otu_table(all_data), "matrix")
if (taxa_are_rows(all_data)) { OTU1 <- t(OTU1) }
df_input_data_a <- as.data.frame(OTU1)


# Run MaAsLin2 for all levels - short title
Maaslin2(
    input_data = df_input_data_a, 
    input_metadata = df_input_metadata, 
    min_prevalence = 0.1,
    min_abundance = 0.001,
    max_significance = 0.05,
    output = paste0(results_dir, "maaslin2/Maaslin2_results_", data_n,""),
    fixed_effects = c("HostTypeDay"),
    reference = c("HostTypeDay,HealthyPupRectal8")
)
}
```

With filtered (non-rarefied) combined pups/dams data sets, all/separate days - ORAL

```{r, include=FALSE}
# Define data set
datasets_sub_fil <- list(ORAL_fil = ORAL_fil,
                         ORAL_day1_fil = ORAL_day1_fil)

# Get dataset names
dataset_names <- names(datasets_sub_fil)

# Loop through each dataset
for (i in 1:length(datasets_sub_fil)) {
  # Set current dataset and its name
  data <- datasets_sub_fil[[i]]
  data_n <- dataset_names[i]

# Modify the levels of CategoricalDay directly
data@sam_data$CategoricalDay <- factor(data@sam_data$CategoricalDay, 
                                           levels = c("1'", "8'"), 
                                           labels = c("Day 1", "Day 8"))


# Prepare metadata
df_input_metadata <- as(sample_data(data), "data.frame")

# Aggregate all levels
all_data <- aggregate_taxa(data, "short_title")
OTU1 <- as(otu_table(all_data), "matrix")
if (taxa_are_rows(all_data)) { OTU1 <- t(OTU1) }
df_input_data_a <- as.data.frame(OTU1)


# Run MaAsLin2 for all levels - short title
Maaslin2(
    input_data = df_input_data_a, 
    input_metadata = df_input_metadata, 
    min_prevalence = 0.1,
    min_abundance = 0.001,
    max_significance = 0.05,
    output = paste0(results_dir, "maaslin2/Maaslin2_results_", data_n,""),
    fixed_effects = c("HostTypeDay"),
    reference = c("HostTypeDay,HealthyPupOral1")
)
}

# Day 8
# Define data set
datasets_sub_fil <- list(ORAL_day8_fil = ORAL_day8_fil)

# Get dataset names
dataset_names <- names(datasets_sub_fil)

# Loop through each dataset
for (i in 1:length(datasets_sub_fil)) {
  # Set current dataset and its name
  data <- datasets_sub_fil[[i]]
  data_n <- dataset_names[i]

# Modify the levels of CategoricalDay directly
data@sam_data$CategoricalDay <- factor(data@sam_data$CategoricalDay, 
                                           levels = c("1'", "8'"), 
                                           labels = c("Day 1", "Day 8"))


# Prepare metadata
df_input_metadata <- as(sample_data(data), "data.frame")

# Aggregate all levels
all_data <- aggregate_taxa(data, "short_title")
OTU1 <- as(otu_table(all_data), "matrix")
if (taxa_are_rows(all_data)) { OTU1 <- t(OTU1) }
df_input_data_a <- as.data.frame(OTU1)


# Run MaAsLin2 for all levels - short title
Maaslin2(
    input_data = df_input_data_a, 
    input_metadata = df_input_metadata, 
    min_prevalence = 0.1,
    min_abundance = 0.001,
    max_significance = 0.05,
    output = paste0(results_dir, "maaslin2/Maaslin2_results_", data_n,""),
    fixed_effects = c("HostTypeDay"),
    reference = c("HostTypeDay,HealthyPupOral8")
)
}
```

### Box Plots

Of differentiated taxa found by Maaslin2.
We first curated the significant taxa manually by these threshold:
- qval < 0.05

Generate the box plots

```{r}
library(phyloseq)
library(ggplot2)
library(dplyr)
library(reshape2)
library(rcompanion) # For CLD
library(ggpubr) # For adding significance labels
library(FSA) # For Dunn’s test

datasets_fil_scaled_sub <- list(
  ORAL_fil = ORAL_fil_scaled,
  RECTAL_fil = RECTAL_fil_scaled
)

# Get dataset names
dataset_names <- names(datasets_fil_scaled_sub)

# Loop through each dataset
for (i in seq_along(datasets_fil_scaled_sub)) {
  data <- datasets_fil_scaled_sub[[i]]
  data_n <- dataset_names[i]

# Load Maaslin results
  maaslin_results <- read.delim(
    paste0(results_dir, "maaslin2/Maaslin2_results_", data_n, "/significant_results.tsv"),
    header = TRUE, sep = "\t", fill = TRUE, quote = "", comment.char = ""
  ) %>%
    mutate(feature = str_replace(feature, "^X\\.", " "),
           feature = str_replace_all(feature, "\\.", " "))


  # Convert ; and space into . to match the Maaslin results and create a new dataframe
maaslin_features <- as.character(maaslin_results$feature)

# make a list of the differentially abundant taxa
feature_vector <- c(" s__Clostridium_T_Clostridium_T disporicum_B", " s__Clostridium_T_Clostridium_T tertium", " s__Clostridium_T_Clostridium_T vincentii", " s__Aerococcus_Aerococcus viridans", " g__Acinetobacter")

# subset the phyloseq object based on the differentially abundant taxa
pruned_phyloseq <- subset_taxa(data, short_title %in% feature_vector)

# Skip if no taxa left after subsetting
if (ntaxa(pruned_phyloseq) == 0) {
  message("No taxa found for dataset: ", data_n, ", skipping...")
  next
}

# Get tax table with short titles
tax_info <- as.data.frame(tax_table(pruned_phyloseq)) %>%
  rownames_to_column("OTU_ID") %>%
  select(OTU_ID, short_title)

# Step 1: Create OTU table dataframe
otu_df <- as.data.frame(t(otu_table(pruned_phyloseq)))
otu_df$SampleID <- rownames(otu_df)

# Step 2: Create tax table dataframe
tax_df <- as.data.frame(tax_table(pruned_phyloseq))
tax_df$OTU_ID <- rownames(tax_df)

# Step 3: Create sample data dataframe
sample_df <- as.data.frame(sample_data(pruned_phyloseq))
sample_df$SampleID <- rownames(sample_df)

# Step 4: Melt the OTU table
library(reshape2)
melted_otu <- melt(otu_df, id.vars = "SampleID", variable.name = "OTU_ID", value.name = "Abundance")

# Step 5: Merge melted OTU with taxonomy
merged_otu_tax <- merge(melted_otu, tax_df, by = "OTU_ID", all.x = TRUE)

# Step 6: Merge with sample data
final_df <- left_join(merged_otu_tax, sample_df, by = "SampleID")

# Get all taxa names from the pruned phyloseq object
all_taxa <- unique(final_df$short_title)

for (taxon in all_taxa) {
    taxon_df <- final_df %>% filter(short_title == taxon)
    aggregated_df <- taxon_df %>%
      group_by(SampleID, HostTypeDay) %>%
      dplyr::summarize(Total_Abundance = sum(Abundance), .groups = "drop")

    aggregated_df$HostTypeDay <- factor(aggregated_df$HostTypeDay,
      levels = c("HealthyPupOral1", "HealthyPupOral8", "HealthyPupRectal1", "HealthyPupRectal8",
                 "DamMilk1", "DamMilk8", "DamOral1", "DamOral8", "DamRectal1", "DamRectal8",
                 "DamVaginal1", "DamVaginal8"))

# Prepare new colors for boxplots
# Copy original colors and override selected groups with white fill
TypeHostNew_colors_custom <- TypeHostNew_colors
groups_to_clear <- c("HealthyPupRectal1", "HealthyPupRectal8", "HealthyPupOral1", "HealthyPupOral8")
TypeHostNew_colors_custom[groups_to_clear] <- "white"  # or "transparent" for invisible fill

# Dunn's test
# Initialize log for skipped taxa
  skipped_log <- data.frame(
    Taxon = character(),
    Reason = character(),
    stringsAsFactors = FALSE
  )
 # Dunn test with error capture
 dunn_result <- tryCatch({
      dunnTest(Total_Abundance ~ HostTypeDay, data = aggregated_df, method = "bh")
    }, error = function(e) {
      skipped_log <<- rbind(skipped_log, data.frame(
        Taxon = taxon,
        Reason = paste("DunnTest error:", e$message)
      ))
      return(NULL)
    })
  
# Compact Letter Display
cld <- rcompanion::cldList(P.adj ~ Comparison, 
                data = dunn_result$res, 
                threshold = 0.05)
names(cld)[names(cld) == "Group"] <- "HostTypeDay"

# Create plot with significance letters
p <- ggplot(aggregated_df, aes(x = HostTypeDay, y = Total_Abundance,
                                   fill = HostTypeDay, color = HostTypeDay)) +
      geom_boxplot(outlier.shape = NA, alpha = 0.5, width = 0.6) +
      geom_jitter(size = 1.5, width = 0.1, alpha = 0.7) +
      geom_text(
      data = cld, 
      aes(y = max(aggregated_df$Total_Abundance) * 1.1, label = Letter),
      size = 3.5, 
      color = "black") +
      scale_fill_manual(values = TypeHostNew_colors_custom) +
      scale_color_manual(values = TypeHostNew_colors) +
      scale_shape_manual(values = TypeHostNew_shapes) +
      labs(y = "Relative Abundance [%]", x = NULL) +
      theme_minimal() +
      theme(
        legend.position = "none",
        axis.text.x = element_text(angle = 45, hjust = 1, size = 5),
        axis.title.x = element_text(size = 2.5),
        panel.grid.major = element_blank(),
        panel.grid.minor = element_blank(),
        panel.background = element_rect(fill = "white", color = "black", linewidth = 0.5)
      )

    dir.create(file.path(results_dir, "plots", "boxplots"), recursive = TRUE, showWarnings = FALSE)
jpeg_filename <- paste0(results_dir, "plots/boxplots/", data_n, "_", taxon, "_boxplot.jpeg")
jpeg(jpeg_filename, width = 6, height = 4, units = 'in', res = 300)
    print(p)
    dev.off()
    
# Save the plot as a EPS file
ggsave(
  filename = paste0(results_dir, "plots/boxplots/Selected for figure 4/EPS/",data_n, "_", taxon,"_boxplot.eps"),
  plot = p,
  device = cairo_ps,
  width = 7,
  height = 4,
  dpi = 600
)
print(p)  # Save the plot to the file
dev.off()    
  }
}
```

Run Statistical Tests 

```{r, include=FALSE}
library(openxlsx)
library(FSA)
library(rcompanion)
library(dplyr)
library(reshape2)

for (i in seq_along(datasets_fil_scaled_sub)) {
  data <- datasets_fil_scaled_sub[[i]]
  data_n <- dataset_names[i]
  existing_sheet_names <- c()

  maaslin_results <- read.delim(
    paste0(results_dir, "maaslin2/Maaslin2_results_", data_n, "/significant_results.tsv"),
    header = TRUE, sep = "\t", fill = TRUE, quote = "", comment.char = ""
  ) %>%
    mutate(feature = str_replace(feature, "^X\\.", " "),
           feature = str_replace_all(feature, "\\.", " "))


  feature_vector <- maaslin_results$feature
  pruned_phyloseq <- subset_taxa(data, short_title %in% feature_vector)

  otu_df <- as.data.frame(t(otu_table(pruned_phyloseq)))
  otu_df$SampleID <- rownames(otu_df)
  tax_df <- as.data.frame(tax_table(pruned_phyloseq))
  tax_df$OTU_ID <- rownames(tax_df)
  sample_df <- as.data.frame(sample_data(pruned_phyloseq))
  sample_df$SampleID <- rownames(sample_df)

  melted_otu <- melt(otu_df, id.vars = "SampleID", variable.name = "OTU_ID", value.name = "Abundance")
  merged_otu_tax <- merge(melted_otu, tax_df, by = "OTU_ID", all.x = TRUE)
  final_df <- left_join(merged_otu_tax, sample_df, by = "SampleID")
  all_taxa <- unique(final_df$short_title)

  wb <- createWorkbook()

  # Initialize log for skipped taxa
  skipped_log <- data.frame(
    Taxon = character(),
    Reason = character(),
    stringsAsFactors = FALSE
  )

  for (taxon in all_taxa) {
    taxon_df <- final_df %>% filter(short_title == taxon)
    aggregated_df <- taxon_df %>%
      group_by(SampleID, HostTypeDay) %>%
      dplyr::summarize(Total_Abundance = sum(Abundance), .groups = "drop")

   # Check for too few groups
    if (length(unique(aggregated_df$HostTypeDay)) < 2) {
      skipped_log <- rbind(skipped_log, data.frame(
        Taxon = taxon,
        Reason = "<2 groups in HostTypeDay"
      ))
      next
    }

    # Check for too few observations
    if (nrow(aggregated_df) < 3) {
      skipped_log <- rbind(skipped_log, data.frame(
        Taxon = taxon,
        Reason = "<3 observations"
      ))
      next
    }
    
  # Kruskal-Wallis test
    kruskal_result <- kruskal.test(Total_Abundance ~ HostTypeDay, data = aggregated_df)

   # Dunn test with error capture
    dunn_result <- tryCatch({
      dunnTest(Total_Abundance ~ HostTypeDay, data = aggregated_df, method = "bh")
    }, error = function(e) {
      skipped_log <<- rbind(skipped_log, data.frame(
        Taxon = taxon,
        Reason = paste("DunnTest error:", e$message)
      ))
      return(NULL)
    })

    if (is.null(dunn_result)) next

    base_name <- substr(make.names(taxon), 1, 25)
    base_name <- gsub("[^[:alnum:]_]", "_", base_name)

    sheet_name <- base_name
    counter <- 1
    while (tolower(sheet_name) %in% tolower(existing_sheet_names)) {
      sheet_name <- paste0(base_name, "_", counter)
      counter <- counter + 1
    }
    existing_sheet_names <- c(existing_sheet_names, sheet_name)

    # Add Kruskal
    kruskal_sheet <- substr(paste0(sheet_name, "_Kruskal"), 1, 31)
    addWorksheet(wb, kruskal_sheet)
    writeData(wb, kruskal_sheet, data.frame(
      Taxon = taxon,
      Statistic = kruskal_result$statistic,
      P.value = kruskal_result$p.value
    ))

    # Add Dunn
    dunn_sheet <- substr(paste0(sheet_name, "_Dunn"), 1, 31)
    addWorksheet(wb, dunn_sheet)
    writeData(wb, dunn_sheet, dunn_result$res)
  }

   # Add Skipped_Taxa sheet
  if (nrow(skipped_log) > 0) {
    addWorksheet(wb, "Skipped_Taxa")
    writeData(wb, "Skipped_Taxa", skipped_log)
  }

  dir.create(file.path(results_dir, "tables"), showWarnings = FALSE, recursive = TRUE)
  excel_path <- file.path(results_dir, "tables", paste0("box_plots_taxa_comparison_test_results_", data_n, ".xlsx"))
  saveWorkbook(wb, excel_path, overwrite = TRUE)
}

```

### Heatmaps

Heatmaps of log2-transformed normalized read counts for taxa that were found to be differentially abundant in Maaslin2.

```{r}
datasets_fil_scaled_sub <- list(
  MILK_fil = MILK_fil_scaled,
  PUPS_oral_fil = PUPS_oral_fil_scaled,
  PUPS_rectal_fil = PUPS_rectal_fil_scaled,
  DAMS_rectal_fil = DAMS_rectal_fil_scaled,
  ORAL_day1_fil = ORAL_day1_fil_scaled,
  ORAL_day8_fil = ORAL_day8_fil_scaled, 
  RECTAL_day1_fil = RECTAL_day1_fil_scaled,
  RECTAL_day8_fil = RECTAL_day8_fil_scaled
)

# Get dataset names
dataset_names <- names(datasets_fil_scaled_sub)

for (i in seq_along(datasets_fil_scaled_sub)) {
  data <- datasets_fil_scaled_sub[[i]]
  data_n <- dataset_names[i]
  existing_sheet_names <- c()

# Load maaslin results
maaslin_results <- read.delim(
    paste0(results_dir, "maaslin2/Maaslin2_results_", data_n, "/significant_results.tsv"),
    header = TRUE, sep = "\t", fill = TRUE, quote = "", comment.char = ""
  ) %>%
    mutate(feature = str_replace(feature, "^X\\.", " "),
           feature = str_replace_all(feature, "\\.", " "))


  # Convert ; and space into . to match the Maaslin results and create a new dataframe
maaslin_features <- as.character(maaslin_results$feature)

# Prepare metadata
df_input_metadata <- as(sample_data(data), "data.frame")

# Aggregate all levels
all_data <- aggregate_taxa(data, "short_title")
OTU1 <- as(otu_table(all_data), "matrix")
if (taxa_are_rows(all_data)) { OTU1 <- t(OTU1) }
df_input_data_a <- as.data.frame(OTU1)
df_subset <- df_input_data_a[, colnames(df_input_data_a) %in% maaslin_features, drop = FALSE]
df_subset <- t(df_subset)

# Log2 transform (add 1 to avoid log(0))
otu_table_log2 <- log2(df_subset + 1)

# Extract sample metadata
metadata <- as.data.frame(sample_data(data))
  
# Ensure metadata is properly leveled for consistent grouping
metadata$HostTypeDay <- factor(metadata$HostTypeDay,
      levels = c("HealthyPupOral1", "HealthyPupOral8", "HealthyPupRectal1", "HealthyPupRectal8",
                 "DamMilk1", "DamMilk8", "DamOral1", "DamOral8", "DamRectal1", "DamRectal8",
                 "DamVaginal1", "DamVaginal8"))

# Create annotation for samples (HostTypeDay)
annotation_col <- data.frame(HostTypeDay = metadata$HostTypeDay)
rownames(annotation_col) <- rownames(metadata)

# Assign colors
ann_colors <- list(HostTypeDay = TypeHostNew_colors)

# Skip if there is only 1 differentiated taxon
  if (nrow(otu_table_log2) < 2) {
    message("Only one taxon found for ", data_n, "; skipping heatmap.")
    next
  }

# Generate heatmap
tiff(filename = paste0(results_dir,"plots/heatmaps/",data_n,"_heatmap_Maaslin2_sig_results.tiff"), width=12, height=6, units='in', res=300)
heatmap <- pheatmap(otu_table_log2, 
           cluster_rows = TRUE, 
           cluster_cols = TRUE, 
           scale = "row", 
           show_rownames = FALSE,
           annotation_colors = ann_colors, 
           color = colorRampPalette(c("blue", "white", "red"))(50),
           main = paste("Heatmap of Log2 Normalized Abundances -", data_n))
print(heatmap)
dev.off()

# Save the plot as a EPS file
ggsave(
  filename = paste0(results_dir, "plots/heatmaps/EPS/",data_n, "_heatmap_Maaslin2_sig_results.eps"),
  plot = heatmap,
  device = cairo_ps,
  width = 7,
  height = 4,
  dpi = 600
)
print(heatmap)  # Save the plot to the file
dev.off()    

}


```

## Picrust2

Prepare the files for running picrust2 in linux

```{r}
library(phyloseq)
library(biomformat)
library(Biostrings)

# Extract the OTU/ASV table
otu_table_filtered_all <- otu_table(ALL_fil)

# Convert to biom format
biom_file_all <- make_biom(data = as.matrix(otu_table_filtered_all))

# Piscrust directory
pi_dir <- paste0(getwd(), "/Picrust2/data/")

# Save as a BIOM file
write_biom(biom_file_all, paste0(pi_dir,"filtered_table_2025.biom"))

# Ensure names match ASV IDs
names(seqs) <- sub("^>", "", names(seqs))  # Remove FASTA '>' if needed

# Add sequences to the phyloseq object
phy_filtered <- merge_phyloseq(ALL_fil, seqs)

# Extract ASV sequences
asv_seqs <- refseq(phy_filtered)

# Clean up ASV sequences (remove only "-" characters)
asv_clean <- DNAStringSet(gsub("-", "", as.character(asv_seqs)))

# Reassign names
names(asv_clean) <- names(asv_seqs)

# Save to FASTA
writeXStringSet(asv_clean, filepath = file.path(pi_dir, "ALL_fil_seqs_2025_cleaned.fna"))

# Save to FASTA
writeXStringSet(seqs_filtered, filepath = paste0(pi_dir,"ALL_fil_seqs_2025.fna"))

```

*** AFTER THE PICRUST2 RUN ON THE SERVER ***

### Load libraries

```{r, include=FALSE}
require(ggpicrust2)
require(readr)
require(edgeR)
library(ggh4x)
library(textshape)
library(dplyr)
require(tidyr)
library(patchwork)
library(tidyverse)
library(Maaslin2)
library(ggplot2)
library(ggprism)
```

### Load metadata

```{r}

metadata_all <- read_delim(paste0(getwd(),"/Picrust2/data/Puppies_Picrust2_metadata.txt"),
    delim = "\t",
    escape_double = FALSE,
    trim_ws = TRUE,
    col_types = cols(SampleID = col_character())  # Explicitly set SampleID as character
)
```

Subset the metadata for downstream comparisons

```{r}
metadata_Dams <- metadata_all %>% dplyr::filter(Host=="Dam")
metadata_Pups <- metadata_all %>% dplyr::filter(Host=="HealthyPup")

metadata_Dams_Milk <- metadata_all %>%
  dplyr::filter(Host=="Dam" & Type %in% c("Milk"))
metadata_Dams_Oral <- metadata_all %>%
  dplyr::filter(Host=="Dam" & Type %in% c("Oral"))
metadata_Dams_Rectal <- metadata_all %>%
  dplyr::filter(Host=="Dam" & Type %in% c("Rectal"))
metadata_Dams_Vaginal <- metadata_all %>%
  dplyr::filter(Host=="Dam" & Type %in% c("Vaginal"))


metadata_Pups_Oral <- metadata_all %>%
  dplyr::filter(Host=="HealthyPup" & Type %in% c("Oral"))
metadata_Pups_Rectal <- metadata_all %>%
  dplyr::filter(Host=="HealthyPup" & Type %in% c("Rectal"))

metadata_Oral_day1 <- metadata_all %>%
  dplyr::filter(Day == "1" & Type %in% c( "Oral"))
metadata_Oral_day8 <- metadata_all %>%
  dplyr::filter(Day == "8" & Type %in% c( "Oral"))
metadata_Rectal_day1 <- metadata_all %>%
  dplyr::filter(Day == "1" & Type %in% c( "Rectal"))
metadata_Rectal_day8 <- metadata_all %>%
  dplyr::filter(Day == "8" & Type %in% c( "Rectal"))

metadata_All_day1 <- metadata_all %>%
  dplyr::filter(Day == "1")
metadata_All_day8 <- metadata_all %>%
  dplyr::filter(Day == "8")

metadata_All_day1_no_rectal <- metadata_all %>%
  dplyr::filter(Day == "1"& Type %in% c( "Milk", "Oral", "Vaginal"))
metadata_All_day8_no_rectal <- metadata_all %>%
  dplyr::filter(Day == "8"& Type %in% c( "Milk", "Oral", "Vaginal"))

```

Load KO input file

```{r}
#KO abudnace file
ko_abun <-
  read_delim(paste0(getwd(),"/Picrust2/results/Remote server/KO_metagenome_out/pred_metagenome_unstrat.tsv"),
    delim = "\t",
    escape_double = FALSE,
    trim_ws = TRUE
  )
ko_abun$`function` <- gsub("ko:", "", ko_abun$`function`)

```

Convert KO to KEGG

```{r}
kegg_abundance <- ggpicrust2::ko2kegg_abundance(data = ko_abun)
```

Differential abundance of KEGG

[1]
Pups only (comparison between days)

```{r, include=FALSE}
#Define variables
daa_method <- "KEGG"

# Define data set
picrust_pups <- list(metadata_Pups_Oral = metadata_Pups_Oral,
                      metadata_Pups_Rectal = metadata_Pups_Rectal)

# Get dataset names
dataset_names <- names(picrust_pups)

# Loop through each dataset
for (i in 1:length(picrust_pups)) {
  # Set current dataset and its name
  data <- picrust_pups[[i]]
  data_n <- dataset_names[i]
  
# [1]pathway_daa()

## Prepare the files
# convert metadata to data.frame if needed
metadata_mas <- as.data.frame(data) 
# make SampleID row names in the metadata file
rownames(metadata_mas) <- metadata_mas$SampleID
# Keep only the samples in the current metadata
# Use sample IDs that exist in both metadata and kegg_abundance
common_samples <- intersect(metadata_mas$SampleID, colnames(kegg_abundance))
# Subset metadata and abundance to common samples
metadata_mas <- metadata_mas[metadata_mas$SampleID %in% common_samples, , drop = FALSE]
kegg_abundance_sub <- kegg_abundance[, common_samples, drop = FALSE]
# transpose to fit the format of Maaslin2
kegg_abundance_t <- t(kegg_abundance_sub)

# Define output path for this dataset
output_path <- paste0(results_dir, "Picrust_Maaslin2/Picrust_Maaslin2_", data_n)

# Make sure the directory exists
if (!dir.exists(output_path)) {
  dir.create(output_path, recursive = TRUE)
}

maaslin_results <- Maaslin2(
  input_data = kegg_abundance_t, 
  input_metadata = metadata_mas, 
  min_prevalence = 0.1,
  min_abundance = 0.001,
  output = output_path,
  fixed_effects = c("Day"),
  random_effects = c("Mother"),
  reference = c("Day,Day 1")
)

# Convert results for ggpicrust2
formatted_results <- data.frame(
  feature = maaslin_results$results$feature,
  method = "Maaslin2",
  group1 = "Day 1",
  group2 = "Day 8",
  p_values = maaslin_results$results$pval,
  q_values = maaslin_results$results$qval,
  coef = maaslin_results$results$coef,
  stderr = maaslin_results$results$stderr
)

# Replace _ with - in pathway name for MetaCyc
formatted_results$feature <- gsub("\\.", "-", formatted_results$feature)
formatted_results$p_adjust <- formatted_results$q_values

# [2] pathway_annotation()
kegg_abun_annotated_kegg_daa_results_df <- pathway_annotation(pathway = "KO",
daa_results_df = formatted_results, ko_to_kegg = T)

# [3] pathway_heatmap()
#filter for only p-value <0.05
kegg_feature_with_p_0.05 <- formatted_results %>% filter(p_adjust < 0.05)

#filter the 'abundance' file based on the values <0.05 and remove the 'pathway' column so that the heatmap function can be run
kegg_abun<-kegg_abundance_sub
kegg_abun$functions <- rownames(kegg_abun)  #add a column to join
kegg_abun2 <- kegg_abun %>%
right_join(kegg_abun_annotated_kegg_daa_results_df %>%
select(all_of(c("feature","pathway_class"))), by = c("functions" = "feature")) %>%
filter(functions %in% kegg_feature_with_p_0.05$feature) 
kegg_abun3 <- kegg_abun2 %>%
    mutate(pathway_class = ifelse(is.na(pathway_class), functions, pathway_class))
kegg_abun4 <- kegg_abun3 %>% select(-"functions")
#for comparisons with >2 groups: remove randomly two rows out of each threesome and then convert the 'description' column to row names because otherwise we can't run the heatmap function 
kegg_abun5 <- kegg_abun4 %>% distinct(pathway_class, .keep_all = T) %>% column_to_rownames("pathway_class")

#export the image
tiff(filename = paste0(getwd(),"/Picrust2/results/R/figures/dark/",data_n,"_", daa_method, "_reference_Day1_qval_0.05_heatmap.tiff"), width=16, height=12, pointsize = 1/300, units='in', res=300)
#create a heatmap
heatmap_plot <- ggpicrust2::pathway_heatmap(abundance = kegg_abun5, metadata_mas, group = "Day", show_row_names = T, font_size = 10)
print(heatmap_plot)
dev.off()

#export the table
write.csv(kegg_feature_with_p_0.05, file = paste0(getwd(),"/Picrust2/results/R/tables/",data_n,"_",daa_method,"_reference_Day1_qval_0.05.csv"))


# [4] pathway_errorbar()

SampleID<-as.data.frame(colnames(kegg_abundance_sub))
metadata2 <- right_join(metadata_mas, SampleID, by=c("SampleID"="colnames(kegg_abundance_sub)"))
# Limit to the Top 30 features
low_p_feature <- formatted_results[order(formatted_results$p_adjust), ]$feature[1:30]

#for comparisons with >2 groups: remove randomly two rows out of each threesome and then convert the 'description' column to row names because otherwise we can't run the heatmap function 
kegg_daa_annotated_results_df2 <- formatted_results %>% filter(group1 == "Day 1")  %>% distinct(feature, .keep_all = T) 

kegg_abun6 <- kegg_abun4 %>% distinct(pathway_class, .keep_all = T) 

if ((sum(kegg_daa_annotated_results_df2$p_adjust<=0.05)<=3)) {
  next
}


#export the image
tiff(filename = paste0(getwd(),"/Picrust2/results/R/figures/dark/",data_n,"_",daa_method,"_Day1_KEGG_qval_0.05_errorbar.tiff"), width=16, height=12, pointsize = 1/300, units='in', res=300)
kegg_error <- pathway_errorbar(abundance = kegg_abundance_sub,
                      daa_results_df = kegg_abun_annotated_kegg_daa_results_df,
                      Group = metadata2$Day,
                      ko_to_kegg = T,
                      p_values_threshold = 0.05,
                      order = "pathway_class",
                      select = low_p_feature,
                      p_value_bar = F,
                      colors = NULL,
                      x_lab = "pathway_name")
print(kegg_error)
dev.off()
 

# [5] pathway_pca()

metadata3 <- metadata2
metadata3$sample_name <- metadata3$SampleID
# Remove pathways with zero variance across samples
kegg_abundance_filtered <- kegg_abun5[apply(kegg_abun5, 1, function(x) var(x) != 0), ]

# PCoA with more control over colors and shapes
# Define colors
TypeHostNew_colors_pi_dams <- c(
    "1" = "cornflowerblue",
    "8" = "blue2"
)

# Run PCA manually (example with prcomp, adjust as needed)
pca_result <- prcomp(t(kegg_abundance_filtered), scale. = TRUE)
pca_scores <- as.data.frame(pca_result$x)
pca_scores$Day <- as.factor(metadata3$Day)  # make sure order matches
pca_var <- pca_result$sdev^2
pca_var_perc <- round(100 * pca_var / sum(pca_var), 1)


tiff(filename = paste0(getwd(),"/Picrust2/results/R/figures/",data_n,"_",daa_method,"_KEGG_qval_0.05_pca_ggplot2.tiff"), width=6, height=4, pointsize = 1/300, units='in', res=300)
picrust_pca_colordams <- ggplot(pca_scores, aes(x = PC1, y = PC2, color = Day)) +
  geom_point(shape = 15,  size = 4, 
             stroke = 2) +   # increase thickness of marker borders) +  
  stat_ellipse(aes(group = Day), linewidth = 1) +  # Add 95% ellipses
  scale_color_manual(values = TypeHostNew_colors_pi_dams) +
  labs(
  x = sprintf("PC1 (%.1f%%)", pca_var_perc[1]),
  y = sprintf("PC2 (%.1f%%)", pca_var_perc[2])
  ) +
  theme_classic() +
  theme(
    legend.text = element_text(size = 15),
    legend.title = element_text(size = 15),
    legend.key.size = unit(0.5, "cm")
  )
print(picrust_pca_colordams)
dev.off()

# Save the plot as a EPS file
ggsave(
  filename = paste0(getwd(),"/Picrust2/results/R/figures/",data_n,"_",daa_method,"_KEGG_qval_0.05_pca_ggplot2.eps"),
  plot = picrust_pca_colordams,
  device = cairo_ps,
  width = 7,
  height = 4,
  dpi = 600
)
print(picrust_pca_colordams)  # Save the plot to the file
dev.off()   

}
```

[2]
Dams only (comparison between days)

```{r, include=FALSE}
#Define variables
daa_method <- "KEGG"

# Define data set
picrust_set <- list(metadata_Dams_Rectal = metadata_Dams_Rectal)

# Get dataset names
dataset_names <- names(picrust_set)

# Loop through each dataset
for (i in 1:length(picrust_set)) {
  # Set current dataset and its name
  data <- picrust_set[[i]]
  data_n <- dataset_names[i]
  
# [1]pathway_daa()

## Prepare the files
# convert metadata to data.frame if needed
metadata_mas <- as.data.frame(data) 
# make SampleID row names in the metadata file
rownames(metadata_mas) <- metadata_mas$SampleID
# Keep only the samples in the current metadata
# Use sample IDs that exist in both metadata and kegg_abundance
common_samples <- intersect(metadata_mas$SampleID, colnames(kegg_abundance))
# Subset metadata and abundance to common samples
metadata_mas <- metadata_mas[metadata_mas$SampleID %in% common_samples, , drop = FALSE]
kegg_abundance_sub <- kegg_abundance[, common_samples, drop = FALSE]
# transpose to fit the format of Maaslin2
kegg_abundance_t <- t(kegg_abundance_sub)

# Define output path for this dataset
output_path <- paste0(results_dir, "Picrust_Maaslin2/Picrust_Maaslin2_", data_n)

# Make sure the directory exists
if (!dir.exists(output_path)) {
  dir.create(output_path, recursive = TRUE)
}

maaslin_results <- Maaslin2(
  input_data = kegg_abundance_t, 
  input_metadata = metadata_mas, 
  min_prevalence = 0.1,
  min_abundance = 0.001,
  output = paste0(results_dir, "Picrust_Maaslin2/Picrust_Maaslin2_",data_n),
  fixed_effects = c("Day"),
  reference = c("Day,Day 1")
)

# Convert results for ggpicrust2
formatted_results <- data.frame(
  feature = maaslin_results$results$feature,
  method = "Maaslin2",
  group1 = "Day 1",
  group2 = maaslin_results$results$value,
  p_values = maaslin_results$results$pval,
  q_values = maaslin_results$results$qval,
  coef = maaslin_results$results$coef,
  stderr = maaslin_results$results$stderr
)

# Replace _ with - in pathway name for MetaCyc
formatted_results$feature <- gsub("\\.", "-", formatted_results$feature)
formatted_results$p_adjust <- formatted_results$q_values

# SKIP if there are no significant features BEFORE calling pathway_annotation
if (sum(formatted_results$p_adjust < 0.05) == 0) {
  message(paste0("Skipping ", data_n, " — no significant features (p_adjust < 0.05)."))
  next
}

# [2] pathway_annotation()
kegg_abun_annotated_kegg_daa_results_df <- pathway_annotation(pathway = "KO",
daa_results_df = formatted_results, ko_to_kegg = T)

# [3] pathway_heatmap()
#filter for only p-value <0.05
kegg_feature_with_p_0.05 <- formatted_results %>% filter(p_adjust < 0.05)

#filter the 'abundance' file based on the values <0.05 and remove the 'pathway' column so that the heatmap function can be run
kegg_abun<-kegg_abundance_sub
kegg_abun$functions <- rownames(kegg_abun)  #add a column to join
kegg_abun2 <- kegg_abun %>%
right_join(kegg_abun_annotated_kegg_daa_results_df %>%
select(all_of(c("feature","pathway_class"))), by = c("functions" = "feature")) %>%
filter(functions %in% kegg_feature_with_p_0.05$feature) 
kegg_abun3 <- kegg_abun2 %>%
    mutate(pathway_class = ifelse(is.na(pathway_class), functions, pathway_class))
kegg_abun4 <- kegg_abun3 %>% select(-"functions")
#for comparisons with >2 groups: remove randomly two rows out of each threesome and then convert the 'description' column to row names because otherwise we can't run the heatmap function 
kegg_abun5 <- kegg_abun4 %>% distinct(pathway_class, .keep_all = T) %>% column_to_rownames("pathway_class")

#export the image
tiff(filename = paste0(getwd(),"/Picrust2/results/R/figures/",data_n,"_", daa_method, "_reference_Day1_qval_0.05_heatmap.tiff"), width=16, height=12, pointsize = 1/300, units='in', res=300)
#create a heatmap
heatmap_plot <- ggpicrust2::pathway_heatmap(abundance = kegg_abun5, metadata_mas, group = "Day", show_row_names = T, font_size = 10)
print(heatmap_plot)
dev.off()

#export the table
write.csv(kegg_feature_with_p_0.05, file = paste0(getwd(),"/Picrust2/results/R/tables/",data_n,"_",daa_method,"_reference_Day1_qval_0.05.csv"))


# [4] pathway_errorbar()

SampleID<-as.data.frame(colnames(kegg_abundance_sub))
metadata2 <- right_join(metadata_mas, SampleID, by=c("SampleID"="colnames(kegg_abundance_sub)"))
# Limit to the Top 30 features
low_p_feature <- formatted_results[order(formatted_results$p_adjust), ]$feature[1:30]

kegg_abun6 <- kegg_abun4 %>% distinct(pathway_class, .keep_all = T) 

# [5] pathway_pca()

metadata3 <- metadata2
metadata3$sample_name <- metadata3$SampleID
# Remove pathways with zero variance across samples
kegg_abundance_filtered <- kegg_abun5[apply(kegg_abun5, 1, function(x) var(x) != 0), ]

# PCoA with more control over colors and shapes
# Define colors
TypeHostNew_colors_pi_dams <- c(
    "1" = "cornflowerblue",
    "8" = "blue2"
)

# Run PCA manually (example with prcomp, adjust as needed)
pca_result <- prcomp(t(kegg_abundance_filtered), scale. = TRUE)
pca_scores <- as.data.frame(pca_result$x)
pca_scores$Day <- as.factor(metadata3$Day)  # make sure order matches
pca_var <- pca_result$sdev^2
pca_var_perc <- round(100 * pca_var / sum(pca_var), 1)


tiff(filename = paste0(getwd(),"/Picrust2/results/R/figures/",data_n,"_",daa_method,"_KEGG_qval_0.05_pca_ggplot2.tiff"), width=6, height=4, pointsize = 1/300, units='in', res=300)
picrust_pca_colordams <- ggplot(pca_scores, aes(x = PC1, y = PC2, color = Day)) +
  geom_point(shape = 15,  size = 4, 
             stroke = 2) +   # increase thickness of marker borders) +  
  stat_ellipse(aes(group = Day), linewidth = 1) +  # Add 95% ellipses
  scale_color_manual(values = TypeHostNew_colors_pi_dams) +
  labs(
  x = sprintf("PC1 (%.1f%%)", pca_var_perc[1]),
  y = sprintf("PC2 (%.1f%%)", pca_var_perc[2])
  ) +
  theme_classic() +
  theme(
    legend.text = element_text(size = 15),
    legend.title = element_text(size = 15),
    legend.key.size = unit(0.5, "cm")
  )
print(picrust_pca_colordams)
dev.off()

# Save the plot as a EPS file
ggsave(
  filename = paste0(getwd(),"/Picrust2/results/R/figures/",data_n,"_",daa_method,"_KEGG_qval_0.05_pca_ggplot2.eps"),
  plot = picrust_pca_colordams,
  device = cairo_ps,
  width = 7,
  height = 4,
  dpi = 600
)
print(picrust_pca_colordams)  # Save the plot to the file
dev.off()   

}
```

[3]
Pups/dams oral/rectal comparison (per specific day)

```{r, include=FALSE}
#Define variables
daa_method <- "KEGG"

# Define data set
picrust_set <- list(metadata_Rectal_day1 = metadata_Rectal_day1,
                    metadata_Rectal_day8 = metadata_Rectal_day8,
                    metadata_Oral_day1 = metadata_Oral_day1,
                    metadata_Oral_day8 = metadata_Oral_day8)

# Get dataset names
dataset_names <- names(picrust_set)

# Loop through each dataset
for (i in 1:length(picrust_set)) {
  # Set current dataset and its name
  data <- picrust_set[[i]]
  data_n <- dataset_names[i]
  
# [1]pathway_daa()

## Prepare the files
# convert metadata to data.frame if needed
metadata_mas <- as.data.frame(data) 
# make SampleID row names in the metadata file
rownames(metadata_mas) <- metadata_mas$SampleID
# Keep only the samples in the current metadata
# Use sample IDs that exist in both metadata and kegg_abundance
common_samples <- intersect(metadata_mas$SampleID, colnames(kegg_abundance))
# Subset metadata and abundance to common samples
metadata_mas <- metadata_mas[metadata_mas$SampleID %in% common_samples, , drop = FALSE]
kegg_abundance_sub <- kegg_abundance[, common_samples, drop = FALSE]
# transpose to fit the format of Maaslin2
kegg_abundance_t <- t(kegg_abundance_sub)

# Define output path for this dataset
output_path <- paste0(results_dir, "Picrust_Maaslin2/Picrust_Maaslin2_", data_n)

# Make sure the directory exists
if (!dir.exists(output_path)) {
  dir.create(output_path, recursive = TRUE)
}

maaslin_results <- Maaslin2(
  input_data = kegg_abundance_t, 
  input_metadata = metadata_mas, 
  min_prevalence = 0.1,
  min_abundance = 0.001,
  output = paste0(results_dir, "Picrust_Maaslin2/Picrust_Maaslin2_",data_n),
  fixed_effects = c("Host"),
  reference = c("Host,Dam")
)

# Convert results for ggpicrust2
formatted_results <- data.frame(
  feature = maaslin_results$results$feature,
  method = "Maaslin2",
  group1 = "Dam",
  group2 = maaslin_results$results$value,
  p_values = maaslin_results$results$pval,
  q_values = maaslin_results$results$qval,
  coef = maaslin_results$results$coef,
  stderr = maaslin_results$results$stderr
)

# Replace _ with - in pathway name for MetaCyc
formatted_results$feature <- gsub("\\.", "-", formatted_results$feature)
formatted_results$p_adjust <- formatted_results$q_values

# [2] pathway_annotation()
kegg_abun_annotated_kegg_daa_results_df <- pathway_annotation(pathway = "KO",
daa_results_df = formatted_results, ko_to_kegg = T)

# [3] pathway_heatmap()
#filter for only p-value <0.05
kegg_feature_with_p_0.05 <- formatted_results %>% filter(p_adjust < 0.05)

#filter the 'abundance' file based on the values <0.05 and remove the 'pathway' column so that the heatmap function can be run
kegg_abun<-kegg_abundance_sub
kegg_abun$functions <- rownames(kegg_abun)  #add a column to join
kegg_abun2 <- kegg_abun %>%
right_join(kegg_abun_annotated_kegg_daa_results_df %>%
select(all_of(c("feature","pathway_class"))), by = c("functions" = "feature")) %>%
filter(functions %in% kegg_feature_with_p_0.05$feature) 
kegg_abun3 <- kegg_abun2 %>%
    mutate(pathway_class = ifelse(is.na(pathway_class), functions, pathway_class))
kegg_abun4 <- kegg_abun3 %>% select(-"functions")
#for comparisons with >2 groups: remove randomly two rows out of each threesome and then convert the 'description' column to row names because otherwise we can't run the heatmap function 
kegg_abun5 <- kegg_abun4 %>% distinct(pathway_class, .keep_all = T) %>% column_to_rownames("pathway_class")

#export the image
tiff(filename = paste0(getwd(),"/Picrust2/results/R/figures/",data_n,"_", daa_method, "_reference_Day1_qval_0.05_heatmap.tiff"), width=16, height=12, pointsize = 1/300, units='in', res=300)
#create a heatmap
heatmap_plot <- ggpicrust2::pathway_heatmap(abundance = kegg_abun5, metadata_mas, group = "Host", show_row_names = T, font_size = 10)
print(heatmap_plot)
dev.off()

#export the table
write.csv(kegg_feature_with_p_0.05, file = paste0(getwd(),"/Picrust2/results/R/tables/",data_n,"_",daa_method,"_reference_Day1_qval_0.05.csv"))


# [4] pathway_errorbar()

SampleID<-as.data.frame(colnames(kegg_abundance_sub))
metadata2 <- right_join(metadata_mas, SampleID, by=c("SampleID"="colnames(kegg_abundance_sub)"))
# Limit to the Top 30 features
low_p_feature <- formatted_results[order(formatted_results$p_adjust), ]$feature[1:30]

kegg_abun6 <- kegg_abun4 %>% distinct(pathway_class, .keep_all = T) 

# [5] pathway_pca()

metadata3 <- metadata2
metadata3$sample_name <- metadata3$SampleID
# Remove pathways with zero variance across samples
kegg_abundance_filtered <- kegg_abun5[apply(kegg_abun5, 1, function(x) var(x) != 0), ]

# PCoA with more control over colors and shapes
# Define colors
TypeHostNew_colors_pi_day1 <- c(
    "HealthyPupOral"   = "gold1",  
    "DamMilk" = "#66C2A5",  
    "DamVaginal"   = "mediumorchid",  
    "DamOral"   = "gold1"
)
TypeHostNew_colors_pi_day8 <- c(
    "HealthyPupOral"   = "darkgoldenrod4",  
    "DamMilk" = "#117733",  
    "DamVaginal"   = "darkorchid4",  
    "DamOral"   = "darkgoldenrod4"
)

TypeHostNew_shapes_pi <- c(
    "HealthyPupOral"   = 22,  
    "DamMilk" = 15,           
    "DamVaginal" = 15,        
    "DamOral" = 15          
)

#  Run PCA manually (example with prcomp, adjust as needed)
pca_result <- prcomp(t(kegg_abundance_filtered), scale. = TRUE)
pca_scores <- as.data.frame(pca_result$x)
pca_scores$TypeHost <- metadata3$TypeHost  # make sure order matches
pca_var <- pca_result$sdev^2
pca_var_perc <- round(100 * pca_var / sum(pca_var), 1)


picrust_pca_colorday1 <- ggplot(pca_scores, aes(x = PC1, y = PC2, color = TypeHost, shape = TypeHost)) +
  geom_point(aes(shape = TypeHost), size = 4, fill = "white", 
             stroke = 2) +   # increase thickness of marker borders) +  
  scale_color_manual(values = TypeHostNew_colors_pi_day1) +
  scale_shape_manual(values = TypeHostNew_shapes_pi) +
  labs(
  x = sprintf("PC1 (%.1f%%)", pca_var_perc[1]),
  y = sprintf("PC2 (%.1f%%)", pca_var_perc[2])
  ) +
  theme_classic() +
  theme(
    legend.position = "none"
  )

print(picrust_pca_colorday1)

picrust_pca_colorday8 <- ggplot(pca_scores, aes(x = PC1, y = PC2, color = TypeHost, shape = TypeHost)) +
  geom_point(aes(shape = TypeHost), size = 4, fill = "white", 
             stroke = 2) +   # increase thickness of marker borders) +  
  scale_color_manual(values = TypeHostNew_colors_pi_day8) +
  scale_shape_manual(values = TypeHostNew_shapes_pi) +
  labs(
  x = sprintf("PC1 (%.1f%%)", pca_var_perc[1]),
  y = sprintf("PC2 (%.1f%%)", pca_var_perc[2])
  ) +
  theme_classic() +
  theme(
    legend.position = "none"
  )

print(picrust_pca_colorday8)

# Save the plot as a EPS file
ggsave(
  filename = paste0(getwd(),"/Picrust2/results/R/figures/dark/",data_n,"_",daa_method,"_KEGG_qval_0.05_pca_day1_colors.eps"),
  plot = picrust_pca_colorday1,
  device = cairo_ps,
  width = 7,
  height = 4,
  dpi = 600
)
print(picrust_pca_colorday1)  # Save the plot to the file
dev.off()  

# Save the plot as a EPS file
ggsave(
  filename = paste0(getwd(),"/Picrust2/results/R/figures/dark/",data_n,"_",daa_method,"_KEGG_qval_0.05_pca_day8_colors.eps"),
  plot = picrust_pca_colorday8,
  device = cairo_ps,
  width = 7,
  height = 4,
  dpi = 600
)
print(picrust_pca_colorday8)  # Save the plot to the file
dev.off()  
}
```

[4]
Pups/dams comparison all body sites (per specific day)

```{r, include=FALSE}
#Define variables
daa_method <- "KEGG"

# Define data set
picrust_set <- list(metadata_All_day1 = metadata_All_day1,
                    metadata_All_day8 = metadata_All_day8)

# Get dataset names
dataset_names <- names(picrust_set)

# Loop through each dataset
for (i in 1:length(picrust_set)) {
  # Set current dataset and its name
  data <- picrust_set[[i]]
  data_n <- dataset_names[i]
  
# [1]pathway_daa()

## Prepare the files
# convert metadata to data.frame if needed
metadata_mas <- as.data.frame(data) 
# make SampleID row names in the metadata file
rownames(metadata_mas) <- metadata_mas$SampleID
# Keep only the samples in the current metadata
# Use sample IDs that exist in both metadata and kegg_abundance
common_samples <- intersect(metadata_mas$SampleID, colnames(kegg_abundance))
# Subset metadata and abundance to common samples
metadata_mas <- metadata_mas[metadata_mas$SampleID %in% common_samples, , drop = FALSE]
kegg_abundance_sub <- kegg_abundance[, common_samples, drop = FALSE]
# transpose to fit the format of Maaslin2
kegg_abundance_t <- t(kegg_abundance_sub)

maaslin_results <- Maaslin2(
  input_data = kegg_abundance_t, 
  input_metadata = metadata_mas, 
  min_prevalence = 0.1,
  min_abundance = 0.001,
  output = paste0(results_dir, "Picrust_Maaslin2/Picrust_Maaslin2_",data_n),
  fixed_effects = c("TypeHost"),
  reference = c("TypeHost,DamMilk")
)

# Convert results for ggpicrust2
formatted_results <- data.frame(
  feature = maaslin_results$results$feature,
  method = "Maaslin2",
  group1 = "Dam",
  group2 = maaslin_results$results$value,
  p_values = maaslin_results$results$pval,
  q_values = maaslin_results$results$qval,
  coef = maaslin_results$results$coef,
  stderr = maaslin_results$results$stderr
)

# Replace _ with - in pathway name for MetaCyc
formatted_results$feature <- gsub("\\.", "-", formatted_results$feature)
formatted_results$p_adjust <- formatted_results$q_values

# [2] pathway_annotation()
kegg_abun_annotated_kegg_daa_results_df <- pathway_annotation(pathway = "KO",
daa_results_df = formatted_results, ko_to_kegg = T)

# [3] pathway_heatmap()
#filter for only p-value <0.05
kegg_feature_with_p_0.05 <- formatted_results %>% filter(p_adjust < 0.05)

#filter the 'abundance' file based on the values <0.05 and remove the 'pathway' column so that the heatmap function can be run
kegg_abun<-kegg_abundance_sub
kegg_abun$functions <- rownames(kegg_abun)  #add a column to join
kegg_abun2 <- kegg_abun %>%
right_join(kegg_abun_annotated_kegg_daa_results_df %>%
select(all_of(c("feature","pathway_class"))), by = c("functions" = "feature")) %>%
filter(functions %in% kegg_feature_with_p_0.05$feature) 
kegg_abun3 <- kegg_abun2 %>%
    mutate(pathway_class = ifelse(is.na(pathway_class), functions, pathway_class))
kegg_abun4 <- kegg_abun3 %>% select(-"functions")
#for comparisons with >2 groups: remove randomly two rows out of each threesome and then convert the 'description' column to row names because otherwise we can't run the heatmap function 
kegg_abun5 <- kegg_abun4 %>% distinct(pathway_class, .keep_all = T) %>% column_to_rownames("pathway_class")

#export the image
tiff(filename = paste0(getwd(),"/Picrust2/results/R/figures/dark/",data_n,"_", daa_method, "_reference_Day1_qval_0.05_heatmap.tiff"), width=16, height=12, pointsize = 1/300, units='in', res=300)
#create a heatmap
heatmap_plot <- ggpicrust2::pathway_heatmap(abundance = kegg_abun5, metadata_mas, group = "TypeHost", show_row_names = T, font_size = 10)
print(heatmap_plot)
dev.off()

#export the table
write.csv(kegg_feature_with_p_0.05, file = paste0(getwd(),"/Picrust2/results/R/tables/",data_n,"_",daa_method,"_reference_Day1_qval_0.05.csv"))


# [4] pathway_errorbar()

SampleID<-as.data.frame(colnames(kegg_abundance_sub))
metadata2 <- right_join(metadata_mas, SampleID, by=c("SampleID"="colnames(kegg_abundance_sub)"))
# Limit to the Top 30 features
low_p_feature <- formatted_results[order(formatted_results$p_adjust), ]$feature[1:30]

kegg_abun6 <- kegg_abun4 %>% distinct(pathway_class, .keep_all = T) 

# [5] pathway_pca()

metadata3 <- metadata2
metadata3$sample_name <- metadata3$SampleID
# Remove pathways with zero variance across samples
kegg_abundance_filtered <- kegg_abun5[apply(kegg_abun5, 1, function(x) var(x) != 0), ]

# PCoA with more control over colors and shapes
# Define colors
TypeHostNew_colors_pi_day1 <- c(
    "HealthyPupOral"   = "gold1",  
    "DamMilk" = "#66C2A5",  
    "DamVaginal"   = "mediumorchid",  
    "DamOral"   = "gold1"
)
TypeHostNew_colors_pi_day8 <- c(
    "HealthyPupOral"   = "darkgoldenrod4",  
    "DamMilk" = "#117733",  
    "DamVaginal"   = "darkorchid4",  
    "DamOral"   = "darkgoldenrod4"
)

TypeHostNew_shapes_pi <- c(
    "HealthyPupOral"   = 22,  
    "DamMilk" = 15,           
    "DamVaginal" = 15,        
    "DamOral" = 15          
)

# Run PCA manually (example with prcomp, adjust as needed)
pca_result <- prcomp(t(kegg_abundance_filtered), scale. = TRUE)
pca_scores <- as.data.frame(pca_result$x)
pca_scores$TypeHost <- metadata3$TypeHost  # make sure order matches
pca_var <- pca_result$sdev^2
pca_var_perc <- round(100 * pca_var / sum(pca_var), 1)


picrust_pca_colorday1 <- ggplot(pca_scores, aes(x = PC1, y = PC2, color = TypeHost, shape = TypeHost)) +
  geom_point(aes(shape = TypeHost), size = 4, fill = "white", 
             stroke = 2) +   # increase thickness of marker borders) +  
  scale_color_manual(values = TypeHostNew_colors_pi_day1) +
  scale_shape_manual(values = TypeHostNew_shapes_pi) +
  labs(
  x = sprintf("PC1 (%.1f%%)", pca_var_perc[1]),
  y = sprintf("PC2 (%.1f%%)", pca_var_perc[2])
  ) +
  theme_classic() +
  theme(
    legend.position = "none"
  )

print(picrust_pca_colorday1)

picrust_pca_colorday8 <- ggplot(pca_scores, aes(x = PC1, y = PC2, color = TypeHost, shape = TypeHost)) +
  geom_point(aes(shape = TypeHost), size = 4, fill = "white", 
             stroke = 2) +   # increase thickness of marker borders) +  
  scale_color_manual(values = TypeHostNew_colors_pi_day8) +
  scale_shape_manual(values = TypeHostNew_shapes_pi) +
  labs(
  x = sprintf("PC1 (%.1f%%)", pca_var_perc[1]),
  y = sprintf("PC2 (%.1f%%)", pca_var_perc[2])
  ) +
  theme_classic() +
  theme(
    legend.position = "none"
  )

print(picrust_pca_colorday8)

# Save the plot as a EPS file
ggsave(
  filename = paste0(getwd(),"/Picrust2/results/R/figures/dark/",data_n,"_",daa_method,"_KEGG_qval_0.05_pca_day1_colors.eps"),
  plot = picrust_pca_colorday1,
  device = cairo_ps,
  width = 7,
  height = 4,
  dpi = 600
)
print(picrust_pca_colorday1)  # Save the plot to the file
dev.off()  

# Save the plot as a EPS file
ggsave(
  filename = paste0(getwd(),"/Picrust2/results/R/figures/dark/",data_n,"_",daa_method,"_KEGG_qval_0.05_pca_day8_colors.eps"),
  plot = picrust_pca_colorday8,
  device = cairo_ps,
  width = 7,
  height = 4,
  dpi = 600
)
print(picrust_pca_colorday8)  # Save the plot to the file
dev.off()  
}
```

[5]
Pups/dams comparison all body sites except for rectal (per specific day)

```{r, include=FALSE}
#Define variables
daa_method <- "KEGG"

# Define data set
picrust_set <- list(metadata_All_day1_no_rectal = metadata_All_day1_no_rectal,
                    metadata_All_day8_no_rectal = metadata_All_day8_no_rectal)

# Get dataset names
dataset_names <- names(picrust_set)

# Loop through each dataset
for (i in 1:length(picrust_set)) {
  # Set current dataset and its name
  data <- picrust_set[[i]]
  data_n <- dataset_names[i]
  
# [1]pathway_daa()

## Prepare the files
# convert metadata to data.frame if needed
metadata_mas <- as.data.frame(data) 
# make SampleID row names in the metadata file
rownames(metadata_mas) <- metadata_mas$SampleID
# Keep only the samples in the current metadata
# Use sample IDs that exist in both metadata and kegg_abundance
common_samples <- intersect(metadata_mas$SampleID, colnames(kegg_abundance))
# Subset metadata and abundance to common samples
metadata_mas <- metadata_mas[metadata_mas$SampleID %in% common_samples, , drop = FALSE]
kegg_abundance_sub <- kegg_abundance[, common_samples, drop = FALSE]
# transpose to fit the format of Maaslin2
kegg_abundance_t <- t(kegg_abundance_sub)

# Define output path for this dataset
output_path <- paste0(results_dir, "Picrust_Maaslin2/Picrust_Maaslin2_", data_n)

# Make sure the directory exists
if (!dir.exists(output_path)) {
  dir.create(output_path, recursive = TRUE)
}

maaslin_results <- Maaslin2(
  input_data = kegg_abundance_t, 
  input_metadata = metadata_mas, 
  min_prevalence = 0.1,
  min_abundance = 0.001,
  output = paste0(results_dir, "Picrust_Maaslin2/Picrust_Maaslin2_",data_n),
  fixed_effects = c("TypeHost"),
  reference = c("TypeHost,DamMilk")
)

# Convert results for ggpicrust2
formatted_results <- data.frame(
  feature = maaslin_results$results$feature,
  method = "Maaslin2",
  group1 = "Dam",
  group2 = maaslin_results$results$value,
  p_values = maaslin_results$results$pval,
  q_values = maaslin_results$results$qval,
  coef = maaslin_results$results$coef,
  stderr = maaslin_results$results$stderr
)

# Replace _ with - in pathway name for MetaCyc
formatted_results$feature <- gsub("\\.", "-", formatted_results$feature)
formatted_results$p_adjust <- formatted_results$q_values

# [2] pathway_annotation()
kegg_abun_annotated_kegg_daa_results_df <- pathway_annotation(pathway = "KO",
daa_results_df = formatted_results, ko_to_kegg = T)

# [3] pathway_heatmap()
#filter for only p-value <0.05
kegg_feature_with_p_0.05 <- formatted_results %>% filter(p_adjust < 0.05)

#filter the 'abundance' file based on the values <0.05 and remove the 'pathway' column so that the heatmap function can be run
kegg_abun<-kegg_abundance_sub
kegg_abun$functions <- rownames(kegg_abun)  #add a column to join
kegg_abun2 <- kegg_abun %>%
right_join(kegg_abun_annotated_kegg_daa_results_df %>%
select(all_of(c("feature","pathway_class"))), by = c("functions" = "feature")) %>%
filter(functions %in% kegg_feature_with_p_0.05$feature) 
kegg_abun3 <- kegg_abun2 %>%
    mutate(pathway_class = ifelse(is.na(pathway_class), functions, pathway_class))
kegg_abun4 <- kegg_abun3 %>% select(-"functions")
#for comparisons with >2 groups: remove randomly two rows out of each threesome and then convert the 'description' column to row names because otherwise we can't run the heatmap function 
kegg_abun5 <- kegg_abun4 %>% distinct(pathway_class, .keep_all = T) %>% column_to_rownames("pathway_class")

#export the image
tiff(filename = paste0(getwd(),"/Picrust2/results/R/figures/dark/",data_n,"_", daa_method, "_reference_Day1_qval_0.05_heatmap.tiff"), width=16, height=12, pointsize = 1/300, units='in', res=300)
#create a heatmap
heatmap_plot <- ggpicrust2::pathway_heatmap(abundance = kegg_abun5, metadata_mas, group = "TypeHost", show_row_names = T, font_size = 10)
print(heatmap_plot)
dev.off()

#export the table
write.csv(kegg_feature_with_p_0.05, file = paste0(getwd(),"/Picrust2/results/R/tables/",data_n,"_",daa_method,"_reference_Day1_qval_0.05.csv"))


# [4] pathway_errorbar()

SampleID<-as.data.frame(colnames(kegg_abundance_sub))
metadata2 <- right_join(metadata_mas, SampleID, by=c("SampleID"="colnames(kegg_abundance_sub)"))
# Limit to the Top 30 features
low_p_feature <- formatted_results[order(formatted_results$p_adjust), ]$feature[1:30]

kegg_abun6 <- kegg_abun4 %>% distinct(pathway_class, .keep_all = T) 

# [5] pathway_pca()

metadata3 <- metadata2
metadata3$sample_name <- metadata3$SampleID
# Remove pathways with zero variance across samples
kegg_abundance_filtered <- kegg_abun5[apply(kegg_abun5, 1, function(x) var(x) != 0), ]

# PCoA with more control over colors and shapes
# Define colors
TypeHostNew_colors_pi_day1 <- c(
    "HealthyPupOral"   = "gold1",  
    "DamMilk" = "#66C2A5",  
    "DamVaginal"   = "mediumorchid",  
    "DamOral"   = "gold1"
)
TypeHostNew_colors_pi_day8 <- c(
    "HealthyPupOral"   = "darkgoldenrod4",  
    "DamMilk" = "#117733",  
    "DamVaginal"   = "darkorchid4",  
    "DamOral"   = "darkgoldenrod4"
)

TypeHostNew_shapes_pi <- c(
    "HealthyPupOral"   = 22,  
    "DamMilk" = 15,           
    "DamVaginal" = 15,        
    "DamOral" = 15          
)

# Run PCA manually (example with prcomp, adjust as needed)
pca_result <- prcomp(t(kegg_abundance_filtered), scale. = TRUE)
pca_scores <- as.data.frame(pca_result$x)
pca_scores$TypeHost <- metadata3$TypeHost  # make sure order matches
pca_var <- pca_result$sdev^2
pca_var_perc <- round(100 * pca_var / sum(pca_var), 1)


picrust_pca_colorday1 <- ggplot(pca_scores, aes(x = PC1, y = PC2, color = TypeHost, shape = TypeHost)) +
  geom_point(aes(shape = TypeHost), size = 4, fill = "white", 
             stroke = 2) +   # increase thickness of marker borders) +  
  scale_color_manual(values = TypeHostNew_colors_pi_day1) +
  scale_shape_manual(values = TypeHostNew_shapes_pi) +
  labs(
  x = sprintf("PC1 (%.1f%%)", pca_var_perc[1]),
  y = sprintf("PC2 (%.1f%%)", pca_var_perc[2])
  ) +
  theme_classic() +
  theme(
    legend.position = "none"
  )

print(picrust_pca_colorday1)

picrust_pca_colorday8 <- ggplot(pca_scores, aes(x = PC1, y = PC2, color = TypeHost, shape = TypeHost)) +
  geom_point(aes(shape = TypeHost), size = 4, fill = "white", 
             stroke = 2) +   # increase thickness of marker borders) +  
  scale_color_manual(values = TypeHostNew_colors_pi_day8) +
  scale_shape_manual(values = TypeHostNew_shapes_pi) +
  labs(
  x = sprintf("PC1 (%.1f%%)", pca_var_perc[1]),
  y = sprintf("PC2 (%.1f%%)", pca_var_perc[2])
  ) +
  theme_classic() +
  theme(
    legend.position = "none"
  )

print(picrust_pca_colorday8)

# Save the plot as a EPS file
ggsave(
  filename = paste0(getwd(),"/Picrust2/results/R/figures/dark/",data_n,"_",daa_method,"_KEGG_qval_0.05_pca_day1_colors.eps"),
  plot = picrust_pca_colorday1,
  device = cairo_ps,
  width = 7,
  height = 4,
  dpi = 600
)
print(picrust_pca_colorday1)  # Save the plot to the file
dev.off()  

# Save the plot as a EPS file
ggsave(
  filename = paste0(getwd(),"/Picrust2/results/R/figures/dark/",data_n,"_",daa_method,"_KEGG_qval_0.05_pca_day8_colors.eps"),
  plot = picrust_pca_colorday8,
  device = cairo_ps,
  width = 7,
  height = 4,
  dpi = 600
)
print(picrust_pca_colorday8)  # Save the plot to the file
dev.off()  
}
```

Add Phylum info to tables (manuscript)

```{r}

datasets_fil_scaled_sub <- list(
  ORAL_day1_fil = ORAL_day1_fil,
  ORAL_day8_fil = ORAL_day8_fil,
  RECTAL_day1_fil = RECTAL_day1_fil,
  RECTAL_day8_fil = RECTAL_day8_fil
)

# Get dataset names
dataset_names <- names(datasets_fil_scaled_sub)

# Prepare taxonomy table
general_tax_table <- as.data.frame(tax_table(rar_ALL_fil))
general_tax_table$feature <- general_tax_table$short_title
phylum_table <- general_tax_table %>%
  dplyr::select(Phylum, feature) %>%   distinct(feature, .keep_all = TRUE)  # This removes duplicates


# Loop through each dataset
for (i in seq_along(datasets_fil_scaled_sub)) {
  data <- datasets_fil_scaled_sub[[i]]
  data_n <- dataset_names[i]

# Read the significant reuslts from Maaslin2
maaslin_results <- read.delim(
    paste0(results_dir, "maaslin2/Maaslin2_results_", data_n, "/significant_results.tsv"),
    header = TRUE, sep = "\t", fill = TRUE, quote = "", comment.char = ""
  ) %>%
    mutate(feature = str_replace(feature, "^X\\.", " "),
           feature = str_replace_all(feature, "\\.", " "))


# Join - add Phylum column from taxa table to the maaslin results
joined_results <- maaslin_results %>%
    left_join(phylum_table, by = "feature")

# Write csv of the joined file as a dataframe in the same location
write.csv(
    joined_results,
    file = paste0(results_dir, "maaslin2/Maaslin2_results_", data_n, "/significant_results_with_phylum.csv"),
    row.names = FALSE
  )
}

```

##FEAST
Source-tracking analyses using FEAST

NO-ORAL FEAST ANALYSIS

```{r no-oral FEAST}
source("feast_refactor.R")

feast_no_oral <- run_feast_analysis(
  phyloseq_by_day = list(
    "Day 1" = rar_ALL_day1_fil,
    "Day 8" = rar_ALL_day8_fil
  ),
  analysis_label = "no_oral",
  sink_sites = c("Rectal"),
  source_sites = c("Milk", "Vaginal", "Rectal"),
  batch_correction = FALSE
)

export_feast_analysis(
  analysis = feast_no_oral,
  workbook_path = paste0(results_dir, "tables/FEAST/FEAST_no_oral_results.xlsx"),
  main_plot_path = paste0(results_dir, "plots/FEAST/FEAST_no_oral_pup_level_stacked_bar.png"),
  main_plot_title = "FEAST sensitivity analysis without oral source or sink",
  drop_plot_path = paste0(results_dir, "plots/FEAST/FEAST_no_oral_drop_one_supplementary.png"),
  drop_plot_title = "Supplementary FEAST drop-one sensitivity analysis (no oral)",
  drop_plot_filter_sites = c("Rectal"),
  box_plot_path = paste0(results_dir, "plots/FEAST/FEAST_no_oral_boxplot_main_figure.png"),
  box_plot_title = "FEAST source contributions without oral source or sink"
)
```

DROP-ONE FEAST WITH BATCH CORRECTION - sensitivity analysis

```{r drop-one FEAST}
feast_batch <- run_feast_analysis(
  phyloseq_by_day = list(
    "Day 1" = rar_ALL_day1_fil,
    "Day 8" = rar_ALL_day8_fil
  ),
  analysis_label = "batch_all_sources",
  sink_sites = c("Oral", "Rectal"),
  source_sites = c("Milk", "Vaginal", "Oral", "Rectal"),
  batch_correction = TRUE,
  batch_column_candidates = c("batch")
)

export_feast_analysis(
  analysis = feast_batch,
  workbook_path = paste0(results_dir, "tables/FEAST/FEAST_results_batchcorrect.xlsx"),
  main_plot_path = paste0(results_dir, "plots/FEAST/FEAST_batch_pup_level_stacked_bar.png"),
  main_plot_title = "Batch-corrected FEAST source tracking",
  drop_plot_path = paste0(results_dir, "plots/FEAST/FEAST_batch_drop_one_plot.png"),
  drop_plot_title = "Batch-corrected FEAST drop-one sensitivity analysis",
  drop_plot_filter_sites = c("Oral", "Rectal")
)
```
````
