## Supplementary material for "Barking up the microbiome tree: a multi-body-site characterization of maternal postpartum and pup early colonization dynamics": Barking up the microbiome tree_Supplementary material.docx

for

Running title: “Postpartum Colonization of Dog Microbiome”

^1^Azrieli Faculty of Medicine, Bar-Ilan University, Safed, Israel

^2^Hula Research Center, Tel-Hai, University of Kiryat Shmona in the Galilee, Israel

^3^Koret School of Veterinary Medicine, The Hebrew University Veterinary Teaching Hospital, Hebrew University of Jerusalem, Rehovot, Israel

* These authors contributed equally to this work

**Figure S1.** Functional differentiation of oral (A) and rectal (B) microbiomes between pups by day postpartum in two body sites. Bar plots of the 30 most significant features detected in oral and rectal samples, respectively, show mean relative abundances (± standard error) of key metabolic pathways differing by day postpartum (day 1 – light shades; day 8 – dark shades). The full list of differentially abundant features is available in Table S7.

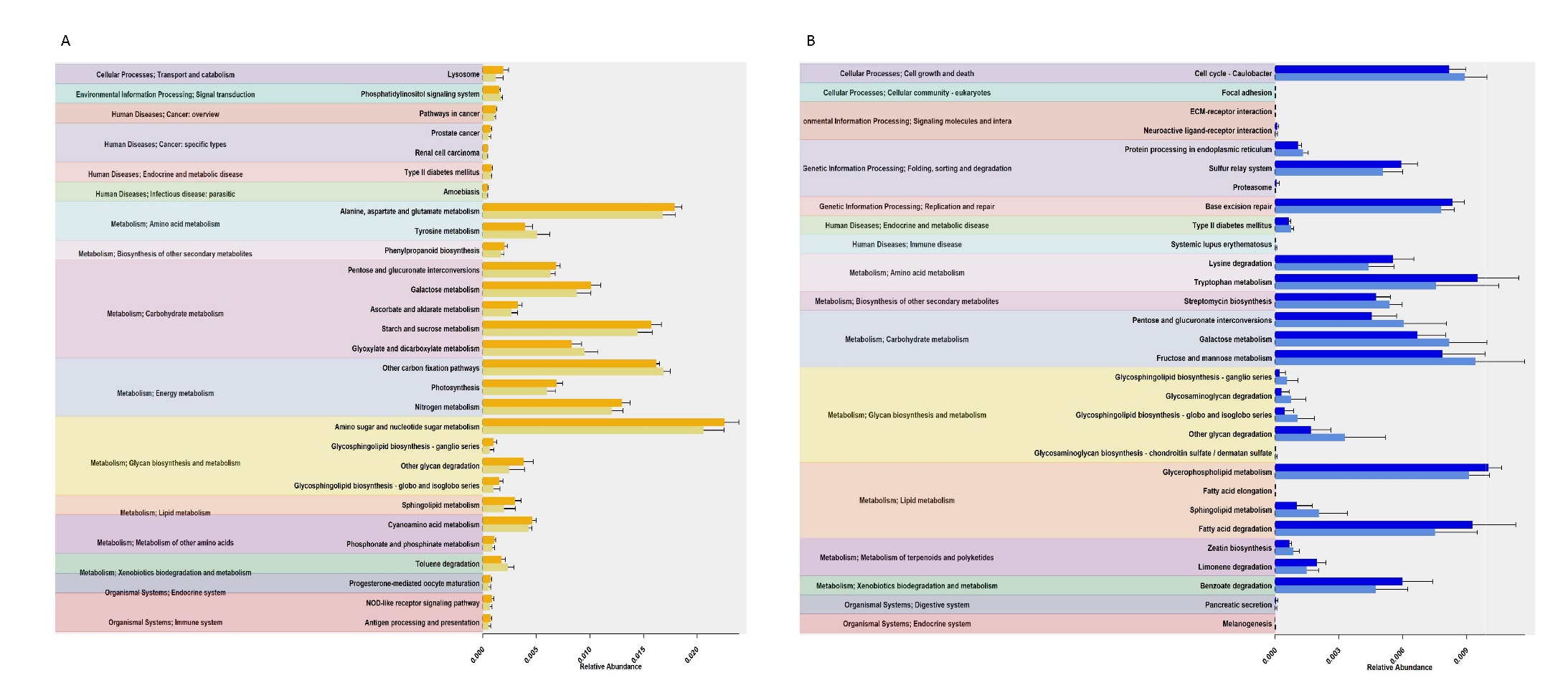
